# Comparative analysis of single-stranded and non-canonical DNA formation in human and other ape cells with telomere-to-telomere genomes

**DOI:** 10.1101/2025.11.03.686349

**Authors:** Jacob Sieg, Huiqing Zeng, Hana Pálová, Linnéa Smeds, Saswat Mohanty, Angelika Lahnsteiner, Francesca Chiaromonte, Kateryna D. Makova

## Abstract

Non-canonical (non-B) DNA secondary structures, e.g., G-quadruplexes and triplex DNA, are mutation hotspots and genome regulators contributing to disease and evolution. Yet they remain uncharacterized in complete genomes *in vivo*. Here we exploited the fact that many non-B DNA structures form single-stranded DNA (ssDNA). Using permanganate/S1 footprinting across 14 cell lines, we generated ssDNA profiles for human and six non-human ape telomere-to-telomere (T2T) genomes. Newly resolved satellite arrays—e.g., at ribosomal DNA and centromeres—displayed high ssDNA levels, implicating non-B DNA in satellite expansion and function. Hidden Markov Models applied to our ssDNA data revealed active genomic domains with specific functions—e.g., replication, transcription, or recombination—each enriched in particular non-B DNA types. Human-specific ssDNA domains correlated with nervous system genes, whereas cancer and embryonic cells showed increased ssDNA in transposable elements. Our ssDNA analysis across ape T2T genomes uncovered conserved and species-specific DNA structural dynamics central to genome regulation.

## Introduction

The telomere-to-telomere (T2T) sequencing and assembly of the human genome^1,2^ represents a milestone in biological research. The human genome has now been deciphered in its entirety, without gaps, paving the way for the comprehensive characterization of its structure, function, variation, and disease associations. Recently, the complete genomes of our closest relatives—the great apes—have also been unraveled.^3,4^ This advance opens opportunities to study human and ape evolution at an unprecedented level of detail.^4^ Together, these two breakthroughs enable the comparative characterization of DNA structures relevant to genome functions—a task that has not been accomplished to date.

As much as 12% of the human genome consists of sequence motifs that can adopt secondary structures distinct from the canonical B-form double helix.^5^ These non-B DNA structures include G-quadruplexes formed by interrupted poly-G tracts (G4 motifs); triplex DNA formed by polypurine or polypyrimidine tracts at mirror repeats (triplex, or TRI, motifs); cruciforms formed by inverted repeats (IR); slipped strand structures formed by direct repeats (DR); Z-DNA formed by alternating purine-pyrimidine repeats (Z motifs); and bent DNA formed by interrupted poly-A tracts (Figure 1A).

**Figure 1.**
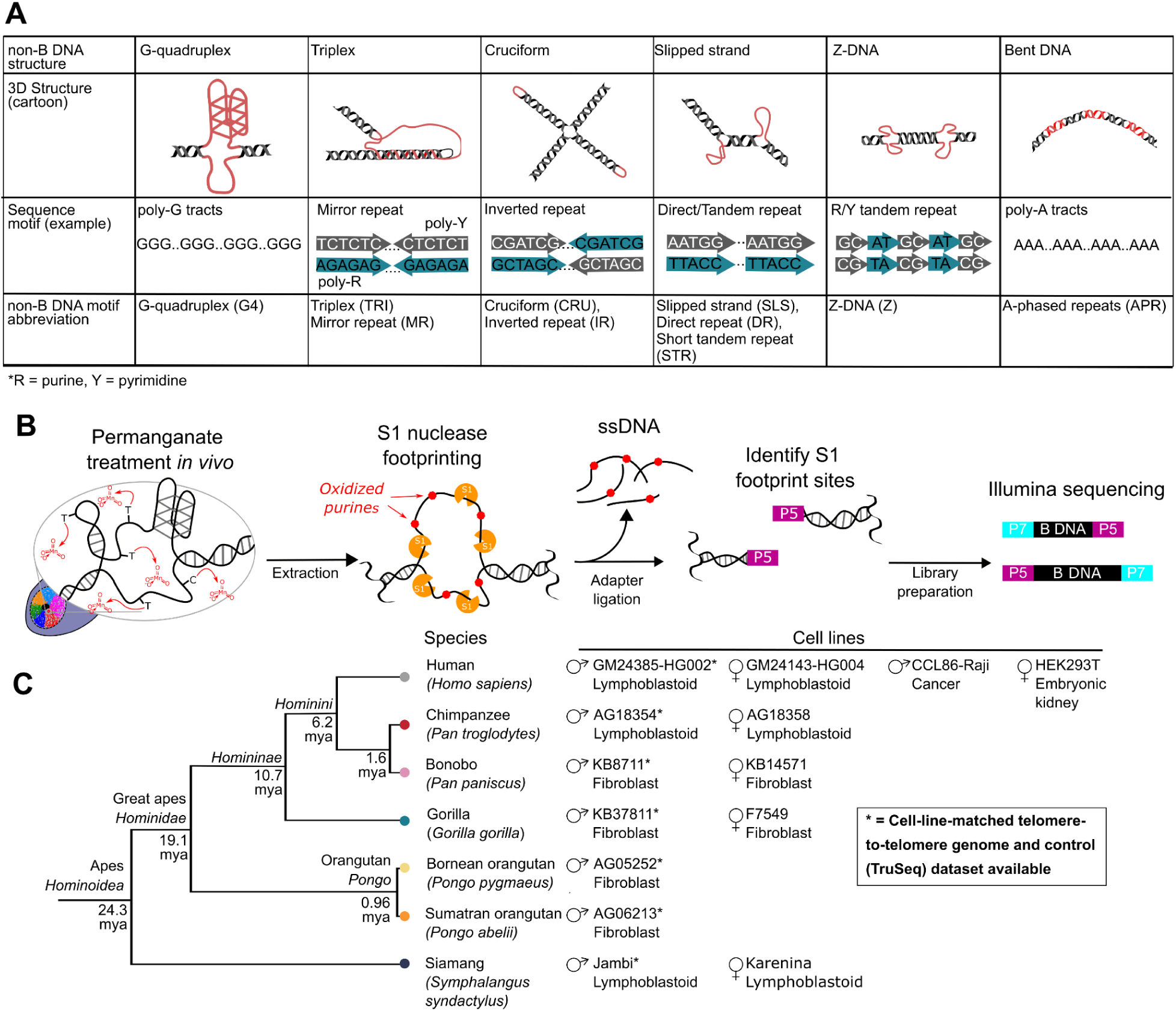
Experimental design. **(A)** Cartoon representation of non-B DNA structures, their associated computationally predicted motifs, and abbreviations used in the text. **(B)** PDAL-Seq experimental workflow where single-stranded DNA (ssDNA) regions are fixed *in vivo* via permanganate oxidation of pyrimidines, ssDNA is footprinted *in vitro* with S1 nuclease, and footprinting breaks are ligated to P5 Illumina adapters before sequencing. **(C)** The phylogenetic relationship and divergence times in million of years ago, mya,(from ^3^) among the studied species and description of the sampled ape cell lines.^4^

Non-B DNA structures are critical to a multitude of functions in normal cells,^6–9^ but they also constitute mutation hotspots^10–14^ and, when dysregulated, are associated with diseases. For instance, G-quadruplexes (G4s) modulate gene expression^15^ by serving as critical components of promoters^15^ and by binding transcription factors.^16–18^ G4s also influence DNA methylation,^19,20^ and remodel chromatin.^21,22^ They contribute to shaping cell-type-specific transcription,^23,24^ and are more abundant in embryonic cells than differentiated cells.^24^ At the same time, G4s mark DNA methylation states associated with diabetes^25^ and cancer,^26^ are overrepresented at oncogenic promoters,^27,28^ are hotspots of rearrangements in cancer genomes,^11,29,30^ and have been suggested to serve as sensors of inflammation during cancer.^31^ Z-DNA facilitates transcription^32^ and is associated with bidirectional promoters,^33^ but its formation also induces genome instability.^34^ Triplex DNA promotes recombination and affects transcription,^35–37^ yet it is unstable and associated with diseases.^38^ As mutation hotspots, non-B DNA structures drive small- and large-scale sequence variation.^13,14,39^ Non-B DNA structures induce polymerase errors,^40,41^ promote recombination,^36,37^ and activate transposable elements.^42,43^ In all, non-B DNA structures appear to represent an evolutionary trade-off, in which their transient formation is required to modulate a multitude of functions, and yet their dysregulation can cause a genetic system’s collapse.

A comparative experimental approach can offer a unique perspective on the conservation of non-B DNA formation and can inform its role in evolution and diseases. Indeed, inter-species comparisons *in vivo* make it possible to separate non-B DNA structures into conserved, i.e., shared among species vs. species-specific. The former are critical to identify because their mutations affecting structure folding can cause genetic diseases. The latter are interesting to detect because they are likely important for adaptation and species-specific traits. In such studies, the experimental determination of non-B DNA structures *in vivo* is critical given their transient nature.

Despite the need for *in vivo* investigations, previous comparative studies of non-B DNA loci were limited mainly to computational predictions.^5,33,44–46^ Importantly, the release of T2T ape genomes^1–4^ provided new genomic sequences that have the potential to form non-B DNA structures.^5^ These non-B DNA motifs comprise ∼9-38% of individual ape chromosomes and are overrepresented at genomic regions unresolved in previous assemblies, including multiple satellite arrays.^4,5^ Annotations in ape T2T genomes^5^ confirmed non-B DNA motif enrichment at functional elements such as promoters and enhancers.^16^ Evolutionary comparisons in great ape T2T genomes revealed conserved G4 motifs at promoters and human-specific G4 expansions associated with neuronal development.^45^ A recent study combined data on G4 structures experimentally validated in human cells with multi-species genome alignments of 241 mammalian genomes, demonstrating that highly-conserved G4 structures are powerful transcription-activating elements.^47^ However, experimental cross-species comparisons of non-B DNA formation have been lacking.

Several antibody-based techniques provided compelling evidence for the *in vivo* formation of specific non-B DNA structures. Immunostaining experiments using structure-specific antibodies demonstrated the formation of G4, triplex, and i-motif structures in cells.^48–51^ Chromatin Immuno-Precipitation with sequencing (ChIP-seq) identified loci that form G4s^24,52^ and Z-DNA.^32^ Recently developed assays (e.g, CUT&RUN and CUT&Tag) have been adapted to detect R-loops, G4s, and i-motifs.^18,53,54^ This progress notwithstanding, questions related to the specificity, sensitivity, and the effects of cell permeabilization on detecting non-B DNA for these techniques remain unanswered.^55^

As an alternative, one can detect non-B DNA structures that form single-stranded DNA (ssDNA) using S1 footprinting techniques.^25,56^ In this approach, ssDNA is digested with nuclease S1, and double-stranded DNA (dsDNA) sites immediately adjacent to the cut locations are sequenced. For example, S1-end-seq has been used to determine triplex DNA formation during replication.^57^ S1 footprinting techniques can be further augmented by permanganate fixation of ssDNA, where it is oxidized with permanganate in cells and identified by sequencing S1-cut sites. Permanganate/S1 footprinting has allowed sensitive detection of diverse non-B DNA loci forming ssDNA, including Z-DNA, triplexes, and G4s.^56^ However, it has never been applied to cells (or cell lines) with completely sequenced genomes. Thus, non-B DNA formation at satellites, suggested to contribute to their function and expansions,^58^ has remained enigmatic.

SsDNA produced by cellular metabolism was also detected by permanganate/S1 footprinting.^56^ For instance, ssDNA is present at replication origins, active promoters, transcription bubbles (where R-loops,^59^ or three-stranded structures consisting of DNA-RNA hybrid and ssDNA, form),^56^ and during the process of recombination.^60^ Uncovering such regions experimentally provides a snapshot of and localizes functional activity across the genome. The permanganate/S1 footprinting technique was previously applied to human and mouse cell lines, identifying ssDNA formation at transcription start sites (TSSs).^61^ Nonetheless, this technique has not been applied to cells with completely sequenced genomes, hindering understanding of genome activity in the previously unaccessible genomic regions.

In this study, we use permanganate/S1 footprinting across 14 cell lines to generate ssDNA profiles in the T2T genomes of humans, chimpanzee, bonobo, gorilla, Bornean orangutan, Sumatran orangutan, and siamang. These ssDNA profiles are analyzed together with non-B DNA motifs and functional annotations from the same genomes (Figure 1A)^5^. Using Hidden Markov Models, we partitioned the ape genomes into states based on levels of ssDNA formation, and studied their associations with genomic functional elements, human-specific loci, cell differentiation, and cancer. As a result, we detected the most active regions of the genome in terms of fundamental molecular processes and studied the interplay among these processes, as well as the role of non-B DNA structures in them. Last, we investigated the formation of non-B DNA structures in newly resolved genomic regions, including satellites.

## Results

### Experimental design

To identify ssDNA forming at several types of non-B DNA structures (Figure 1A) and at functionally active regions in ape genomes, we used <u>P</u>ermanganate/S1 footprinting with <u>D</u>irect <u>A</u>dapter <u>L</u>igation and <u>Seq</u>uencing (PDAL-Seq),^62^ a modification of a previously developed technique.^56^ In PDAL-Seq (Figure 1B), cells are treated with permanganate, which preferentially oxidizes single-stranded pyrimidine bases, breaking the ring structure of the base and oblating base-pairing properties. Thus, ssDNA loci cannot reform B DNA structure during subsequent extraction and library preparation steps, enhancing the sensitivity of PDAL-Seq for non-B DNA detection. Next, DNA is extracted from the cells, and ssDNA is digested with nuclease S1. These S1-cut sites are then ligated to sequencing adapters, purified, and sequenced. The resulting PDAL-Seq reads are thus located immediately adjacent to ssDNA. Therefore, PDAL-Seq is expected to detect ssDNA genome-wide.

We applied PDAL-Seq to 14 cell lines from the seven ape species with T2T genomes, including human, five non-human great apes, and the siamang, a lesser ape (Figure 1C, Table 1). This dataset included seven cell lines with T2T genomes: human, chimpanzee, and siamang male lymphoblastoid cell lines, and bonobo, gorilla, Bornean orangutan, and Sumatran orangutan male fibroblast cell lines. It also included female cell lines from the same species, except for the two orangutans. A human embryonic kidney cell line (HEK293T) and a human Burkitt’s lymphoma cell line (CCL86-Raji, later called “Raji”) were also included to represent embryonic and cancer cell states, respectively. All experiments were performed in at least three technical replicates (see Supplemental Methods). The read mapping statistics (Table S1) were generally well correlated between replicates (except for one replicate that was discarded, Figure S1), allowing us to merge them for subsequent analyses (Table S2). To identify potential artifacts arising from sequencing bias and/or mapping errors, we also analyzed sequencing reads for controls composed of cell-line-matched TruSeq libraries, to which PDAL-Seq was not applied. For both PDAL-Seq and TruSeq libraries, singly mapped reads were reported, where reads either mapped to a single genomic locus or were randomly assigned to one of several equally high-scoring locations (Table S2). The multi-mapping of reads reflects ssDNA formation in ape satellites and other repetitive regions, which were unresolved in pre-T2T genomes (see below; Supplementary Note 1).

**Table 1.** Cell lines used in this study.

| Species | Latin name | ID; aliases | Sex | Cell line type | Source |
| --- | --- | --- | --- | --- | --- |
| Human | *Homo sapiens* | GM24385; HG002 | Male | Lymphoblastoid | Coriell |
| Human | *Homo sapiens* | GM24143; HG004 | Female | Lymphoblastoid | Coriell |
| Human | *Homo sapiens* | Raji; CCL-86 | Male | Burkitt's lymphoma | ATCC |
| Human | *Homo sapiens* | HEK293T/17; CRL-11268 | Female | Embryonic kidney | ATCC |
| Bonobo | *Pan paniscus* | KB8711 | Male | Fibroblast | San Diego Zoological Society |
| Bonobo | *Pan paniscus* | KB14571 | Female | Fibroblast | San Diego Zoological Society |
| Chimpanzee | *Pan troglodytes* | AG18354 | Male | Lymphoblastoid | Coriell |
| Chimpanzee | *Pan troglodytes* | AG18358 | Female | Lymphoblastoid | Coriell |
| Sumatran orangutan* | *Pongo abelii* | AG06213 | Male | Fibroblast | San Diego Zoological Society |
| Bornean orangutan* | *Pongo pygmaeus* | AG05252/3860 (Ken Allen) | Male | Fibroblast | Laura Carrel (PSU), originally from Coriell |
| Gorilla | *Gorilla gorilla* | KB3781 (Jim) | Male | Fibroblast | San Diego Zoological Society |
| Gorilla | *Gorilla gorilla* | F7549 | Female | Fibroblast | San Diego Zoological Society |
| Siamang | *Symphalangus syndactylus* | Jambi | Male | Lymphoblastoid | Lucia Carbone (Oregon Health and Science University) |
| Siamang | *Symphalangus syndactylus* | Karenina | Female | Lymphoblastoid | Lucia Carbone (Oregon Health and Science University) |

### Association between ssDNA structure formation and non-B DNA motifs

We first asked whether ssDNA, as measured by PDAL-Seq reads, was enriched at regions annotated with computationally predicted non-B DNA motifs. We examined non-B DNA motifs previously annotated in ape T2T genomes: G4, TRI, mirror repeats (MR), cruciform repeats (CRU), IR, slipped-strand sequence motifs (SLS), DR, short tandem repeats (STR), Z-DNA, and APR (Figure 1A). Among these, TRI are a subset of MR motifs, SLS and STR are subsets of DR motifs, and CRU are a subset of IR motifs (see Methods).^44^ The subsets are more prone to structural formation than their parent groups.^44^ From these motifs, G4, TRI, IR, CRU, SLS, DR, STR, and Z-DNA motifs are expected to form structures with ssDNA, and thus be detected with PDAL-Seq.^56^

A visual inspection revealed that PDAL-Seq reads exhibited a non-uniform genomic distribution with peaks corresponding to high non-B DNA densities (see Figure 2A for one human cell line and Figure S2 for the other cell lines). This pattern was similar between cell lines from the same species, indicating that ssDNA forms in distinct regions of ape genomes. Regions with especially high PDAL-Seq read density corresponded to areas with high SLS, DR, STR, and/or G4 motif density. These associations were either not observed or inverted (i.e., were negative) in TruSeq datasets (Figure S2), implying that the distribution of PDAL-Seq reads differs from that of conventional sequencing reads.

**Figure 2.**
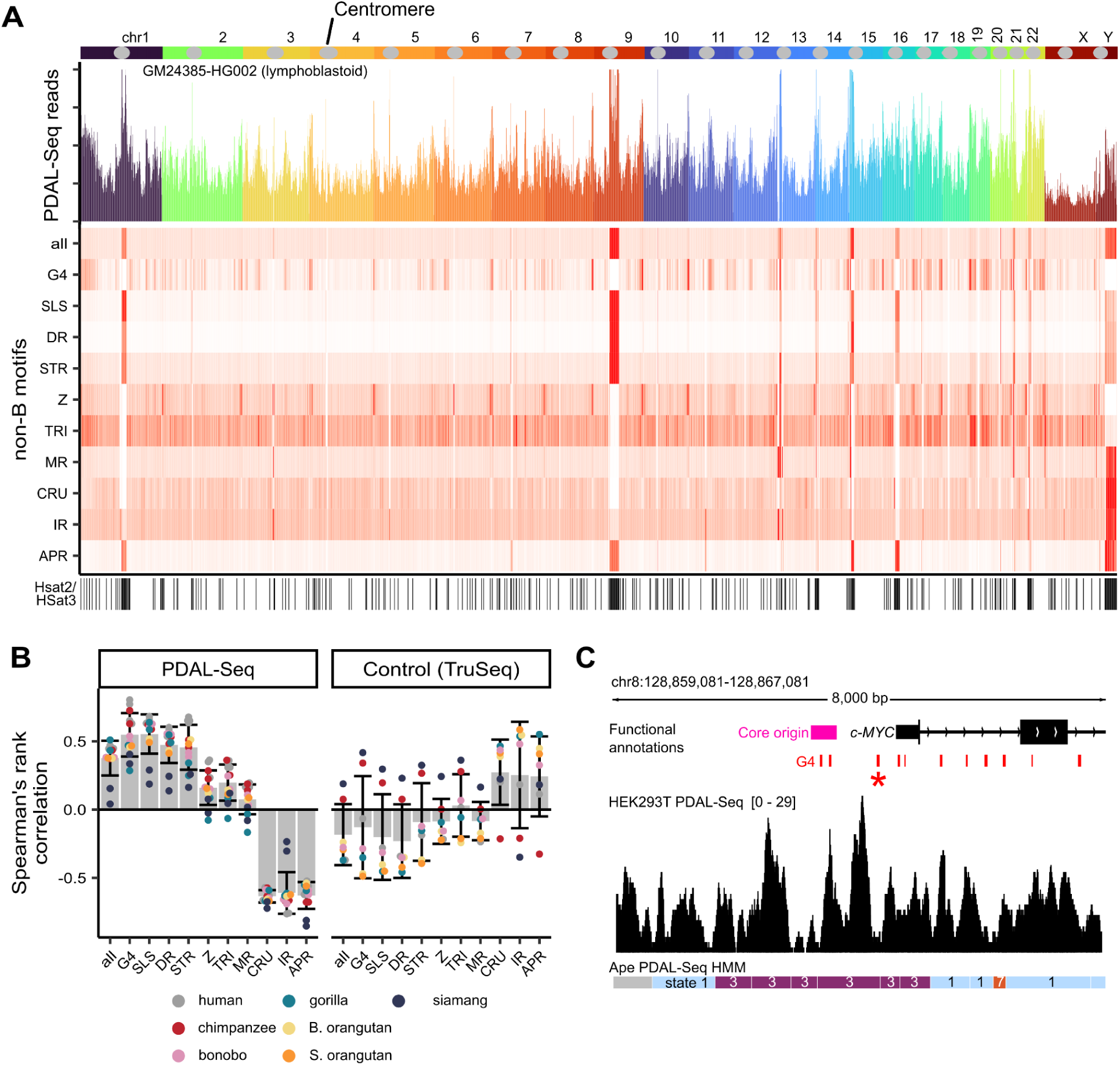
PDAL-Seq signal is correlated with non-B DNA motif annotations across the ape phylogeny. **(A)** Human chromosomes with centromeres marked with grey circles (top), PDAL-Seq reads per Mbp mapped to the human genome, non-B DNA motif coverage in 1-Mbp non-overlapping windows, and human HSat2 and HSat3 satellite array annotations (bottom). Plots share an x-axis which corresponds to chromosomes 1 to 22, chrX, and chrY according to the ideogram colored left to right by chromosome. Complete plots for every species are available in Figure S2. A 99% winsorization transformation was applied to reduce the effect of spurious outliers on non-B DNA motifs and read density values. **(B)** Spearman’s rank correlation coefficients between PDAL-Seq read numbers and non-B DNA motif coverage computed using 1-Mbp non-overlapping windows for each cell line and non-B DNA motif. Bars and error bars represent means and standard error of the means for 14 ape cell lines shown as points. **(C)** Integrated genome viewer trace of the human *MYC* P1 promoter comparing PDAL-Seq data (top), functional annotations (middle), and a hidden Markov model fit to ape-phylogeny wide PDAL-Seq data(bottom). The G4 marked with an asterisk (*) promotes *MYC* transcription.^66^

To quantify an association between ssDNA and non-B DNA at the large scale, we examined Spearman’s rank correlation between the density of PDAL-Seq reads (i.e., the number of reads per window) and the coverage of each non-B DNA motif type (i.e., the proportion of a window annotated as non-B) in 1-Mbp windows (Figure 2B left, Figure S3). PDAL-Seq reads were correlated positively with G4s, especially in human, chimpanzee, and bonobo (rho of 0.49-0.80, Figure 2B). PDAL-Seq reads were also positively correlated with SLS, DR, and STR motifs, especially in great apes (rho of 0.19-0.68, Figure 2B). PDAL-Seq reads were weakly correlated (except for some gorilla and siamang cell lines) with MR, TRI, and Z-DNA motifs (mean rho of 0.14; Figure 2B). Finally, PDAL-Seq reads exhibited a negative correlation with APR, IR, and CRU motifs (mean rho of −0.62; Figure 2B). These correlation patterns were not observed in TruSeq controls (Figure 2B right), where Spearman’s correlation coefficients were usually weaker and opposite in sign compared to the PDAL-Seq datasets (Figure 2B left) for all species except for chimpanzee and siamang. However, for these two species, correlation patterns were still very different between PDAL-Seq and TruSeq datasets (Figures S3B and S3G). Thus, the PDAL-Seq signal showed strong positive correlations with several types of predicted non-B DNA motifs, whereas conventional (i.e., TruSeq) sequencing reads did not. These results were consistent across ape species and indicate that PDAL-Seq is detecting diverse non-B DNA structures in ape genomes.

We also deconvoluted the relative contribution of individual non-B DNA motif types to the PDAL-Seq signal at a smaller scale—in 1-kbp windows—while restricting our analysis to regions with uniquely mapping reads (Table S3). Such an analysis is particularly relevant for non-B DNA structures because their motifs have a median length of 12-50 bp.^5^ After removing multicollinearities arising due to overlaps in motif annotations, we fit Poisson generalized linear models (GLM) using PDAL-Seq read counts as the response and non-B DNA motif densities and GC content as predictors (see Supplemental Note 2). The resulting model explained 48% of variability with minimal multicollinearity (mean McFadden’s R^2^ = 0.48, Table S4; VIFs < 2.1, Table S5). GC content was the strongest positive predictor (Figure S4), consistent with the detection of G4 structures and R-loops in gene-rich regions of the genome. Similarly, TRI, Z-DNA, and DR motifs were positive determinants (Figure S4). G4 motifs were negative determinants in the presence of GC content as a predictor but positive determinants in the absence of GC content (Figure S4). APR and CRU motifs had a strong depleting effect (Figure S4). The positive predictors of PDAL-Seq, especially GC content and G4 motifs, had an opposite, depleting effect on TruSeq reads (Figure S4), consistent with known sequencing biases.^41^ In general, the results from our linear modeling approach at a 1-kbp scale in the uniquely mapping regions of ape genomes support the observations presented for the 1-Mbp scale and indicate that PDAL-Seq identifies various non-canonical DNA conformations.

We examined the promoter of the c*-MYC* proto-oncogene as a locus-specific example (Figure 2C). Transcription of *c-MYC* is dependent on G4 formation in the promoter,^66^ and this region also contains a core origin of replication.^69^ We observed an elevated PDAL-Seq signal in this promoter region in HEK293T, consistent with ssDNA formation. We also observed a PDAL-Seq footprint around a G4 structure known to enhance *c-MYC* transcription.^66^

We next sought to determine whether our PDAL-Seq signal corresponded to antibody-based results for detecting non-B DNA structures in cells. We analyzed several publicly available datasets for the HEK293T cell line, including G4-ChiP-seq, G4-CUT&Tag, and Rloop-CUT&Tag (Figure S5).^53,54,63–66^ The signals in all three groups of datasets displayed strong correlations with the PDAL-Seq signal, with Spearman’s rank correlation coefficients of 0.7 (for two out of two datasets used), 0.66–0.76 (for 13 out of 14 datasets used), and 0.73-0.74 (for six out of six datasets used), respectively (Figure S5A). Overall, a comparison to antibody-based non-B DNA detection datasets indicated that PDAL-Seq detects G4s and R-loops (Figure S5B).

### ssDNA formation is correlated between species and is a determinant of genome function

We next asked how ssDNA formation varies at syntenic locations among ape species. We retained Cactus^67^ multi-species alignment blocks corresponding to established syntenic relationships among great ape chromosomes, while allowing for complex rearrangements between great ape and siamang genomes (see Methods; Table S6). Initially, we considered four species groups separately (Figure 1C): Hominini, Homininae, Hominidae, and Hominoidae; but the results were similar (Table S7, Figure S6). Therefore, below we present our analysis for Hominoidea, which includes all species considered. The final filtered alignment blocks had a median length of ∼500 base pairs, covered 65% of the ape genomes (Table S7), and had PDAL-Seq read unique mapping frequencies >91% in all but one sample (Table S8). To allow comparisons between cell lines, we computed PDAL-Seq read density as reads mapping to each alignment block divided by the block length and dataset size (in bp), and then applied a final *z*-score normalization. We excluded the human HEK293T and Raji cell lines from the cross-species analysis to avoid variance being driven by the distinct ssDNA profiles of embryonic and cancerous cells. We found that PDAL-Seq signals were correlated in syntenic regions between cell lines (Figure S7); phylogenetic effects on ssDNA formation were not apparent, whereas variance was driven by a mixture of cell-line and species-specific effects (Figure S6; Figure S8).

To identify genomic domains with different levels of ssDNA formation, we partitioned alignment blocks using *k*-means clustering and Multivariate Gaussian Hidden Markov Models (MG-HMM) applied to the PDAL-Seq signals (Figure S9). The two approaches produced similar results. Below, we discuss the 8-state MG-HMM for Hominoidea (Figure 3A), chosen because it partitions the alignment blocks into domains of high-to-low PDAL-Seq signal without producing an excessive number of cell-line specific states (Figure S10; see Figure S11 for transition probabilities). For each state, we computed a conservation score (a more similar PDAL-Seq signal will result in a higher conservation score; see Methods) and the enrichment of computationally predicted non-B DNA motifs, genic compartments (promoters, 5’UTRs, exons, introns, and 3’UTRs), other functional genomic elements (enhancers, recombination sites, and core and stochastic replication origins), repetitive elements, and non-genic non-repetitive regions (see Methods). We additionally computed, again for each state, the enrichment of publicly available human RNA-Seq reads against what would be expected from their random distribution across states (see Methods).

**Figure 3.**
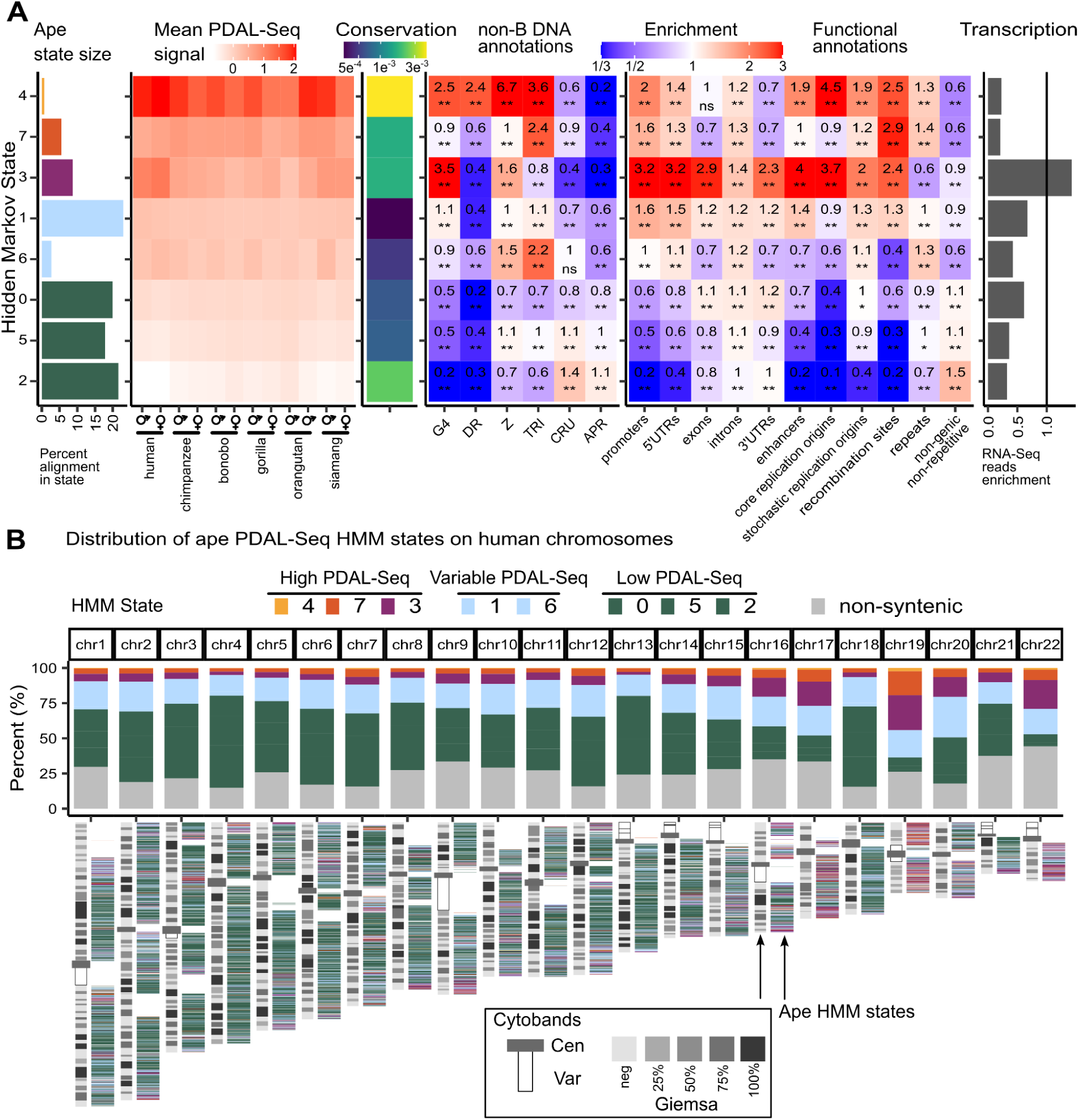
Functionally active genomic domains conserved across ape cell lines are enriched in non-B DNA. **(A)** ssDNA states identified by a Multivariate Gaussian Hidden Markov Model across alignment blocks, characterized through non-B DNA motif predictions and functional genomic element annotations. Percent alignment in state is the base pairs annotated in each state divided by the total alignment size. Mean PDAL-Seq signals are produced by averaging the number of PDAL-Seq reads in each state for each cell line. Conservation scores reflect how similar PDAL-Seq signals are between experiments in each state (see Methods; briefly, conservation scores are inversely proportional to the Euclidean distance of vectors composed of PDAL-Seq signals for each cell line in each state). Enrichments are computed dividing the observed number of annotations overlapping with a state by the number that would be observed by random chance, and are marked as significant (asterisks, * is p ≤ 0.01 nd ** is p ≤ 0.05) or non significant (ns) based on a permutation test (see Methods; briefly, 100 permutations are run to test the null hypothesis that the same intersection would be observed by selecting random genomic loci from the same chromosome with the same annotation lengths). RNA-Seq enrichment is the number of RNA-Seq reads mapping to a state divided by the number that would be observed by random chance. **(B)** Ape states associated with high, variable, and low PDAL-Seq signal mapped onto *Homo sapiens* chromosomes. The top panel is the percentage of each chromosome covered by states with high PDAL-Seq. The bottom panel shows human autosomes with cytoband patterns (left) and ape PDAL-Seq HMM states (right). Cytobands correspond to annotated centromeres (cen), variable regions (var), and Giemsa staining (grey shades), which stain heterochromatin regions.

Three states—4, 7, and 3—were characterized by the highest PDAL-Seq signal across all species and occupied 327 Mb (or 15%) of the human genome present in alignments. These states also showed the strongest association with functional genomic elements (Figure 3A). State 4 had the highest level of ssDNA formation and was the rarest (occupying 15 Mb) and the most conserved state. Compared to average genome-wide levels, it had a 6.7-fold enrichment in Z-DNA motifs, a 3.6-fold enrichment in TRI motifs, a 2.5-fold enrichment in G4 motifs, and a 2.4-fold enrichment in DR motifs. Functionally, it had a 4.5-fold enrichment in core replication origins and a 2.5-fold enrichment in recombination sites. It also had a relatively weak, ∼2-fold enrichment of promoters and enhancers, but these regions were not highly transcribed. State 7 had the second-highest level of ssDNA formation, was more abundant than state 4 (it occupied 121 Mb), and had a less conserved PDAL-Seq signal. It was 2.4-fold enriched in TRI motifs and 2.9-fold enriched in recombination sites. Last, state 3 had the third-highest level of ssDNA formation, with the strongest enrichment in G4 motifs (3.2-fold). It contained the highest densities of genomic regulatory elements, including enhancer, promoter, and 5’-UTR enrichments of 4.0-, 3.2-, and 3.2-fold, respectively. Consistent with this, state 3 had the highest level of gene expression, as measured by RNA-Seq reads. This state was 3.7-fold enriched in core replication origins and 2.4-fold enriched in recombination sites. In summary, the ape PDAL-Seq states with the highest and consistent rates of ssDNA formation across cell lines were associated with genome functions such as replication initiation, recombination, and gene expression.

All three high PDAL-Seq states were spatially related (Figure S11) and were not randomly distributed within the human genome (Figure 3B). Gene-rich human chromosomes 16, 17, 19, 20, and 22 showed the highest proportion of their sequence occupied by high PDAL-Seq states. Likewise, high PDAL-Seq states were not evenly distributed within chromosomes and were localized away from AT-rich regions marked by high rates of Giemsa staining. In summary, high ssDNA-forming states tended to localize in gene- and GC-rich regions of the genome.

States 1 and 6 had intermediate and non-conserved PDAL-Seq signals. Together, they occupied 564 Mbp, or 26%, of the human genome present in alignments. State 1 was the most abundant state, as well as the one with the lowest conservation (Figure 3A). It had a low occurrence of non-B DNA motifs and functional regulatory elements. State 6 also had low conservation but was less abundant. It contained no strong associations with non-B DNA motifs, except for TRI motifs, which were 2.2-fold enriched. Thus, states 1 and 6 were consistent with ssDNA formation in some cell types, but without any strong associations with genomic functional elements. The remaining states—0, 5, and 2—were low PDAL-Seq states, which occupied 1,306 Mbp, or 60%, of the human genome in alignments. They differed only in their relative conservation of low PDAL-Seq signal and depletion of functional genomic elements. Interestingly, state 2, which exhibited the lowest and most conserved level of ssDNA formation, had the highest rate (1.5-fold enrichment) of non-genic non-repetitive sequences, e.g., DNA loci devoid of annotations and considered to have no (known) genomic function.

In summary, our results demonstrate that a substantial portion of the genome forms ssDNA. Moreover, we identified functionally active domains characterized by particular molecular processes and specific non-B DNA types.

### The relationship between ssDNA and methylation, origins of replication, and transcription start sites

We next investigated the relationship between ssDNA and DNA methylation, i.e., the level of 5-methylcytosine (5mC) modification at CpG sites. 5mC levels were computed in each state for the great apes using Oxford Nanopore Technologies long reads (Figure 4A; see Methods). 5mC signals were more consistent among the great ape cell lines for high PDAL-Seq states than for low PDAL-Seq states. State 3, which contains the highest density of G4 motifs and genic regulatory elements, had the lowest rate of CpG methylation (Figure 4A). This result is consistent with an antagonistic relationship between CpG methylation and G4 folding.^19,68^ In contrast, states 4 and 7, which had high frequencies of recombination sites and origins of replication, had the highest level of CpG methylation (Figure 4A). These results indicate that 5mC marks in a region can be compatible with ssDNA formation.

**Figure 4.**
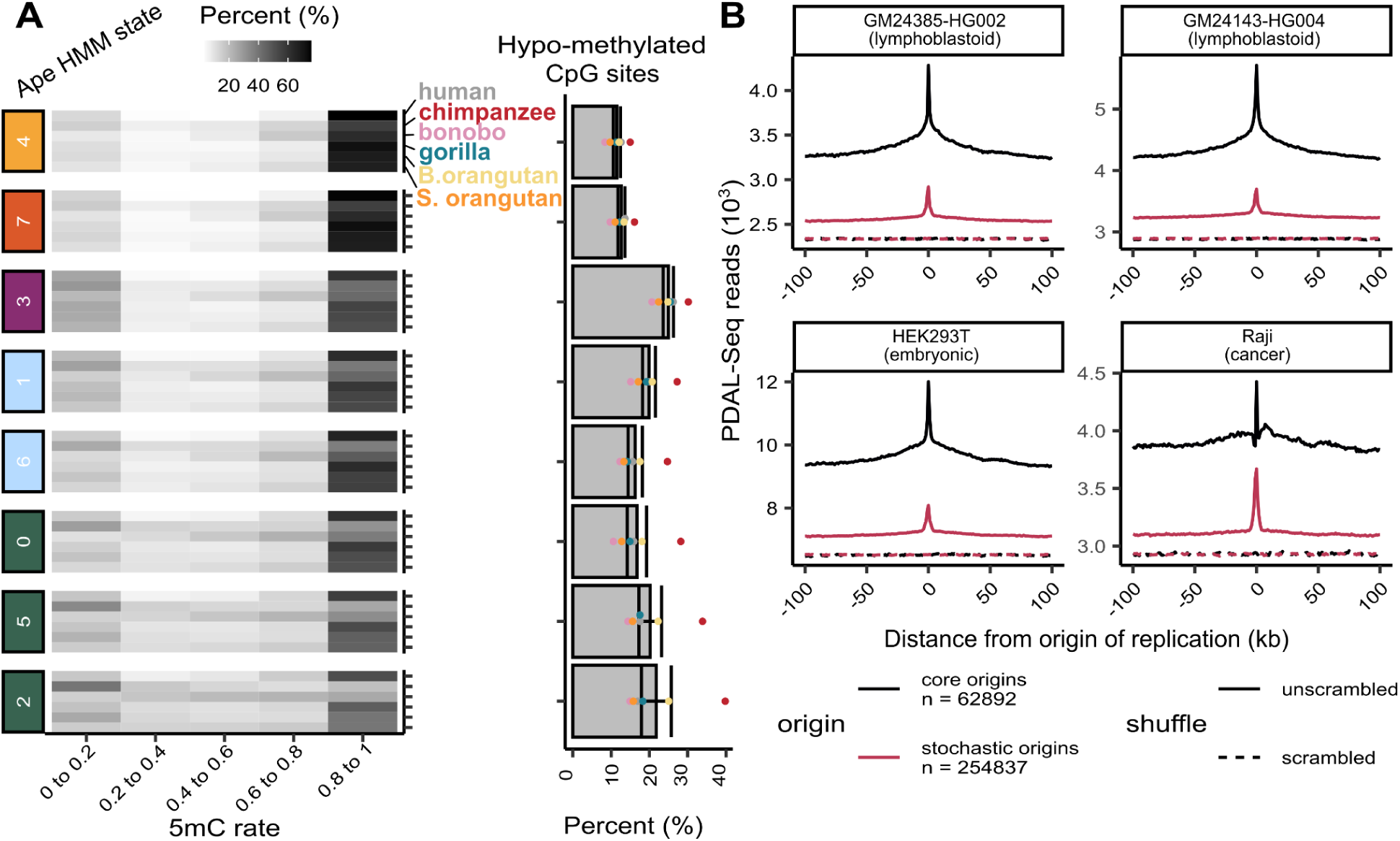
CpG methylation of ape HMM states and ssDNA around origins of replication. **(A)** 5-methyl-cytosine levels at CpG sites called from direct Oxford Nanopore Technology reads. Left: Rows correspond to CpG rates calculated on direct ONT reads for six great ape cell lines with T2T genomes. Fill corresponds to the percent of CpG sites with methylation levels that are within the bins specified on the x-axis. Right: Percent of CpG sites that are hypomethylated (same data as the 0 to 0.2 column in the left panel). Bars and error bars represent means and standard error of the means for six great ape cell lines shown as points. **(B)** Mean PDAL-Seq signals (number of reads) calculated in 1-kbp windows upstream and downstream of the center of core and stochastic replication origin annotations (see Methods). The dashed lines (“scrambled”) show the same quantities computed after assigning origin annotations to random coordinates on the same chromosome.

Because of an interesting link found between the origins of replication and high PDAL-Seq states 4 and 3 (Figure 3A), we explored the relationship between ssDNA formation and core and stochastic replication origins genome-wide (Figure 4B). Core origins are shared among cell types and host ∼80% of all DNA replication initiation events, while stochastic origins are activated only in some cell types.^69^ Analyzing four human cell lines, we found that both core and stochastic origins exhibited elevated PDAL-Seq baselines compared to the genomic average, with a stronger baseline elevation observed for core origins. PDAL-Seq signals tended to peak around both core and stochastic origins, with more marked peaks around stochastic origins for all non-cancer cell lines (Figure 4B). In contrast, the Raji cancer cell line showed a higher average PDAL-Seq baseline around stochastic origins. These results indicate that all replication origins tend to form ssDNA locally, but that core origins are also localized into regions that already have high background ssDNA levels.

Last, we examined the relationship between ssDNA and transcription start sites (TSSs, Figure S12). Average PDAL-Seq signals around ∼20,000 human genes peaked at TSSs for human lymphoblastoid cell lines genome-wide. However, the average PDAL-Seq signal did not peak at TSS for HEK293T and formed a valley at TSS in the Raji cell line. The PDAL-Seq signal exhibited a consistent peak in all four human cell lines at TSSs located within ape MG-HMM state 4 (Figure S12), which corresponds to genomic loci with a high and conserved PDAL-Seq signal in all ape cell lines (Figure 3A). Likewise, the PDAL-Seq signal exhibited a consistent valley in all four human cell lines in TSS located within ape MG-HMM state 2 (Figure S12), which corresponds to genomic loci with low and conserved PDAL-Seq signal (Figure 3A). TSSs located within other MG-HMM states exhibited variation among cell lines. These results indicate that ssDNA formation at TSSs depends on cell type and varies across the genome.

### Biologically significant ssDNA-forming loci in human cell lines illuminated by ape PDAL-Seq states

We next identified ssDNA-forming regions that are likely to be significant in human genome evolution and disease. We postulated that, first, regions with high PDAL-Seq signal in human cell lines, particularly the ones without a syntenic relationship with other apes, would be associated with human-specific genomic features. Second, regions with elevated PDAL-Seq signal for the Raji and HEK293T cell lines would reflect cancer and embryonic cell states, respectively. To identify these regions, we quantified PDAL-Seq reads mapping to the human genome in 1-kbp windows for each of the four human cell lines. Here we considered the entire uniquely mappable portion of human autosomes, including that present in ape multi-species alignments (see Methods). We fit a new MG-HMM to the normalized PDAL-Seq read densities (Figure 5A), choosing a partition in 10 states (Figure S13).

**Figure 5.**
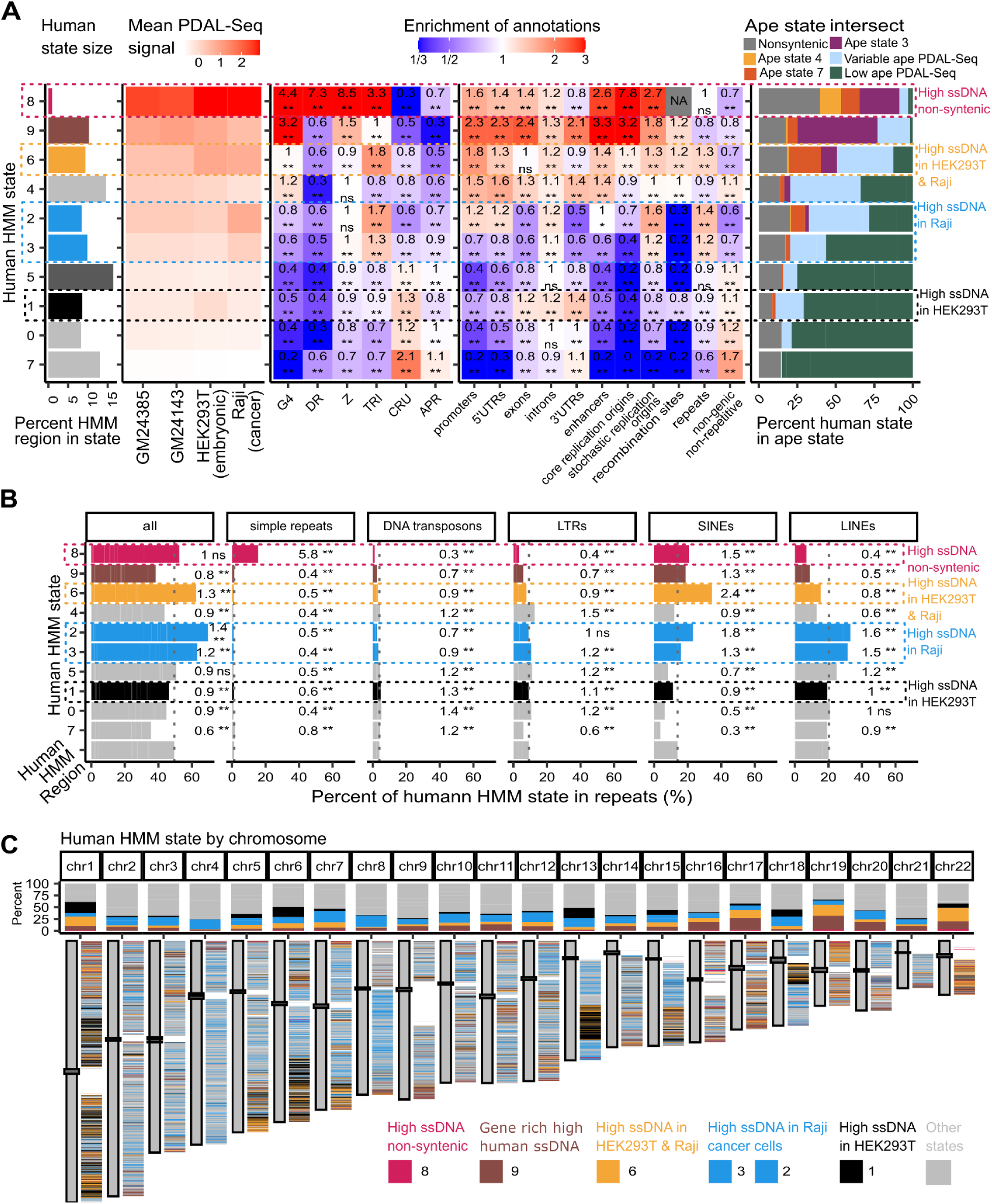
Identification of ssDNA-forming loci associated with human-specific evolution, embryonic cells, and cancer cells. **(A)** ssDNA states identified by a Multivariate Gaussian Hidden Markov Model in the human cell lines, characterized through non-B DNA motif predictions, functional genomic element annotations, and intersections with the ape HMM states. Mean PDAL-Seq signals and enrichments are as described for Figure 3A. For the intersections between human and ape HMM states, percents represent the length of overlap (in base pairs) between each human and ape state, divided by the length of each human state. Non-syntenic regions correspond to human regions absent from ape multi-species alignments. NA (grey) indicates no overlapping annotations. **(B)** Analysis of the repeat content in human HMM states, determined by intersecting them with RepeatMasker annotations. “Human HMM Region” represents the entire region of the human genome subject to segmentation through the HMM fit. Vertical grey-dotted lines mark averages computed across this whole region. States have the same color coding as in panels A and B, and enrichments of repeat annotations and their significance are again computed as described for Figure 3A. **(C)** Top: the percentage of each chromosome covered by the human HMM states, with color coding matching that of panel A. Bottom: human autosomes with annotated centromeres next to annotated human HMM states. In addition to the colors used to represent such states, grey represents all other human HMM states, and white represents regions removed because of low unique mapping rates.

Overall, human and ape MG-HMMs agreed well with each other (Figure 5A, Figure 3A). Both models identified a high PDAL-Seq state associated with G4s and regulatory elements (human state 9, ape state 3). There were intermediate human PDAL-Seq states (e.g., human state 4) corresponding to variable ape PDAL-Seq states. There were also two low human PDAL-Seq states (human states 0 and 7) corresponding to low ape PDAL-Seq states. The human MG-HMM also revealed states not identified by the ape MG-HMM, capturing very high ssDNA formation in all human cell lines, as well as cell-line-specific ssDNA formation (see below).

### Human-specific ssDNA-forming loci are enriched in simple repeats, regulatory loci, and repetitive elements

Our human MG-HMM revealed human-specific states with distinct patterns (Figure 5A). Human state 8, the rarest state (occupying 24 Mbp, or 1% of the human genome analyzed), was ∼40% non-syntenic with the ape states and had a ∼50% intersection with high ape PDAL-Seq states. This state was highly enriched in TRI, Z, and G4 motifs at 3.3-, 7.3-, and 4.4-fold, respectively, and had the highest DR motif enrichment among all human states at 8.5-fold. It was also enriched in enhancers (2.6-fold) and core origins (7.8-fold). In human state 8, the enrichment of non-B DNA motifs, high incidence of enhancers and replication origins, and low rate of synteny were consistent with functionally active human-specific loci.

If human state 8 corresponded to functionally active human-specific loci, we would expect it to have evolutionarily significant functions. To test this hypothesis, we analyzed its gene enrichment^70^ using the Gene Ontology (GO)^71,72^ and the Kyoto Encyclopedia of Genes and Genomes (KEGG) databases.^73,74^ Both analyses resulted in significant enrichments for genes associated with the nervous system, cell signalling, infection, diabetes, and cancer (Table S10). Therefore, this state appears to be enriched in genes with functions particularly important for the human species.

Because human state 8 had the highest enrichment in DR repeats in the human MG-HMM (Figure 5A), we analyzed its repeat content in more detail. We found that it had a 5.8-fold enrichment in simple and low-complexity repeats (Figure 5B; Figure S14) and a 6-fold enrichment in SVA retrotransposons (Figure S14), known to carry predicted non-B DNA motifs.^75,76^ These results suggest that loci in state 8 are often produced by DR expansion of existing non-B DNA-forming motifs and/or SVA insertions into already high ssDNA-forming genomic regions.

### ssDNA formation specific to embryonic and cancer cell lines

The embryonic and cancer cell lines examined in our study exhibited different PDAL-Seq signals in comparison to the human lymphoblastoid cell lines. For example, the Raji cancer cell line clustered separately from all other ape cell lines in the PCA analysis (Figure S15). Additionally, we found a different distribution of PDAL-Seq reads between functional genomic elements and repeats in the Raji cell line and, to a lesser extent, in the HEK293T cell line, in comparison to human lymphoblastoid cell lines (Figure S16). PDAL-Seq reads in the lymphoblastoid cell lines were elevated at promoters (1.3-1.4-fold), enhancers (1.5-1.6-fold), 5’-UTRs (1.2-1.4-fold), exons (1.2-1.3-fold), and core replication origins (2-fold), in comparison to the genome-wide averages. PDAL-Seq reads in HEK293T were less elevated in the same functional elements, with read enrichments of 1.2-fold, 1.4-fold, 1.2-fold, 1.2-fold, and 1.9-fold, respectively. PDAL-Seq reads in Raji were even less elevated and sometimes depleted in the same functional elements, with read enrichments of 1.0-fold, 1.1-fold, 0.8-fold, 0.8-fold, and 1.6-fold, respectively. In contrast, repeats were 1.1-fold, 1.2-fold, and 1.3-fold enriched in PDAL-Seq reads for lymphoblastoid cell lines, HEK293T, and Raji, respectively. These results demonstrate lower levels of ssDNA formation at genic regulatory elements and replication origins, but higher levels of ssDNA formation at repeats, for HEK293T and particularly Raji cell lines, as compared to lymphoblastoid cell lines.

Our MG-HMM models also revealed human states with cell-line-specific patterns for Raji and HEK293T cell lines in comparison to human lymphoblastoid cell lines (Figure 5A). State 6 had higher PDAL-Seq signals in both HEK293T and Raji vs. lymphoblastoid cell lines. States 2 and 3 showed elevated PDAL-Seq signals only in the Raji cell line. State 1 showed elevated PDAL-Seq signals only in HEK293T. In contrast to human states 8 and 9, these states had no strong functional and non-B DNA associations, but instead had modest associations with repeats (1.3-fold, 1.4-fold, and 1.2-fold enrichment for states 6, 2, and 3, respectively). This result again supports ssDNA formation at repeats in HEK293T and Raji cell lines.

We next analyzed the repeat content of human states 6, 2, and 3. All three states were composed of 60% RepeatMasker annotations, which were higher than the 50% repeat content of the entire human genome included in our MG-HMM (Figure 5B). This increase was driven by different repetitive elements for each state. State 6 comprised 35% Short Interspersed Nuclear Elements (SINEs), more than double the 15% SINE content for the entire human genome included in our MG-HMM (Figure 5B), with 2-fold enrichment in *Alu* and SVA elements (Figure S14). For state 2, the increase in repeat content was driven by both Long Interspersed Nuclear Elements (LINEs; ∼30%) and SINEs (∼20%, Figure 5B), with 2-fold enrichment in L1 and *Alu* elements (Figure S14). For state 3, the increase in repeat content was driven by LINEs (∼30%, Figure 5B), particularly L1 elements, which were 1.5-fold enriched (Figure S14). Note that, despite these enrichments, we are likely underestimating ssDNA formation at repeats because we are only considering largely uniquely mapping regions of the genome in human MG-HMMs (see Methods). In summary, compared to lymphoblastoid cell lines, HEK293T and particularly Raji cell lines had elevated ssDNA formation at SINEs and LINEs.

These human, cell-line-specific PDAL-Seq states also exhibited different chromosomal localizations (Figure 5C). State 6, characterized by high PDAL-Seq in both HEK293T and Raji cells, localized with gene-rich regions, especially on chromosomes 1, 17, 19, and 22. In contrast, states 2 and 3, characterized by high PDAL-Seq in Raji cancer cells, were more evenly distributed throughout the genome. Last, state 1, characterized by high PDAL-Seq in HEK293T, localized to large blocks on chromosomes 1, 6, 13, and 18. In summary, ssDNA appears to be activated at different genomic loci in cancer and embryonic cell states in comparison to healthy and differentiated human and ape cell lines.

### non-B DNA structures in ape satellite arrays

Our application of PDAL-Seq to cell lines from species with T2T genomes provides the first opportunity to evaluate ssDNA formation at satellite arrays, which were previously unassembled. Satellites are tandemly repeated arrays of homogenized sequence motifs ranging in length from hundreds of base pairs to several megabases.^77^ Due to their high repetitiveness, satellites were mostly excluded from our ape and human MG-HMMs, which focused on alignable and not-multimapping regions of the human genome, respectively. Below, we specifically studied the enrichment of PDAL-Seq reads at different satellites vs. the rest of the genome, knowing that our results based on short reads will capture genome-wide trends and not locus-specific information. As a control for mapping artifacts, we investigated enrichment for TruSeq sequencing reads (generated without PDAL-Seq) at satellites. Our results suggest that satellites frequently form ssDNA and are enriched in non-B DNA annotations, including but not limited to DR motifs, which are expectedly annotated at many satellites. Below, we highlight our most notable observations.

Multiple centromeric and pericentromeric satellite arrays exhibited significant ssDNA formation (Figure 6A), consistent with the suggested role of non-canonical DNA in centromere definition and evolution.^58,78^ These included active higher-order repeats (HORs) having higher PDAL-Seq read frequencies than inactive or divergent HORs for all species analyzed, in agreement with enrichment of at least one type of non-B DNA in the majority of active centromeres.^5^ Beta satellites in *Pan*, gorilla, and siamang, also showed enrichment in PDAL-Seq reads consistent with their high coverage by G4, Z-DNA, and triplex motif annotations (Figure 6A). We observed PDAL-Seq signal enrichment at the beta-satellite-associated LSAU repeat for most cell lines (Figure S17), consistent with LSAU sequences forming a G4s *in vitro*.^5^ This satellite has variable methylation levels in apes^79^ and was suggested to affect gene expression.^80^

**Figure 6.**
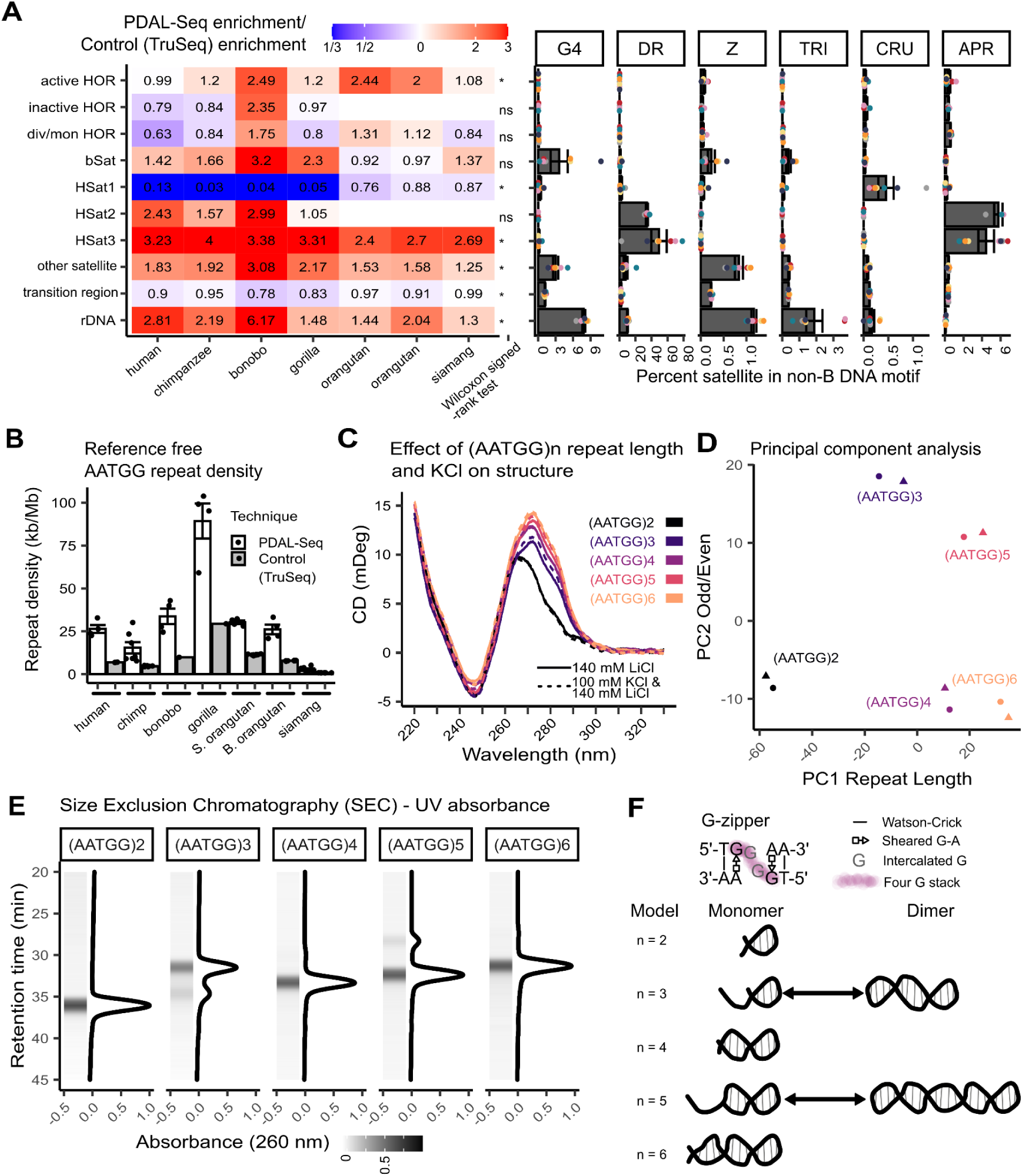
Characterization of ssDNA structure in ape satellites *in vivo* and *in vitro*. **(A)** Analysis of PDAL-Seq reads mapping to ape satellites and non-B DNA motif coverage in ape satellites This includes the seven male cell lines with matched PDAL-Seq and control (TruSeq) datasets. Left: PDAL-Seq read enrichment divided by TruSeq read enrichment (see Methods). For each satellite class in a row, a Wilcoxon signed-rank test is used to contrast the mean enrichments of the seven ape PDAL-Seq datasets paired with seven ape TruSeq datasets (*p* ≤ 0.05 significant differences are marked by asterisks, non-significant ones by ns). Right: amount of each satellite that intersects non-B DNA motifs divided by the satellite length. Points, bars, and error bars represent species, means, and standard error of the means, respectively. Species are color-coded as in Figure 2B. **(B)** Reference-free analysis of HSat3 (AATGG)_n_ tandem repeats in raw PDAL-Seq and TruSeq (control) sequencing reads. The same male cell lines are used as in panel A. Bars and error bars represent means and standard error of the means for technical sequencing replicates (shown as points). **(C)** Effect of KCl and (AATGG)_n_ repeat number on the CD spectrum. **(D)** Principal component analysis of the CD spectrum in panel C. **(E)** Size Exclusion Chromatography (SEC) - Ultraviolet (UV) absorbance analysis of (AATGG)_n_ conformations. The heatmap strip and line plots display the same data. The retention time (y-axis) is reversed for direct comparison to the native gel data in Figure S20. **(F)** Simplified molecular model for G-zipper hairpins.

rDNA arrays present at acrocentric chromosomes displayed an enrichment for PDAL-Seq reads (Figure 6A). This observation is in agreement with rDNA arrays being the most triplex-rich element in the human genome.^81^ Pericentromeric SST1 arrays, frequently present at acrocentric chromosomes, were also enriched in PDAL-Seq reads in all ape species (Figure S17). On acrocentric chromosomes, these arrays have an elevated coverage of G4 motif annotations within their repeated units and of Z-DNA motifs between repeated units, compared to the genome-wide average,^5^ and have recently been shown to drive chromosomal rearrangements leading to Robertsonian translocations.^82^

Interestingly, the newly annotated variable number tandem repeat 148 (VNTR-148)^5^ exhibited 2.3- to 3.6-fold enrichment in PDAL-Seq reads (Figure S17). This satellite is highly abundant in gorilla, bonobo, and chimpanzee genomes, where it occupies 3.8 Mb, 841.9 kb, and 55.9 kb, respectively.^5^ It forms ssDNA most likely because its repeated unit <u>GGG</u>CCA<u>GGG</u>CCCA<u>GGG</u>TTA<u>GGG</u>TT<u>G</u>TTTTA<u>GGG</u>TCA is consistent with G4 folding.

A large number of siamang genomic loci contained high densities of PDAL-Seq reads and IR annotations at subterminal alpha-satellites,^4^ suggesting ssDNA and non-B DNA structure formation in them (Figure S18). This pattern was not observed in the great apes (Figure S3A-F).

Great ape genomes, in contrast, exhibited strong PDAL-Seq signal at loci with large expansions of the pericentromeric HSat3 and, to a lesser extent, HSat2 arrays (Figure 2A, Figure S19). In fact, HSat3 satellite arrays make up the largest portion of satellite content in humans.^77^ Some of these signals were shared across species, whereas others were species-specific (Figure S19). HSat3 arrays are difficult to map unambiguously due to their high repetitive content. We therefore confirmed enrichment of PDAL-Seq signal in them (Figure 6A) using an alignment-free analysis based on counting short tandem repeats in raw sequencing reads (Figure 6B), previously utilized to determine satellite turnover in great apes.^83^ From this analysis, we determined that PDAL-Seq reads contain the HSat3 repeated unit (AATGG) at a higher level than control TruSeq data for human, bonobo, gorilla, and orangutans (Figure 6B), consistent with a large portion of their genomes annotated as this satellite (69 Mb, 63 Mb, 129 Mb, 93-98 Mb, respectively). In comparison to these species, siamang and chimpanzee had a smaller increase in HSat3 repeated units for PDAL-Seq vs. TruSeq experiments, consistent with shorter HSat3 expansions in their genomes (12 Mb and 27 Mb, respectively).^3,83^ These results indicate that HSat3 satellites form non-B DNA structures in ape genomes.

We next sought to characterize the structure of HSat3 sequences *in vitro*. We focused on the purine-rich strand because it exhibits a higher melting temperature than the corresponding duplex^84^ and was the subject of a previous biophysical characterization for (ATGGA)_4_.^85^ We denatured and annealed (AATGG)_4_ under *high* ionic strength conditions in 140 mM LiCl buffer and 100 mM KCl/140 mM LiCl buffer and collected circular dichroism (CD) spectra. The CD spectra were not KCl-sensitive, indicating that the repeats were not forming a G4 (Figure 6C, in grey). Instead, the CD spectra were consistent with the previously characterized G-zipper hairpin structure for (ATGGA)_4_ under *low* ionic strength conditions (50 mM potassium acetate/5 mM magnesium acetate buffer or 25 mM sodium phosphate buffer; Figure 6C).^85^ These results indicate that (AATGG)_4_ also forms a G-zipper hairpin at higher ionic strength and high K^+^ conditions.

To test the effect of repeat length on structure, we denatured and annealed (AATGG)_2_, (AATGG)_3_, (AATGG)_5_, and (AATGG)_6_ in both 140 mM LiCl buffer and 100 mM KCl/140 mM LiCl buffer. We observed the distinct CD signature for the G-zipper hairpin structure at a repeat length of *n* = 3-6 (Figure 6C). CD signatures were also independent of the presence of 100 mM KCl. Indeed, a principal component (PC) analysis of the CD spectra indicated that spectra could be discriminated by repeat length (PC1) and odd vs. even repeat numbers (PC2), but not by buffer conditions (Figure 6D). To further explore these effects, we fractionated samples by molecular weight using size exclusion chromatography with in-line ultraviolet absorbance - multi-angle light scattering - refractive index (SEC-UV-MALS-RI) analysis and native polyacrylamide electrophoresis (Figure 6E; Figure S20). All samples had a peak in the SEC-UV trace (Figure 6E) or a band on the native gel (Figure S20A) that migrated in a manner consistent with a unimolecular weight complex. The oligos with odd repeat numbers—(AATGG)_3_ and (AATGG)_5_—showed a trace or band consistent with high-molecular-weight intermolecular complexes (Figure 6E; Figure S20A). MALS confirmed that the higher molecular weight SEC peak is consistent with a dimer formation (Figure S20B). In summary, our biophysical characterization of (AATGG)_n_ repeats at HSat3 arrays is compatible with a model (Figure 6F) where they form intra-molecular G-zipper hairpins robust to ionic strength conditions and repeat number. Oligos with odd repeat numbers have a higher propensity for forming intramolecular structures than those with even repeat numbers.

## Discussion

Our *in vivo* analysis of ssDNA formation using PDAL-Seq in 14 ape cell lines identified associations between non-canonical DNA and genome function and structure. Such associations were highly localized in ape genomes. Indeed, PDAL-Seq detected a concurrent ssDNA and non-B DNA motif enrichment at functional genomic domains, including those important for replication initiation, recombination, and regulation of gene expression. It also detected ssDNA at satellites, specifically in large HSat3 expansions in great apes, where, according to our biophysical characterization, it is likely caused by the non-canonical G-zipper structure. Last, we used PDAL-Seq signals in multi-species alignments to partition ape genomes by molecular function—pinpointing evolutionarily significant, human-specific non-B DNA loci, as well as a redistribution of ssDNA regions in cancerous and embryonic cell lines.

### non-B structure detection in ape genomes by PDAL-Seq

Compared with other techniques,^52,54^ PDAL-Seq has the following advantages for detecting ssDNA and non-canonical DNA structures. First, the *in vivo* DNA conformations are quickly fixed during the permanganate treatment, which prevents their reannealing, preserves them in their original state, and minimizes the potential for alteration during laboratory manipulations. Subsequent S1 footprinting reaction conditions are optimized to ensure cleavage specificity for ssDNA. Second, PDAL-Seq can detect many non-canonical structures simultaneously. Permanganate preferentially reacts with pyrimidine bases (Cs and Ts), potentially allowing detection of any non-B structure that produces ssDNA, including structures that are yet to be determined.^56,62^ Consistent with this, we observed a strong positive genome-wide correlation between PDAL-Seq signal and G4 motifs (Figure 2B), which can form a well-studied structure with many single-stranded pyrimidines. Some of the highest and most conserved PDAL-Seq regions in ape genomes are enriched in TRI and Z-DNA motifs (Figure 2A). We found moderate positive genome-wide correlations between PDAL-Seq signal and triplex and Z-DNA motifs (Figure 2B); this correlation remained positive for triplex and Z-DNA motifs after accounting for GC content and other motifs. Triplex DNA structure includes single-stranded regions (Figure 1A). In the case of Z-DNA, unpaired bases form at the B-Z junction^86^ and are cleaved by S1.^32^ Thus, PDAL-Seq is likely detecting G4, Z-DNA, and triplex structures in cells. These results are consistent with previous permanganate/S1 footprinting experiments, which detected ssDNA at G4, triplex DNA, and Z-DNA sites.^61^

We observed a negative correlation between the PDAL-Seq signal and bent DNA (APR) as well as CRU motifs (Figure 2B), which form structures not expected to produce large ssDNA regions. Linear modeling indicated that such a negative correlation between PDAL-Seq signal and APR motifs can be explained by their low GC content (Figure S6). APR motifs can contribute but are not equivalent to stress-induced duplex destabilization motifs, which form non-B DNA under certain conditions.^56^ However, IR and CRU motifs, which are predicted to form an ssDNA region in their structures, are negatively correlated with PDAL-Seq reads. CRU motifs maintained their strong depleting effect on PDAL-Seq reads even after GC content was accounted for (Figure S6). It is presently unclear whether the absence of ssDNA signal results from the failure of cruciform structures to form *in vivo* or from PDAL-Seq’s inability to detect them. Cruciforms were also not detected in a previous permanganate/S1 footprinting study.^56^

In some cases, the structure-agnostic detection of non-B DNA structures by PDAL-Seq can make a clear structural interpretation difficult. For instance, PDAL-Seq reads are well correlated with DR motifs genome-wide (Figure 2B), and DR motifs are expected to form slipped-strand non-B DNA conformations, including ssDNA regions. However, it is unlikely that DR motifs are forming only these conformations. For example, 18%, 4%, and 8% of bases annotated as DR motifs are also annotated as TRI, G4, and Z-DNA motifs.^5^ In these cases, slipped-strand structures could be competing with other non-B DNA structures at the same sites. We also determined *in vitro* that HSat3 repeats can form a G-zipper hairpin, a less-studied structure, which is likely also detected by PDAL-seq.^85^

Moreover, PDAL-Seq and other S1-footprinting techniques produce footprints,^62^ not peaks, where different types of structure can produce distinct signal shapes^56^—requiring analysis beyond traditional peak-calling algorithms. We addressed this limitation by analyzing PDAL-Seq reads immediately adjacent to the footprints using Hidden Markov Models, which can capture spatial relationships between genomic loci.^87,88^ Thus, PDAL-Seq is an efficient and structure-agnostic method of detecting ssDNA-forming non-canonical DNA in the genome. As its drawbacks, it detects only structures forming ssDNA and does not differentiate between alternative structures. These limitations could be alleviated by supplementing analysis with structure-specific techniques in the future. As a benchmark, we found that our PDAL-Seq signal was well-correlated with the signals from structure-specific studies that detect G4s and R-loops. Note that PDAL-Seq detects functionally active ssDNA loci genome-wide, even outside previously studied non-B DNA motifs.

### Relationship between ssDNA and genome evolution

The PDAL-Seq data we collected for seven ape species allowed us to determine whether cell line or species effects are more important for ssDNA formation. Our data are consistent with a model in which conserved DNA loci are differentially regulated by cell type at the local scale, but are positioned on chromosomes according to species. First, at the local genomic level, ssDNA formation appears to be conserved among species, with variance driven by cell-type effects. We compared PDAL-Seq data between species in relatively short syntenic blocks (median size of alignment blocks of 500 bp). The PDAL-Seq signal in these syntenic regions was strongly correlated between ape cell lines overall. The variance that existed in the data clustered according to cell type, rather than phylogeny. It is expected that cell-type effects on ssDNA formation dominate in the ∼70% of ape genomes that are amenable to direct sequence comparison between species because of the high conservation of these genomic regions. Thus, a conserved ssDNA-forming locus in a human lymphoblast might have the same structure/function as its syntenic equivalent in every other great ape, but could have a different structure/function in a human/ape fibroblast.

In contrast, when considering the chromosomal scale, the species effect becomes more pronounced. Indeed, the PDAL-Seq signal in 1-Mb windows is nearly superimposable between cell lines of the same species (Figure S2). Even the cancer Raji cell line, which has ssDNA dysregulation at a local level, has nearly identical PDAL-Seq traces to the lymphoblastoid GM24385-HG002 cell line at the chromosomal scale. In part, this pattern can be explained by the fact that some of the highest ssDNA-producing regions in the genome map to satellites, which compose the fastest evolving regions of ape genomes.^4^ We note that comparing PDAL-Seq signals at the chromosomal scale is challenging between species because it is frequently non-syntenic.

### ssDNA and non-B DNA formation determines genome function

The genomic regions that have the highest and most conserved PDAL-Seq signal in all ape cell lines analyzed also have the highest densities of predicted non-B DNA, the strongest enrichment of genic regulatory elements, the highest frequencies of recombination sites, and the highest frequencies of replication origins (ape states 4, 7, and 3, Figure 3A; human states 8 and 9, Figure 5A). Importantly, permanganate/S1 footprinting, the technique from which PDAL-Seq originated, was shown to detect non-B DNA structure formation signals in regions undergoing active transcription, i.e., in transcription bubbles.^56^ Therefore, PDAL-Seq signal detects ssDNA at non-B DNA structures in addition to ssDNA forming due to functional activity of the genome.

We have, for the first time, demonstrated consistent ssDNA formation at core replication origins—the ones that are highly efficient and shared across cell lines.^89^ G4 structures likely form at these regions, as they have been shown to facilitate replication origin firing,^69,90,91^ and origin efficiency is positively correlated with G4 density.^89^ Additionally, we have confirmed and extended previous observations about ssDNA enrichment at transcription start sites.^56^ For differentiated healthy human cell lines, we found ssDNA enrichment at TSSs in all ape states but one (Figure S12). Again, G4s are the likely contributors to this association given their prominent role in many mammalian promoters.^17,21^ Our PDAL-Seq analysis also validated a previously identified association between triplex motifs and recombination.^36,37^ These multi-functional, hyperactive genomic regions are not evenly distributed throughout the genome, but instead occur at higher concentrations on specific chromosomes and at certain genomic locations (Figures 3B and 5C).

Our results are consistent with previous studies showing co-localization between replication origins and genes (reviewed in ^92^; see however^93^). In fact, it has been demonstrated that human replication initiation zones are frequently adjacent to transcribed genes and are enriched in open chromatin and enhancer marks, even when not flanked by genes.^94^ Thus, transcription acts to modulate replication initiation.^94^ Moreover, human activated replication origins are often located at TSSs,^95^ and human constitutive origins (i.e., the most active core origins) are enriched within gene bodies,^89^ providing another explanation for our observations of co-localization of ssDNA between genes and replication origins. Similarly, in humans and mice, replication firing has a distinct pattern in meiosis and contributes to determining recombination patterns across the genome.^96^ We now propose non-B DNA conformations as another likely contributor to these functional co-localizations in the human genome.

This organization of functional genomic regions with high gene expression, high density of replication origins, and/or high density of recombination sites, all accompanied by high levels of non-B-DNA-forming sequences, might have evolutionary advantages given the transient status of non-B DNA formation. The cell cycle requires that, to have any functional impact, alternative structures must outcompete the B-form helix present following DNA replication. Organization of non-B DNA content into distinct chromosomal loci makes it possible for several factors to accumulate, promote, and stabilize such non-canonical DNA formation. These factors include the reorganization/depopulation of nucleosomes,^16^ accumulation of torsional stress to destabilize helices,^97^ active transcription to unfold B-form helixes,^22^ R-loop formation to stabilize alternative DNA structures,^98^ and localization of transcription factors to interact with them.^17^ Additionally, non-B structures are impediments to DNA replication and must be resolved prior or during replication progression.^99–102^ Organization of non-B DNA content into distinct chromosomal domains allows for the accumulation of factors necessary to resolve such alternative structures (reviewed in ^103^), and co-localization with replication origins would provide time for this resolution to occur.

### Human-specific ssDNA-forming loci

We identified regions of the human genome with especially high and consistent levels of ssDNA as evident from PDAL-Seq signals (human state 8; Figure 5A). We found that some of these regions are also enriched in genes that are important for nervous system function and human disease. Non-B structures have been implicated in nervous system development and disease in previous studies. For example, the G4 regulatory landscape changes as stem cells differentiate into neurons.^24^ Additionally, non-B DNA formation has been shown to play a role in several neurodegenerative diseases, and levels of aberrant G4 formation are higher in older patients with such diseases.^104^ Our results now point towards a link between non-B DNA and emergent human loci, which are enriched in genes with nervous system functions. Relatedly, we have recently found that human-specific predicted G4s are enriched in genes from pathways associated with neuronal development and regulation, and are also associated with neurodegenerative diseases.^45^ The intriguing possibility that non-B DNA is involved in the regulation of human-specific traits, including cognition, should be investigated in future studies.

Surprisingly, these high human PDAL-Seq states exhibit ssDNA structures in the cell types used in this study (lymphoblast, embryonic, and cancer), yet show enrichments in genes important for nervous system function and disease. Enhancers are proposed to contain epigenetic information about the current and future developmental potential of the cell, where poised enhancers are inactive but maintain the potential to become active.^105^ A comparative analysis of epigenetic states in primate lymphoblasts identified poised enhancers with gene pathway enrichments for cell-type-unrelated functions (including nervous system function),^106^ consistent with the latent ssDNA structure we observed. One possibility is that these sequences form the same non-B DNA structures in all cell types; however, these structures interact with different regulators, depending on the cell type, leading to differences in gene expression. Another possibility is that non-B DNA structures at these loci differ depending on cell-type-specific stimuli. Future studies should differentiate between these possibilities.

### ssDNA in embryonic and cancer cells

Both the cancerous Raji and embryonic kidney HEK293T cell lines are widely used biomedical models, valued for their fast growth and being amenable to epigenetic manipulation.^107^ Our PDAL-Seq data indicate that the ssDNA structure in these cell lines is redistributed from genic regions to regions of the genome that do not normally produce ssDNA structure in ape cells, particularly transposable elements. This dysregulation is more severe in the Raji than in the HEK293T cell line. SINEs appear to form ssDNA in both cell lines, whereas ssDNA in LINEs is overrepresented mostly in the Raji cell line (Figure 5B). Links between transposable elements and embryonic and cancerous cell states were identified previously. DNA demethylation in primordial germ cells stimulates transcription at transposable elements that are repressed in differentiated cells.^108,109^ Similarly, transposable element reactivation was termed to be “an emerging hallmark of cancers” (reviewed in ^110^). Hypomethylation of TEs in cancer cells is thought to be a contributing factor to reactivation.^111,112^ In terms of function, both cancer-promoting and cancer-suppressive roles of TEs have been shown (reviewed in ^110^), where the former were proposed to mediate cis-regulation of gene activity.^113–117^ We show ssDNA formation in TEs in the cancerous cell line away from gene-rich regions, which could play a role in activating these TEs.

We demonstrate distinct patterns of ssDNA distribution around replication origins and TSSs in the cancerous Raji cell line. Stochastic origins exhibit strong ssDNA enrichment in this cell line only, consistent with increased stochasticity in the use of replication origins accompanying immortalization of cancer cells.^69^ Also, the Raji cell line shows an *underrepresentation* of ssDNA at and adjacent to TSS in seven out of eight ape states in our HMM; in contrast, human lymphoblastoid cell lines show an *overrepresentation* in seven out of eight states. This suggests a shift in transcription regulation between cancer and normal cells. Note that both Raji and HEK293T cell lines were subject to transformation and have well-established genomic abnormalities ^107,118,119^ that could have contributed to the PDAL-Seq signals presented here.

### Non-B DNA and satellites

We detected strong associations between ssDNA structure and several satellite types in the genomes of all apes studied, confirming computational predictions of non-B DNA motif enrichment at these satellites.^5^ These observations would not be possible without T2T genomes,^1,4^ as the loci involved were absent from previous assemblies. For instance, siamangs have large, IR-annotated subterminal alpha satellites^120^ where ssDNA forms at high rates. Subterminal satellites were suggested to be involved in non-homologous recombination between chromosomes in other apes,^121^ and non-B DNA formation in them might facilitate this process.

Great ape PDAL-Seq reads are enriched in HSat2 and HSat3 sequences. Our results indicate that the (AATGG)_n_ HSat3 sequence forms a previously characterized for (AATGG)_4_ G-zipper structure,^85^ and that this structure is robust to changes in repeat number and ionic conditions. The formation of a G-zipper hairpin in HSat3 arrays is consistent with our PDAL-Seq data, as this structure would promote stretches of pyrimidine-rich (CCATT)_n_ ssDNA on the opposite strand, an excellent target for permanganate footprinting.^56,62^ The G-zipper hairpin structure represents an intriguing explanation for HSat3 repeat structure and function. The structure melts well above the physiological temperature range,even in ionic strength conditions below the physiological range, indicating that it is stable.^84,85^ Each G-zipper creates a stack of four guanines that could make a unique interaction site for ligands, proteins, complementary DNA, and/or complementary RNA (proposed by ^85^). The hairpin could form at different lengths. Our results indicate that odd repeat numbers have a higher propensity for intermolecular interactions than even repeat lengths, providing a means for sequence-structure-function specificity.

HSat2 and HSat3 are usually located close to the centromere (except for large, complex HSat3 expansions on the long arm of the Y chromosome).^122,123^ They have been hypothesized to play a structural role in the formation of constitutive chromatin in the nucleus, with the potential to affect gene expression, meiotic recombination, and inter-chromosomal links (reviewed in ^123^). Moreover, HSat2 and HSat3 are transcribed in early embryonic,^124^ senescent,^125^ and cancer cells.^125–127^ The large ssDNA-enriched HSat3 array on chromosome 9 in humans is associated with cellular heat shock^128^ and encodes megabase-scale transcription factor binding sites.^129^ Thus, far from being inert, these satellites appear to have many functions, and a transient non-canonical DNA conformation might be an active regulator of these functions.

We have detected an enrichment in ssDNA at centromeric satellites, including at active higher-order repeats. This observation, together with enrichment in non-B DNA motifs, suggests non-B DNA formation at active centromere arrays. Previous work has shown an enrichment of non-B DNA motifs at ape centromeres.^5,130^ This line of observations is consistent with the recent model proposed by Talbert and Henikoff,^58^ in which non-B DNA structures lead to frequent replication stalling at satellites and break-induced replication drives changes in satellite copy number. Importantly, it was also suggested that this process can provide raw material for centromere drive and eventually to speciation.^58^ This mechanistic working model explaining satellite repeat number expansions and contractions could also operate outside of centromeres,^58^ because non-B DNA motifs and, as we show here, ssDNA are common at ape satellites in general.^5^

### Future directions

PDAL-Seq detected strong associations between ssDNA structure formation, predicted non-B motifs, and genome function in the T2T genomes of apes. Genomic regions that were newly resolved in T2T genomes are enriched in non-B DNA motifs,^1–4^ and our experimental PDAL-Seq data suggest that these motifs are forming non-canonical structures. These structures appear to be folding at highly repetitive regions of the genome. PDAL-Seq is a technique that uses short reads, which have high rates of multimapping in such regions (Table S3).^131^ Therefore, it is frequently impossible to unambiguously assign these reads to an exact location in the genome. Nevertheless, our PDAL-Seq data support the previous computational predictions that newly resolved, repetitive regions of T2T genomes have enormous potential for non-B formation.^1–5,81^

Potential ambiguities linked to short reads could be solved by establishing long-read, direct single-molecule techniques for non-B DNA detection.^132,133^ Such techniques would eliminate normalization steps in data analysis and allow for the determination and comparison of true non-B DNA formation levels between cell lines—and eventually between healthy and diseased tissues. The increased mappability of long reads would also allow researchers to fully leverage telomere-to-telomere genomes. Of particular interest would be assessing non-B DNA formation and its function at individual satellite loci, as they can drive rearrangements between acrocentric chromosomes,^82^ and may contribute to the definition and function of centromeres,^78^ as well as to centromere drive.^58^ Centromeres are among the most variable and dynamic regions in the genome,^134^ and it would be of great interest to evaluate ssDNA and non-B DNA formation at individual centromeres, considering different chromosomes and haplotypes separately.

While not the focus of the present study, the permanganate/S1 footprinting analysis of ssDNA pointed towards the role of non-B DNA in determining nucleosome occupancy.^56^ Namely, the nucleosome occupancies were reduced at non-B DNA-forming ssDNA *in vivo*. This relationship should be studied in more detail using T2T genomes.^56^ Additionally, application of ssDNA and non-B DNA detection methods at the single-cell level would allow one to assess the dynamics of formation of alternative DNA structures at sites with multiple annotations. Future studies should include multiple cancerous cell lines, as well as tumor samples from patients.

## Star Methods

### Data analysis, availability and reproducibility

Raw sequencing reads are available as FASTQ files on the Gene Expression Omnibus (GEO) database at NCBI. Custom analysis was performed using R, Python, awk, and unix shell scripts compiled in a Snakemake pipeline.^135^ This Snakemake pipeline, code for the analysis and to make figures and tables is available at the GitHub link: https://github.com/makovalab-psu/PDAL-seq_submitted. Raw PDAL-Seq reads generated in this study were deposited on the NCBI Sequencing Read Archive BioProject accession numbers **PRJNA1354469** and **PRJNA1354485**. Graphics were generated using R scripts. Minor adjustments to text formatting and panel layouts were performed in Inkscape (V1.3).

### Cell culture

We cultured 14 cell lines (Table 1) prior to applying the PDAL-Seq protocol to them. All cell lines were incubated in a 5% CO2 incubator at 37°C. Passaging and cryopreservation for all the cell lines followed the standard cell culture protocol. Prior to harvesting, cell morphology, viability, and count were confirmed microscopically and using Trypan Blue staining.

Media:

- Human and chimpanzee LCLs and Raji: Roswell Park Memorial Institute Medium (RPMI) Medium 1640 (1×) with L-glutamine (Gibco) supplemented with 15% fetal bovine serum (FBS; Gemini Bio), 1×Penicillin Streptomycin Solution (Pen/Strep; Corning), and freshly added 1× GlutaMAX (Gibco).
- HEK293T: Dulbecco’s Modification of Eagle’s Medium (DMEM; 1×) with glucose, L-glutamine, and sodium pyruvate (Corning) supplemented with 10% FBS, 1× Pen/Strep, and freshly added 1× GlutaMAX.
- Bornean orangutan, gorilla, and bonobo fibroblasts: Minimum Essential Medium (MEM) Alpha (1×) with L-glutamine, ribonucleosides, and deoxyribonucleosides (Gibco) supplemented with 10% FBS, 1× Pen/Strep, and freshly added 1× GlutaMAX.
- Sumatran orangutan fibroblasts: MEM Alpha (1×) with L-glutamine, ribonucleosides, and deoxyribonucleosides supplemented with 15% FBS, 1× nonessential amino acids (NEAA; Gibco), 1× Pen/Strep, and freshly added 1× GlutaMAX.
- Siamang LCLs: RPMI Medium 1640 (1×) with L-glutamine supplemented with 10% FBS, 1× NEAA, 1× sodium pyruvate, 1× Pen/Strep, and freshly added 1× GlutaMAX.

The storage protocol for handling, subculturing procedures, and cryopreservation is listed on the manufacturer’s website for commercially obtained cell lines (ATCC, Corriell). For non-commercial fibroblasts, general fibroblast cell culture principles were followed.^136^ Siamang LCLs were handled and cultured according to the gibbon cell culture protocol established in the Carbone Lab (Oregon Health & Science University). Prior to harvesting, cell morphology, viability, and count were confirmed microscopically and using Trypan Blue staining.

### Permanganate/S1 footprinting with Direct Adapter Ligation and Sequencing (PDAL-Seq)

PDAL-Seq sequencing libraries were prepared as described previously,^137^ with optimizations to improve DNA recovery and sequencing quality. The complete optimized protocol is available in the Supplemental Methods. Briefly, 1-10 million cells were treated with 20 mM or 40 mM potassium permanganate for 80 seconds and quenched/lysed with a buffer containing a molar excess of 700 mM2-mercaptoethanol and 1% sodium dodecyl sulfate. Cell lysate was digested overnight with proteinase K, followed by phenol-chloroform extraction and ethanol precipitation. RNA was digested with RNase A, and DNA was purified by phenol-chloroform extraction and ethanol precipitation. DNA breaks created by physical DNA shearing were blocked with cordycepin and terminal deoxynucleotidyl transferase (TDT). The TDT reaction was further incubated with proteinase K, followed by phenol-chloroform extraction and ethanol precipitation. End-blocked DNA was then treated with S1 nuclease to generate double-stranded DNA breaks at permanganate footprints, followed by phenol-chloroform extraction and ethanol precipitation. S1 cleavage sites were repaired with T4 DNA polymerase to generate blunt DNA ends. This was followed by phenol-chloroform extraction and ethanol precipitation. S1 cleavage sites were then ligated to biotin-ligated P5 Illumina adapters using T4 DNA ligase, and the genomic DNA was purified with magnetic AMPure XP beads to remove unincorporated adapters. Genomic DNA was then fragmented using ultrasonication, and adapter-ligated fragments were bound to magnetic streptavidin beads for purification. Breaks created by sonication for bead-bound fragments were repaired with a NEB Next End Repair enzyme mix (NEB), followed by the ligation to P7 Illumina adapters. Final libraries were amplified with 15 cycles of PCR and sequenced on an Illumina NextSeq 2000 with 150-nt reads for every cell line except HEK293T, which was sequenced using 100-nt reads.

### PDAL-Seq data preprocessing

First, sequencing reads originating from the P5 Illumina sequencing primer were trimmed with `fastp`^138^ with the following settings: default adapter detection, the last base from each read was removed, reads shorter than 15 nt were discarded, a threshold of 30% base complexity was required (defined as percentage of bases that are different from the next base), PolyG and PolyX sequences were trimmed using the Illumina NextSeq/NovaSeq settings with a minimum tail length of 10, and per-read cutting was performed on the head and tail with a sliding window size of 4 and a threshold of Q20. Second, trimmed sequencing reads were mapped to T2T ape genomes^1–4^ using `BWA-MEM` with default settings.^141^ If multiple mappings of a read had the same highest score, we followed the `BWA-MEM`’s default behavior and assigned reads randomly to one of several equally high-scoring locations. Third, reads originating from potential PCR duplicates were removed with `RmDup` (`SAMtools`),^143^ retaining only one copy. BAM files representing technical sequencing replicates that were well correlated (see Mapping Statistics section below) for each cell line were merged using `SAMtools`.^143^

### Controls for sequencing and mapping artifacts

To determine whether signals produced by PDAL-Seq were affected by sequencing and mapping artifacts, we used controls consisting of sequencing data originating from PCR-free Illumina TruSeq libraries for the same cell lines we analyzed with PDAL-Seq. Data were obtained from the Sequencing Read Archive (SRA), and accession numbers are available in Table S1.^3^ Control datasets were preprocessed in the same way we processed PDAL-Seq datasets (described in the previous section).

### Mapping statistics

Mapping statistics are available for different sequencing runs of the same library preparation (Table S1), and for different libraries originating from the same cell line (Table S2). Such technical replicates were merged. Raw and trimmed reads were determined by counting the number of reads before and after trimming with `fastp`.^138^ Primary versus supplementary reads were determined from `samflags` extracted with `samtools view`.^143^ Correlation between technical replicates for each species was assessed in 1-Mb, 100-kb, and 10-kb windows. Genomic windows were generated for each species using `bedtools makewindows`.^144^ The number of PDAL-Seq reads mapping to each window was calculated using `megadepth`.^145^ Spearman’s rank correlation coefficients were determined using the `cor.test` function in base R and plotted in Figure S1.^146^ One technical replicate was removed because it had low correlation with other technical replicates. Uniquely mapped reads were determined by subtracting the MAPQ for each read’s primary mapping location (MAPQ1) from the MAPQ (MAPQ2) for each read’s next best mapping location with a custom AWK script. Uniquely mapped reads were defined as reads where the MAPQ1 - MAPQ2 ≥ 10, which means that the primary mapping location was at least 10 times more probable than the secondary mapping location.

### Genomic element annotations

*Non-B DNA motif annotations* for T2T ape genomes were obtained from Smeds and colleagues.^5^ Briefly, all motifs except G-quadruplexes were annotated with `gfa`^44^ using the following definitions: APRs with minimum three 3-9 bp A-tracts and 10-11 bp between A-tract centers; DRs with units of 10-300 bp and spacer length ≤10 bp; IRs with arms ≥6 bp and spacer length ≤100 bp; MRs with arms ≥10 bp and spacer length ≤100 bp; STRs with units of 1-9 bp and a total length of ≥10 bp; and Z-DNA motifs with alternating purines-pyrimidines of length ≥11. SLS, CRU, and TRI are subsets of DR, IR, and MR motifs, respectively, that are marked by gfa as likely to form slipped-strand, cruciform, or triplex structures, respectively;^44^ these were extracted using grep ‘subset=1’. G4s were annotated using Quadron^43^ with default settings. All output files were converted to BED format for downstream analysis.

*Functional annotations* for ape genomes were obtained from Mohanty and colleagues, see section on “Functional annotations for humans and other great apes”.^45^

*Recombination annotations* were downloaded from NCBI RefSeq GCF_009914755.1, and include regions defined by the International Nucleotide Sequence Database Collaboration as: (1) “meiotic”: a genomic region in which there is an exchange of genetic material as a result of the repair of meiosis-specific double-strand breaks that occur during meiotic prophase; (2) “mitotic”: a genomic region where there is an exchange of genetic material with another genomic region, occurring in somatic cells; (3) “non-allelic homologous”: a genomic region at a non-allelic position where exchange of genetic material happens as a result of homologous recombination; (4) “chromosome breakpoint”: a chromosomal region that may sustain a double-strand break, resulting in a recombination event; (5) “recombination feature”: fits any of the above categories. We combined these annotations for our analysis using bedtools merge^144^ to merge overlapping annotations.

*RepeatMasker annotations* were downloaded from https://github.com/marbl/CHM13 (for human) and https://www.genomeark.org/t2t-all/ for all non-human apes.^4^

*Centromeric satellite* annotations for all species were downloaded from https://www.genomeark.org/.^4^ Active centromeres (annotated as “CEN” or “Cen”) were extracted with grep from the GenomeFeatures tracks downloaded from the same source.

*RNA-seq tracks* were downloaded from https://hgdownload.soe.ucsc.edu/gbdb/hs1/rnas.^122^

### Genome-wide correlation of PDAL-Seq reads and predicted non-B DNA motifs

Non-overlapping 1-Mb genomic windows were generated for each species using `bedtools makewindows`.^144^ The number of PDAL-Seq reads mapping to each window was computed using `megadepth`.^145^ The number of non-B DNA annotations mapping to each window was computed using `bedtools coverage`. Spearman’s rank correlation coefficient (*ρ*) was computed using the `cor.test` function in base R.^146^

### Identification of multimapping regions

Multimapping regions contained genomic regions that produced high rates of multimapping. To identify such regions, first, non-overlapping 10-kb genomic windows were generated for each species using `bedtools makewindows`.^144^ The number of TruSeq control reads mapping to each window was calculated using `megadepth` for BAM files containing all alignments and for a second set of BAM files containing only high-quality alignments (MAPQ > 20). Then, multimapping regions were defined as windows where the number of total alignments was >1.1× the number of high-quality alignments. Unique mapping rates were calculated for regions outside the multimapping regions described above and ranged from 85% to 90%, except for one sample with a rate of 72% (Table S3). Sex chromosomes were excluded from the analysis to eliminate the effects of ploidy.

### Generalized linear modeling of PDAL-Seq read counts on non-B DNA motifs and GC content

To investigate the association between PDAL-Seq and non-B DNA motifs, we regressed PDAL-Seq read counts in uniquely mapping regions of the genome, partitioned into 1-kb windows, on the coverage of such motifs and on GC content. The number of PDAL-Seq reads mapping to a window was calculated using `megadepth`.^145^ GC content was calculated dividing the number of Gs and Cs in each annotation, as determined with `bedtools getfasta`,^144^ by the total length of the annotation. Coverage for a given type of non-B motif in a window was calculated dividing the number of base pairs covered by that type of motif by the total number of base pairs in the window using `bedtools coverage`.^144^ For each cell line, we then fitted a Poisson regression (a Generalized Linear Model) using the `glm` function in R.^146^ In such a regression, the expected PDAL-Seq read count *μ* is expressed as

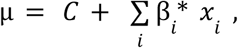

where *C* is an intercept, the *x*_*i*_’s are predictors, and the β_*i*_’s are their regression coefficients. The initial set of predictors considered was

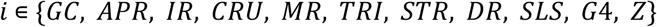

where, in addition to GC content, we included the coverage of ten different non-B DNA motif types. We assessed multicollinearity in this set of predictors by calculating their variance inflation factors (VIFs) in R.^146^ Non-B DNA motif types exhibiting VIF>2 were excluded, as they would duplicate other, correlated types. For example, DR motifs were retained, representing also the highly correlated STR and SLS motifs. Similarly, CRU motifs were retained, representing IR motifs, and TRI motifs were retained, also representing MR motifs. Motifs were chosen to represent IR and CRU motifs; and TRI motifs were chosen to represent MR motifs. Thus, the resulting pruned set of predictors

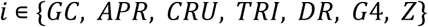

had VIFs at or below 2 for all cell lines. Last, we also fitted Poisson regressions excluding from this set the GC content predictor, to assess its effect on the response and potential changes in the parameter estimates for the other predictors (the non-B DNA types) This further restricted set was

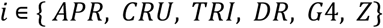

### Alignment preprocessing and PDAL-Seq read assignment

Seven-species CACTUS^67^ alignments of T2T ape genomes were downloaded from https://cgl.gi.ucsc.edu/data/cactus/t2t-apes/8-t2t-apes-2023v2/8-t2t-apes-2023v2.hs1.maf.gz and preprocessed to remove low-quality alignments and enforce syntenic relationships among primate chromosomes.^4,45^

We next processed the alignments following the five steps described below. First, short alignment blocks (<100 bp) and blocks that did not contain all the expected primate species in a species group were removed (for example, Hominini contains humans, chimpanzees, and bonobos, so this step would remove alignment blocks with human and chimpanzee but not bonobo sequences). Second, duplicates from each species in the species group (i.e., blocks containing more than one alignment from each species) were removed, retaining only a single alignment with the best bit score match to the reference alignment. Third, blocks containing alignments with <90% bit score match to the reference were removed. Steps 1-3 were dependent on the species groups used in the analysis (e.g., Hominoidae, Hominidae, Homininae, or Hominini). Fourth, alignments were sorted by coordinates in the CHM13 human reference genome assembly. The fifth step was enforcing synteny. While the syntenic relationship between great ape chromosomes is known,^4^ that between siamang and great ape chromosomes reflects numerous chromosomal rearrangements.^147^ We therefore developed a method that maintained known syntenic relationships in great apes, without prior knowledge, so that alignments from all species could be processed similarly. We reasoned that homologous chromosomes in apes should compose the majority of alignments to the CHM13 reference, with a small population of alignments from other chromosomes due to rearrangements or misalignments. Thus, the chromosome(s) mostly mapping to each chromosome in CHM13 were determined for every ape species. Chromosomes that composed less than 20% of this total were excluded, and these alignment blocks were removed. We confirmed that retained chromosomes reflected the known homology for great ape autosomes and indicated that the retained siamang blocks also reflect homologous sequences (Table S6).^45^ The last step removed blocks corresponding to sex chromosomes in the reference and converted the alignments to one BED-formatted file per species.

Alignment parsing statistics are available in Table S7 and were generated by comparing alignment blocks for each species before and after the BED files underwent a bookended merge using bedtools merge.^144^ The percentage of continuous blocks was calculated as:

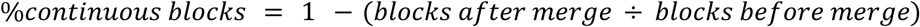

The duplication rate was calculated as:

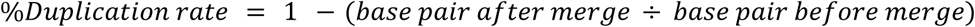

### Read assignment to alignments, normalization, and preliminary analysis

Reads were assigned to alignment blocks using `megadepth`,^145^ and unique mapping was evaluated (Table S8). Unique mapping rates exceeded 91% for all datasets, except for HEK293T, which had a unique mapping rate of >86% (Table S8), likely due to the 100-nt read length used for this cell line. For comparisons between experiments, reads were divided by the length of alignment blocks and by the total number of reads mapping to each alignment, and a final Z-score normalization was applied. This produced a final matrix where rows corresponded to alignment blocks and columns corresponded to the normalized PDAL-Seq signal in each alignment block. Pairwise Spearman’s correlation coefficients were computed for the columns using the `cor.test` function in the base R stats package.^146^ Principal components analysis of PDAL-Seq signal in alignment blocks was performed with the `prcomp` function in the same package. A scree plot of the percentage of variance explained versus principal component was used to determine that two principal components could explain the majority of the variance.

K-means clustering was performed using the `kmeans` function in the base R stats package. For each number of clusters in a range from 2 to 50, we performed 25 random initializations of the centroids and used the best among the corresponding solutions.

### Multivariate Gaussian Hidden-Markov Models

We fitted Multivariate Gaussian Hidden Markov Models (MG-HMM) to the normalized PDAL-Seq data matrix using the `vhmm.VariationalGaussianHMM` function in the `hmmlearn` python library version 0.3.3, with `python 3.9.21`. We used the “full covariance” option, and allowed a maximum of 1,000 iterations. The Bayesian information criterion (BIC) was calculated as follows:

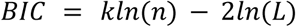

where *k* is the number of parameters estimated in the model (including entries of mean vectors and covariance matrices, as well as prior and transition probabilities), *n* is the number of observations (rows in the data matrix), and *L* is the maximized likelihood function of the model.

A conservation score was calculated to determine how similar PDAL-Seq signals are between experiments for each state. This is inversely proportional to the Euclidean distance of vectors composed of PDAL-Seq signals for each cell line in each state. In more detail: first we calculated the Euclidean distance of the PDAL-Seq signal between experiments for each alignment block, then we computed the mean of such distances across alignment blocks in each state, then we took the inverse of this mean distance, and finally multiplied by the mean PDAL-Seq signal in each state.

### Enrichment of functional data in MG-HMM states

We computed the enrichment for two types of data: (1) functional annotations in discrete genomic windows (BED format; e.g., non-B DNA motifs, functional genomic loci, and repeats), and (2) continuous data (BED graph format; e.g., RNA-seq). Starting with discrete genomics annotations, each annotation type was first subsampled at random to 10,000 annotations. All data were retained if the number of annotations (or of continuous observations) was ≤10,000. Second, the length in base pairs of the intersection between the MG-HMM states segments and the subsampled annotations was recorded using `BEDtools intersect`^144^ and a custom AWK script. Third, the subsample was shuffled to random genomic coordinates – but keeping annotations on the same chromosome. Fourth, the intersection between the MG-HMM states and the shuffled subsampled annotations was computed. This process (subsampling, intersect, shuffle locations, intersect again) was repeated 100 times, and enrichment was calculated by dividing the mean of the first intersect by the mean of the second, shuffled intersect. Significance was determined using a Wilcoxon test in R, if the number of annotations in the original query was >10,000. Alternatively, for annotations with a smaller number of loci, significance was determined by determining how many times the null distribution (shuffling) produced a value that differed from the null mean at least as much as the value computed on the original annotations. The same process was used for continuous data, except that the MG-HMM states segments were subsampled and shuffled, and the sum of the observations in the intersect was used for enrichment instead of the base pairs in the intersect. A similar process was also used for PDAL-Seq read enrichment in human functional annotations; here the annotations were subsampled and shuffled, and the number of reads mapping to the subsample was determined from aligned sequencing reads in the BAM format and `megadepth`.^145^ We used 100 iterations of subsampling because considering a larger number of subsamples resulted in all enrichments being significant due to the large size of the datasets.

### 5-methyl-cytosine rates at CpG sites in MG-HMM states

5-methyl-cytosine modification rates at CpG sites from direct Oxford Nanopore Technologies long reads in the BEDGraph format were intersected with MG-HMM states using `BEDtools intersect` and divided into bins. The percent of CpG sites in each state was reported. These data were originally published in ^4,148^, compiled by ^45^, and downloaded from https://zenodo.org/records/17123210.

### Gene enrichment analysis

Gene enrichments for ape state 4 and human state 8 for pathways in the GO^71,72^ and KEGG^73,74^ databases were performed using `EnrichR (v 0.29.2)`.^70^ Pathways were reported as significant if they produced more than 25 overlapping genes, had a *p*-value from `EnrichR` of ≤0.05, and were not observed by subsampling related MG-HMM states. For ape state 4, pathways were removed from enrichment results if they were observed >25 times in `EnrichR` analysis of 500 subsamples of ape states 3 and 7 (*p* < 0.05). The same process was applied for human state 8 using human state 9 as a null. Thus, we removed gene pathway hits that would be produced simply by subsampling gene-rich MG-HMM states.

### Enrichment of PDAL-Seq reads in CenSat annotations

*Centromeric satellite* (CenSat) annotations for ape species were downloaded from https://www.genomeark.org/.^4,122^ and represent the most extensive and accurate curation of ape genomic satellite arrays currently published. Centromeric satellite files were processed into previously published categories^122^ using a custom R script to parse the CenSat file for the human CHM13 genome by recognizing the following text patterns in the annotation names (Figure 6A):

- “active HOR” satellites with the following names: “S1C1/5/19H1L”, “S2C2H1L”, “S01/1C3H1L”, “S2C4H1L”, “S01C6H1L”, “S1C7H1L”, “S2C8H1L”, “S2C9H1L”, “S1C10H1L”, “S3C11H1L”, “S1C12H1L”, “S2C13/21H1L”, “S2C14/22H1L”, “S2C15H1L”, “S1C16H1L”, “S3C17H1L”, “S2C18H1L”, “S2C20H1L”, “S3CXH1L”, and “S4CYH1L”.
- “inactive HOR” satellites specified with the “hor” classification but not in the “human active HOR” category mentioned above.
- “div/mon HOR” satellites specified with the “dhor” and “mon” classification.
- “HSat1” satellites specified with the “hsat1A” and “hsat1B” classification.
- “HSat2” satellites specified with the “hsat2” classification.
- “HSat3” satellites specified with the “hsat3” classification.
- “bSat satellites specified with the “bsat” classification.
- “other satellite” satellites specified with the “censat” and “gsat” classifications.
- “rDNA” satellites specified with the “rDNA” classification.
- “transition region” satellites specified with the “ct” classification.

The same categories were generated for the ape T2T genomes^4^ using the following text patterns:

- “active HOR” satellites specified with the “active_hor”
- “inactive HOR” satellites specified with the “hor”
- “div/mon HOR” satellites specified with the “mon”, “mon/hor”, and “dhor” classifications.
- “HSat1” satellites specified with the “HSat1B” and “HSat1A” classifications.
- “HSat2” satellites specified with the “HSat2”
- “HSat3” satellites specified with the “HSat3”
- “bSat” satellites specified with the “bSat”
- “other satellite” satellites specified with the “gSat” and “cenSat” classifications.
- “rDNA” satellites specified with the “rDNA”
- “transition region” satellites specified with the “ct”
- Categories that were not included in the analysis were satellites specified with the “mixedSuperFamily”, “subTerm”, “GAP”, “GAP,HSat3”, “HSat2_3”, “subTerm_mon”, and “subTerm_mon/hor” classifications.
- *S. syndactylus* subterminal alpha-satellite arrays were defined as centromeric satellite annotations corresponding to “subTerm_mon/hor(SF4)” arrays and extracted with `grep`.

Read enrichment (Figure 6A) in each category was calculated by fragmenting parsed annotations into 1-kp windows. These 1-kb windows were randomly subsampled to 1,000 windows. All data were retained if the number of annotations or of continuous observations was ≤1,000. The total number of PDAL-Seq reads mapping to the subsampled annotation was recorded using `megadepth`^145^ and a custom AWK script. Next, the subsample was shuffled to random genomic coordinates while keeping annotations on the same chromosome. The total number of PDAL-Seq reads mapping to the subsampled and shuffled annotation was recorded. This process (subsample, compute reads, shuffle, and compute reads) was repeated 100 times, and enrichment was computed by dividing the mean of the unshuffled intersect by the mean of the shuffled intersect. Significance was determined using a Wilcoxon test in base R, if the number of windows in the original query was greater than 1,000. Alternatively, for CenSat annotations with a smaller number loci, significance was determined by determining how many times the null distribution produced a value that differed from the mean of the null at least as much as in the original annotation.

The newly annotated variable number tandem repeat 148 (VNTR-148)^5^ was extracted from CenSat annotations using grep to recognize the pattern “cenSat/VNTR_148”. Some analysis used annotations from Repeat Masker directly. HSat2 and HSat3 arrays were defined as RepeatMasker annotations corresponding to all circular permutations of “HSATII” and to “(ATTCC)_n_” satellite arrays, respectively, and extracted with grep (Figure S2). SST1, SATR1, SATR2, and LSAU categories were extracted with grep and read enrichment was calculated as described above (Figure S17).

### Reference-free short tandem repeat analysis

Short tandem repeat content was quantified as described previously.^83^ Trimmed reads were converted from the FASTQ to FASTA format. Then tandem repeats were determined using the Tandem Repeats Finder program (TRF, `trf409.legacylinux64 {input} 2 7 7 80 10 50 2000 -l 6 -f -d -h -ngs`).^149^ Raw TRF outputs were parsed using `parseTRFngsKeepHeader.py` from https://github.com/makovalab-psu/heterochromatin to identify reverse complements and cyclic permutations of the same sequence. Results were further parsed into a tabular format and reported as thousand base pairs (kb) of tandem repeat detected per million base pairs sequenced (Mb).

### Biophysical characterization of (AATGG)_n_ repeats

#### DNA synthesis, purification, and folding

DNA for biophysical studies was ordered from Integrated DNA Technologies (IDT) with a standard desalting purification. Oligos were suspended in nuclease-free water to 100 µM; the concentration was determined using the absorbance at 260 nm and extinction coefficients from IDT ((AATGG)_2_ = 0.1087 µM^-1^cm^-1^, (AATGG)_3_ = 0.1622 µM^-1^cm^-1^, (AATGG)_4_ = 0.2157 µM^-1^cm^-1^, (AATGG)_5_ = 0.2692 µM^-1^cm^-1^, and (AATGG)_6_ = 0.3227 µM^-1^cm^-1^). Purity was confirmed with high-pressure liquid chromatography. Briefly, sample fractions were monitored with the absorbance at 260 nm and 280 nm using a Waters AQUINITY Arc UPLC system, equipped with a Waters Xbridge C18 2.3 μm 4.6×150 mm column, a denaturing 60 ℃ column temperature, and a two-component reverse-phase-ion-pair chromatography system. Buffer 1 was 0.1 M triethylamonium acetate and buffer 2 was 20:80 acetonitrile:0.1 M triethylamonium acetate. The buffer gradient was 5 min in 100% buffer 1, 5-30 min transition to 100% buffer 2, 30-35 min in 100% buffer 2, and 35-40 min transition back to 100% buffer 1. Purity was determined by the presence of one peak in the HPLC chromatogram. DNA was purified using the same HPLC method and speed-vacuumed to remove the HPLC solvent if required. DNA was then prepared in either 140 mM LiCl, 20 mM LiMOPS pH 7.2 buffer or 100 mM KCl, 140 mM LiCl, 20 mM LiMOPS pH 7.2 buffer to a final 80 µM AATGG monomer concentration. The 80 µM AATGG monomer concentration was used to maintain consistent signal between repeat lengths (i.e., 40 µM (AATGG)_2_, 26.7 µM (AATGG)_3_, 20 µM (AATGG)_4_, 16 µM (AATGG)_5_, and 13.3 µM (AATGG)_6_). Samples were denatured and slowly annealed using a PCR thermocycler (1.5 min at 95℃, 10 min at 90℃, 10 min at 80℃, 10 min at 70℃, 10 min at 60℃, 10 min at 50℃, 10 min at 40℃, 10 min at 30℃, 10 min at 20℃, and 10 min at 10℃) to ensure thermodynamic equilibrium, and were stored at 4℃.

#### Circular dichroism (CD) spectroscopy

CD spectra were collected in a 0.1 cm quartz cuvette at 20℃ on a Jasco J-1500 spectrometer with the following settings: photometric mode-CD; HT. Abs, CD scale: 200 mdeg/1.0 d0D; FL scale: 200 mdeg/1.0 d0D; DIT: 4 sec; bandwidth: 1 nm; scanning speed: 50 nm/min; baseline correction: No Correction; accumulation: 3 times. The sample chamber and lamp were continuously purged with dry nitrogen to prevent condensation of ambient water. Buffer CD was subtracted at each wavelength. Molar ellipticity (*e*) was calculated using the following equation:

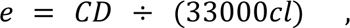

where *CD* is the circular dichroism in mDeg, *c* is the precise molar concentration, and *I* is the path length in cm. The precise molar concentration *c* was determined with the A260 of the sample at 95℃ in sealed 0.1 cm cuvettes.

#### Native gel electrophoresis

67 pmol of AATGG monomer was fractionated on a 15% 19:1 acrylamide:bisacrylamine gel containing 10 mM KCl and 1×TBE buffer (this corresponds to 34 pmol of (AATGG)_2_, 22 pmol of (AATGG)_3_, and so on). Samples were prepared by diluting 1 µL CD sample in 19 µL CD buffer and weighing samples with 4 µL of 60% sucrose and 0.025% bromophenol blue loading solution. Gels were preheated for 30 min at 150 V in a 4℃ cold room prior to loading. 20 µL of the sample was fractionated at 150 V in a 4℃ cold room for ∼6 hours. A NEB Low molecular weight ladder (NEB N3233) was included as a size comparison. The gel was stained in 1×SYBR gold working reagent in water for 15 min and imaged on a Bio-Rad GelDocGo imaging system.

*Size exclusion chromatography - UV absorbance - Multi-angle light scattering - Refractive Index analysis (SEC-UV-MALS-RI).* Samples were fractionated on an Agilent liquid chromatography system with an in-line Agilent UV detector, Wyatt DAWN Multi-angle light scattering detector, and Wyatt Optilab Differential refractive index (RI) detector. A SEC Analytical Column (size: 5µM, 100Å, 7.8mm) was equilibrated overnight with 140 mM LiCl and 20 mM LiMOPS (pH 7.2) buffer at 4°C at a flow rate of 0.5 mL/min. Then, 50 µL of prefolded 80 µM AATGG monomer was injected onto the column and monitored for ∼1 hour. Molecular weights in MALS peaks were calculated using Wyatt Optilab analysis software with both UV and RI as concentration sources.

## Acknowledgments

We are grateful to Oliver Ryder and Cynthia Steiner (San Diego Zoological Society), Lucia Carbone, Laura Carrell, Coriell, and ATCC for providing cell lines, and to members of the Carbone lab for sharing their protocol to culture siamang LCLs and for troubleshooting its use in our lab. We thank Sarah Craig and Kaivan Kamali for training us in the PDAL-Seq experimental technique and data analysis. We thank Kristin Eckert for invaluable feedback on the initial draft of this paper and to Harmit Malik for discussing our results. The co-authors would like to acknowledge the Penn State University Huck Institutes’ Genomics Core Facility (RRID:SCR_023645) staff for collecting the PDAL-seq data on the Illumina NextSeq 2000; the Huck Institutes X-Ray Crystallography and Scattering Core Facility (RRID:SCR_024464) for the use of the Wyatt SEC-MALS; Julia Fecko for collecting the data for Figure 6E and Figure S20B; and the Huck Institutes’ Biomolecular Interactions Core Facility (RRID:SCR_024458) for use of the Jasco CD instrument. This work was supported by a National Institutes of Health National Institute of General Medical Sciences R35 grant number GM151945 and by the Willaman Endowment Fund to KDM.

