## Supplemental notes, methods, and Figures for "Comparative analysis of single-stranded and non-canonical DNA formation in human and other ape cells with telomere-to-telomere genomes"

### Table of contents


**Table of contents** 1

**Supplemental Notes** 3

Note 1. PDAL-Seq data preprocessing and TruSeq control datasets 3

Note 2. Biophysical characterization of Hsat2,3 repeats 3

**Supplemental methods** 4

Optimized PDAL-Seq protocol 5

Reagent preparation, storage, and handling 5

Day 1: Harvest and treatment 6

Day 2: DNA extraction 6

Day 3: RNase treatment 7

Day 4: Block 3'-DNA ends 7

Day 5: S1 Nuclease digestion trial (find the optimal concentration) 7

Day 6: Large scale S1 Nuclease digestion 8

Day 7: End Repair with T4 DNA Polymerase 8

Day 8: P5 adapter ligation 9

Day 9: Bead size-selection, sonification, streptavidin pull-down, end repair, P7 ligation 9

Day 10: PCR 10

**Supplemental figures** 12

Figure S1. Spearman's rank correlation coefficients between PDAL-Seq technical sequencing replicates in differently sized genomic windows 12

Figure S2 Comparison of PDAL-Seq read density with non-B DNA motif density for each cell line 13

Figure S3. PDAL-Seq read density versus non-B DNA motif density for each cell line, separated by non-B DNA motif 16

Figure S4. Poisson generalized linear modeling (GLM) to identify the relationship between ssDNA versus non-B DNA motifs in 1-kbp windows corresponding to uniquely mapping regions of the genome 19

Figure S5. Comparison of PDAL-Seq to 2 published G4 ChiP-Seq, 14 G4 CUT&Tag, and 6 R-loop CUT&Tag datasets in the HEK293T cell line. 20

Figure S6. Principal component analysis of PDAL-Seq reads in alignment blocks 22

Figure S7. PDAL-Seq signal in syntenic regions of the genome is correlated between ape species cell lines 23

Figure S8. Pairwise Spearman's rank correlation coefficients from Figure S8 for different types of comparisons 24

Figure S9. Scree plots to determine state numbers for the analysis of PDAL-Seq data in ape alignments 25

Figure S10. Partitioning of variance as a function of hidden state number for Multivariate Gaussian Hidden Markov Model for the Hominoidae species group 26

Figure S11. Transition probability matrix for the 8-state Multivariate Gaussian Hidden Markov Model presented in Figure 3 27

Figure S12. Mean PDAL-Seq signal upstream and downstream of transcription start sites (TSSs) 28

Figure S13. Scree plots to determine state numbers for the analysis of PDAL-Seq data in the human genome 28

Figure S14. Enrichment of repeat families in human PDAL-Seq MV-HMM models 30

Figure S15. Principal component analysis of PDAL-Seq reads in alignment blocks where each point represents a cell line x sequencing method (either PDAL-Seq or TruSeq) combination 31
Figure S16. Enrichment of PDAL-Seq reads in human functional genomic element annotations illustrates dysregulation of ssDNA distribution in the cancerous Raji cell line 32
Figure S17. Enrichment of PDAL-Seq and control TruSeq reads in ape satellite arrays. 33
Figure S18. Analysis of PDAL-Seq reads and non-B DNA motifs in Siamang subterminal alpha satellites 34
Figure S19. Analysis of PDAL-Seq reads, DR motifs, and human Hsat2 and Hsat3 arrays in great ape chromosomes, organized by groups of homologous chromosomes 35
Figure S20. Native PAGE and Size exclusion chromatography - multi-angle light scattering (SEC-MALS) analysis of Hsat3 (AATGG)n repeats in vitro 40
Figure S21. An example of S1 digestion test gel 40
**Supplemental References** **41**

### Supplemental Notes

#### Note 1. PDAL-Seq data preprocessing and TruSeq control datasets

At least three technical PDAL-Seq sequencing replicates (with treatment conditions ranging from 20-40 mM permanganate) were analyzed per cell line. Reads were trimmed to remove low-quality base calls at the end, mapped, and PCR duplicates were removed. This produced an average of ~25 million reads with ~19 million reads exhibiting high mapping quality per technical replicate (Table S1). Technical replicates were well correlated within the same cell lines, with Spearman's rank correlation coefficients >0.75 for rates of read mapping in 10-kbp genomic windows for all but one technical replicate (Figure S1). Well-correlated technical sequencing replicates were merged for subsequent analyses, resulting in 50 to 200 million reads per cell line (Table S2).

Because non-B DNA motifs can induce sequencing errors[1] and are enriched at repetitive, difficult-to-map regions of the genome,[2] we also sought to identify potential artifacts arising from erroneous sequencing and/or mapping. Using the same computational pipeline, we analyzed cell-line-matched Illumina TruSeq libraries, which were prepared without the ssDNA enrichment step, but had a similar read length distribution to our PDAL-Seq libraries. Artifacts from sequencing and mapping can thus be identified by cross-referencing PDAL-Seq signals with the TruSeq datasets for each species.

#### Note 2. Biophysical characterization of Hsat2,3 repeats

We sought to characterize the structure of Hsat2,3 sequences *in vitro* due to the high PDAL-Seq signal observed in Hsat2,3 arrays compared to TruSeq controls. We focused on the purine-rich strand because this strand exhibits a higher melting temperature than the corresponding duplex[3] and was the subject of a previous biophysical characterization.[4] This purine-rich strand is known to form a mixture of hairpin and self-complementary duplexes composed of non-cannonical base pairs.[5–10] A recent study used a combination of NMR spectroscopy and x-ray crystallography to determine that the $(ATGGA)_4$ sequence forms an intramolecular hairpin in solution with non-cannonical base pairs (Yatsunyk et. al. NAR 2025).[4] These non-cannonical base pairs are composed of a guanine zipper motif composed of a G-G intercalation flanked by two sheared A-G basepaires and interspaced by A-T base-pairs. This study used relatively low ionic strength conditions, including 50 mM potassium acetate/5 mM magnesium acetate buffer or 25 mM sodium phosphate buffer. We were curious if this guanine zipper motif would form at different repeat lengths. Additionally, we hypothesized that ionic strength conditions that were closer to the 0.1 to 0.5 M monovalent metal ion conditions observed in cells[11] could change the structure. Higher $K^+$ concentrations especially could favor alternative conformations such as a two-tiered G4 structure.

We first denatured and annealed $(AATGG)_4$ in 140 mM LiCl buffer and 100 mM KCl/140 mM LiCl buffer, then collected circular dichroism (CD) spectra. The CD spectra were not KCl-sensitive, indicating that the repeats were not forming a G4 (Figure 6D). Likewise, the CD spectra were consistent with the previously characterized G-zipper hairpin structure for $(ATGGA)_4$, including a trough at ~245 nm, a peak at ~270 nm, and a shoulder at ~280 nm.[4] Thus, these results are consistent with the $(AATGG)_4$ sequence also forming a G-zipper hairpin at higher ionic strength and high $K^+$ conditions.

We next tested the effect of repeat length on structure, where we denatured and annealed $(AATGG)_2$, $(AATGG)_3$, $(AATGG)_5$, and $(AATGG)_6$ in both 140 mM LiCl buffer and 100 mM KCl/140 mM LiCl buffer. We observed the distinct CD signature for the G-zipper hairpin structure at a repeat length of n=3-6 (Figure 6D). CD signatures were also independent of the presence of 100 mM KCl. Indeed, a principal components (PC) analysis of the CD spectra indicated that spectra were classifiable by repeat length but not buffer condition (Figure 6E). Spectra were separated according to overall length in PC1, which composed 85.9% of the variance. Interestingly, PC2 composed 12.8% of the variance and separated spectra by odd versus even repeat numbers (Figure 6E). To further explore these effects, we fractionated samples by molecular weight using native polyacrylamide electrophoresis (Figure S20). All samples had a major band that migrated consistent with a unimolecular weight complex. The oligos with odd repeat number—$(AATGG)_3$ and $(AATGG)_5$— showed a higher propensity for forming high-molecular-weight complexes than even-numbered AATGG repeat constructs.

We confirmed these results with size exclusion chromatography (SEC) - Multi-angle light scattering (MALS). The SEC chromatograms were consistent with the native gel results, showing single major peaks in even-numbered monomer lengths and two peaks for odd-numbered monomer lengths (Figure 7E). We used an in-line MALS, instrument to estimate molecular weights for each peak, which were consistent with low molecular weight peaks being composed of unimolecular structures and higher molecular weight peaks being composed of dimers (Figure 6F, Table S11)

In summary, our biophysical characterization of $(AATGG)_n$ repeats is consistent with a model where the repeats form previously characterized intra-molecular G-zipper hairpins (Figure 7G).[4] The CD indicates that this core structure is surprisingly robust towards ionic strength conditions, repeat number, and register (Figure 7A-B). However, oligos with odd repeat numbers seem to have a higher propensity for forming intramolecular structures than oligos with even repeat numbers (Figure 7C-F), perhaps because the former odd repeat numbers can create a left-over single-stranded repeat that nucleates intramolecular interactions (see Discussion).

### Supplemental methods

#### Optimized PDAL-Seq protocol

Reagent preparation, storage, and handling

**Buffers** were either ordered as nuclease-free stocks or prepared by dissolving high-purity salts in nuclease-free ultra-pure water followed by 0.2 µm filtering using cell-culture grade equipment. Nuclease-free, ultra-pure water was used for all steps.

**Proteinase K (20 mg/mL)** was obtained from MP Biomedicals (Cat #183988) and stored at 4°C.

**RNase A (20 mg/mL)** was obtained from Invitrogen™ PureLink™ (Cat #12091-021) and stored at 4°C.

**Cordycepin triphosphate** was obtained from MilliporeSigma (Cat #C9137) and dissolved to a final concentration of 10 mM. The cordycepin solution was aliquoted, stored at -20°C, and discarded after 1 freeze-thaw cycle.

**Terminal transferase (TdT; NEB 20 units/µL)** was obtained from New England Biolabs® (NEB, Cat #M0315L). On receipt, the enzyme was stored in a -20°C cooler block to minimize temperature fluctuation associated with opening and closing the freezer door.

**S1 Nuclease (79 units/µL)** was obtained from Promega (Cat #M5761). On receipt, the enzyme was stored in a -20°C cooler block to minimize temperature fluctuation associated with opening and closing the freezer door.

**10 mM dNTPs** were obtained from NEB (Cat #N0447), aliquoted, stored at -20°C, and discarded after 1 freeze-thaw cycle.

**T4 DNA Polymerase (3 units/µL)** was obtained from Promega (Cat #M0203). On receipt, the enzyme was stored in a -20°C cooler block to minimize temperature fluctuation associated with opening and closing the freezer door.

**Adapter oligos** were ordered from Integrated DNA Technologies (IDT) with HPLC purification.

- *P5-bio-top:* 5'-/5Biosg/ACACTCTTTCCCTACACGACGCTCTTCCGATCT-3' (5'-end is Biotinylated)
- *P5-bottom:* 5'-/5Phos/AGATCGGAAGAGCGTCGTGTAGGGAAAGAGTGT/3InvdT/-3' (5'-end is Phosphorylated and 3'-inverted deoxy-thymidine).
- P7-top 5'-/5Phos/AGATCGGAAGAGCACACGTCTGAACTCCAGTCAC/3InvdT/-3' (5'-end is Phosphorylated and 3'-end is 3'-inverted deoxy-thymidine)
- P7-bottom 5'-GTGACTGGAGTTCAGACGTGTGCTCTTCCGATCT-3'

Oligos were suspended in nucleus water at a final concentration of 100 µM, aliquoted, and stored at -20°C. Aliquots were discarded after 3 freeze-thaw cycles.

**T4 DNA Ligase (400 units/µL)** was obtained from NEB (Cat #M0202L). On receipt, the enzyme was stored in a -20°C cooler block to minimize temperature fluctuation associated with opening and closing the freezer door. The 10xT4 ligase buffer was aliquoted, stored at -20°C, and discarded after 1 freeze-thaw cycle.

**AMPure XP beads** were obtained from Beckman Coulter (Cat #A63881), completely homogenized vortexing for at least 1 minute, aliquoted into 1 mL single-use vials, and stored at 4°C. Aliquots were discarded after each use to avoid cross-contamination.

**Dynabeads™ KilobaseBINDER™ Kits** were obtained from Invitrogen (Cat #60101) and stored at 4°C.

**NEBNext® End Repair Module** was obtained from NEB (E6050S). On receipt, the enzyme was stored in a -20°C cooler block to minimize temperature fluctuation associated with opening and closing the freezer door. The 10xT4 ligase buffer was aliquoted, stored at -20°C, and discarded after 1 freeze-thaw cycle.

**Indexing PCR primers** were obtained from NEB (Cat #E7600S), aliquoted, stored at -20°C, and discarded after one-time use to avoid cross-contamination.

**Phusion HSII Polymerase** was obtained from ThermoFisher Scientific (Cat #F549S). On receipt, the enzyme was stored in a -20°C cooler block to minimize temperature fluctuation associated with opening and closing the freezer door.

Day 1: Harvest and treatment

**Potassium permanganate ($KMnO_4$) preparation.** Fresh 250 mM $KMnO_4$ was prepared by dissolving $KMnO_4$ powder in nuclease-free water in a conical tube. The tube was covered with foil to protect it from light and incubated in a 37℃ shaker for >2 hours to ensure complete homogenization of the $KMnO_4$ solution. The $KMnO_4$ was diluted in water to a working concentration of 20 mM or 40 mM and incubated at 37℃ for 1 hour prior to treatment.

**Treatment (A T75 flask of adherent cells is used as an example for the experiment)** Cells were grown to 5 million cells per T75 flask. The samples were processed one at a time in a laminar flow hood so that the entire processing time from culture to lysis was less than 5 min. The medium was removed and cells were washed twice with ~10 mL of pre-warmed 37℃ Phosphate Buffered Saline (PBS) without divalent calcium/magnesium. Excess PBS was removed, and the cells were incubated in 2 mL of pre-warmed LOW SALT BUFFER (15 mM Tris-HCl pH 7.5, 15 mM NaCl, 60 mM KCl, 300 mM Sucrose, 0.5 mM EGTA) for 60 seconds at room temperature. Cells were treated by adding 667 µL of either prewarmed water (Control) or $KMnO_4$ working solutions. A timer was started on the addition of the treatment solution, while samples were gently mixed and stored at 37℃. The treatment solution was quenched and cells were lysed at exactly 80 seconds by the addition of 2.67 mL of pre-warmed STOP SOLUTION (50 mM EDTA pH 8.0, 700 mM beta-Mercaptoethanol, 1% SDS). Samples were mixed well and visually inspected for complete quenching of $KMnO_4$ and lysis (the purple color of the $KMnO_4$ will disappear completely, and globs of DNA will be visible in the flask). Treated samples were set aside at room temperature while other samples were processed. The lysate was then digested with 300 µg/mL proteinase K at 37℃ for ~20 hours (overnight).

Day 2: DNA extraction

**DNA extraction and precipitation.** DNA was extracted by adding 6 mL of phenol:chloroform:isoamylalcohol (25:24:1) to the lysate and inverting the tubes 10 times to mix well. Samples were spun at 2,000 xg for 5 min to separate phases. The supernatant was transferred to a new 50 mL tube, then mixed with 4 mL of 5M ammonium acetate (final con. 2M) and 20 mL of ice-cold absolute ethanol. Samples were mixed by gently inverting the tubes until phase lines disappeared and DNA precipitates were visible. DNA was further incubated at -20℃ for >30 min. DNA was pelleted by spinning for 60 minutes at 4℃, 2,000 x g before the supernatant was removed by pouring. The pellet was dislodged in 2 mL ice-cold 80% ethanol and then repelleted for 5 minutes at 4℃, 2,000 xg. Excess supernatant was removed, and the pellet was dried inverted for 30 minutes and upright for 30 min in a laminar flow hood. DNA was resuspended in 1 mL of 10 mM Tris-HCl pH 8.0. To avoid DNA shearing, only tap the tubes gently to mix the samples (No pipetting). Samples were incubated at room temperature for ~20 hours (overnight).

Day 3: RNase treatment

**RNase treatment, DNA extraction, and precipitation**. DNA was transferred to a 1.5-mL low-binding tube. 2.5 µL of 20 mg/mL RNase A was added to 1 mL sample, and samples were mixed by inverting to avoid DNA shearing. RNA was digested at 37℃ for 1 hour. DNA was extracted by adding 1 mL of phenol:chloroform:isoamylalcohol (25:24:1) to the lysate and inverting the tubes 10 times to mix well. Samples were spun at 2,000 xg for 5 min to separate phases. The supernatant was transferred to a new 15 mL tube, then mixed with 0.667 mL of 5M ammonium acetate (final con. 2M) and 3.34 mL of ice-cold absolute ethanol. Samples were mixed by gently inverting the tubes until phase lines disappeared and DNA precipitates were visible. DNA was further incubated at -20℃ for >30 min. DNA was pelleted by spinning for 60 minutes at 4℃, 2,000 x g before the supernatant was removed by pouring. The pellet was dislodged in 1 mL ice-cold 80% ethanol and then repelleted for 5 minutes at 4℃, 2,000 xg. Excess supernatant was removed, and the pellet was dried inverted for 30 minutes and upright for 30 min in a laminar flow hood. DNA was resuspended in 0.29 mL of 10 mM Tris-HCl pH 8.0. To avoid DNA shearing, only tap the tubes gently to mix the samples (No pipetting). Samples were incubated at room temperature for ~20 hours (overnight).

Day 4: Block 3'-DNA ends

**Block 3'-DNA ends** DNA were transferred to a 1.5-mL low-binding tube and mixed with 51.5 µL of water, 50.0 µL of 10x TdT Buffer (NEB), 100 µL of 25mM Cobalt Chloride, 6.0 µL of 10 mM Cordycepin Triphosphate, and 2.5 µL of Terminal Transferase (TdT; NEB 20 units/µL). Samples were inverted to mix and incubated at 37℃ for 1 hour. Reactions were stopped with 20 µL of 0.5M EDTA pH 8.0. Then TdT was digested by adding proteinase K to a final concentration of 150 µg/mL and incubating at 37℃ for 1 hour.

**DNA extraction and precipitation.** DNA was extracted by adding 0.5 mL of phenol:chloroform:isoamylalcohol (25:24:1) to the lysate and inverting the tubes 10 times to mix well. Samples were spun at 12,000 xg for 5 min to separate phases. The supernatant was transferred to a new 15-mL low-binding tube, then mixed with 0.333 mL of 5M ammonium acetate (final con. 2M) and 2.0 mL of ice-cold absolute ethanol. Samples were mixed by gently inverting the tubes until phase lines disappeared and DNA precipitates were visible. DNA was further incubated at -20℃ for >30 min. DNA was pelleted by spinning for 60 minutes at 4℃, 2,000 x g before the supernatant was removed by pouring. The pellet was dislodged in 1 mL ice-cold 80% ethanol and then repelleted for 5 minutes at 4℃, 2,000 xg. Excess supernatant was removed, and the pellet was dried inverted for 30 minutes and upright for 30 min in a laminar flow hood. DNA was resuspended in 0.6 mL of 10 mM Tris-HCl pH 8.0. To avoid DNA shearing, only tap the tubes gently to mix the samples (No pipetting). Samples were incubated at room temperature for 1 hour.

**Second precipitation to remove residual cordycepin.** The DNA was mixed with 0.4 mL of 5M ammonium acetate (final con. 2M) and 2.0 mL of ice-cold absolute ethanol. Samples were mixed by gently inverting the tubes until phase lines disappeared and DNA precipitates were visible. DNA was further incubated at -20 ℃ for >30 min. DNA was pelleted by spinning for 60 minutes at 4℃, 2,000 xg before the supernatant was removed by pouring. The pellet was dislodged in 1 mL ice-cold 80% ethanol and then repelleted for 5 minutes at 4℃, 2,000 xg. Excess supernatant was removed and the pellet was dried inverted for 30 minutes and upright for 30 min in a laminar flow hood. DNA was resuspended in 0.5 mL of 10 mM Tris-HCl pH 8.0. To avoid DNA shearing, only tap the tubes gently to mix the samples (No pipetting). Samples were incubated at 4℃ until the resuspended high-molecular-weight DNA formed a homogeneous solution (~3 days or over a weekend).

Day 5: S1 Nuclease digestion trial (find the optimal concentration)

**S1 Nuclease digestion trial.** S1 nuclease activity varies by lot, age, and DNA substrate. The following procedure was used to determine optimum S1 concentrations for a bulk S1 digest.

DNA concentration and homogeneity was confirmed by determining the DNA concentration with Qubit high-sensitivity dsDNA assay (Invitrogen) in triplicate using 1 µL samples. Samples taken from different positions in the solution (top, middle, bottom) should exhibit consistent Qubit readings. Test S1 digestion reactions were set up on ice. As an example, the following experiment tested the efficacy of a 200 ng DNA and 0.4 units of S1 nuclease in a 15 µL reaction volume. S1 nuclease was diluted with water to a final concentration of 0.4 unit/µL and kept on ice. Prepare a DNA master mix by combining 4 µL of 100 ng/µL DNA (400 ng DNA total), 3 µL 10x S1 Nuclease Reaction Buffer (0.5M sodium acetate pH 4.5, 2.8M NaCl, 45mM $ZnSO_4$), and 21 µL of nuclease-free water on ice (this is a 28 µL total volume). The DNA master mix was tapped to mix gently and spun down briefly. The control reaction was prepared by mixing 14 µL DNA master mix with 1 µL of water on ice (200 ng DNA). The S1 nuclease reaction was prepared by mixing 14 µL DNA master mix with 1 µL of 0.4 unit/µL S1 nuclease on ice (200 ng DNA). Reactions were quickly tapped to mix gently, spun down, and incubated for 20 min on a preheated 37℃ thermocycler. The reactions were stopped at exactly 20 min by the addition of 15 µL of S1 quenching solution (20 mM EDTA required to sequester the $Zn^{2+}$ S1 enzyme cofactor, 20% glycerol, 0.025% Bromophenol blue). Reaction products were analyzed on a 0.6% agarose gel in 1xTAE buffer and 1xGel Red working reagent. The gel was imaged on a BioRad Gel-Doc Go imager and products were compared to a NEB 1 kb ladder.

Figure S21 is provided as an example of a good S1 reaction, which has the following features. (1) S1 nuclease should reduce high-molecular-weight DNA clumping in permanganate-treated samples. (2) There should be a consistently strong band migrating at >10 kbp. (3) An untreated DNA control should not show electromobility changes after S1 treatment. (4) The $KMnO_4$ will produce a stronger smear running from 1 kbp to 10 kbp following S1 treatment. (5) The intensity of the S1-induced smear should be proportional to the $KMnO_4$ concentration in the treatment.

Day 6: Large scale S1 Nuclease digestion

The large-scale S1 nuclease digest used the same ratio of S1 nuclease, to DNA mass, to volume as the test S1 reaction. For example, if 20 µg of end-blocked-DNA is available, the reaction components from the test reaction would have been scaled by 20 µg total/0.2 µg per test reaction = 100x. Reactions were prepared on ice, S1 nuclease was added last, the reactions were tapped to mix quickly, and incubated on a preheated 37℃ heat block. The reactions were stopped at exactly 20 min by the addition of 0.5 M EDTA to a final concentration of 10 mM (a >2-fold molar excess to the $Zn^{2+}$ S1 enzyme cofactor). DNA was extracted by adding an equal volume of phenol:chloroform:isoamylalcohol (25:24:1) to the reaction and inverting the tubes 10 times to mix well. Samples were spun at 12,000 xg for 5 min to separate phases. The supernatant was transferred to a new low-binding tube, then mixed with ½ volume of 5M ammonium acetate (final con. 2M), and 3x-volume of ice-cold absolute ethanol, and 1 µL glycogen solution as a carrier. Samples were mixed by gently inverting the tubes until phase lines disappeared and DNA precipitates were visible. DNA was further incubated at -20℃ for >30 min. DNA was pelleted by spinning for 30 minutes at 4℃, 2,000 x g before the supernatant was discarded by pouring. The pellet was dislodged in 0.5 mL ice-cold 80% ethanol and then repelleted for 5 minutes at 4℃, 2,000 xg. Excess supernatant was removed, and the pellet was dried inverted for 30 minutes and upright for 30 min in a laminar flow hood. DNA was resuspended in 84 µL of 10 mM Tris-HCl pH8.0, and gently tapped to mix thoroughly, not pipetted. Samples were incubated at 4℃ overnight before proceeding.

Day 7: End Repair with T4 DNA Polymerase

The DNA concentration was measured in triplicate using the Qubit High Sensitivity assay to determine the amount of T4 DNA Polymerase needed (1 unit of T4 DNA Polymerase was used to repair 1 µg DNA). The end-repair reactions were prepared in a PCR tube on ice, mixing 80 µL DNA, 10 µL of 10x NEB Buffer r2.1, 1.0 µL of dNTPs (10mM each), and T4 DNA Polymerase to a final amount of 1 unit polymerase/ 1 µg DNA. Nuclease-free water was added to bring the final volume to be 100 µL. The reaction was incubated for 15 min at 12°C in a thermocycler, stopped with 2 µL 0.5M EDTA, and enzymes were heat-inactivated at 75°C for 20 min. The volume was increased by adding 100 µL Tris-HCl buffer pH8.0 to the sample. DNA was extracted, precipitated, and resuspended using the same protocol as the large-scale S1 nuclease digestion.

Day 8: P5 adapter ligation

**P5 Adapter Preparation and Ligation** An annealing reaction was prepared by mixing 1 µL 100 µM P5-bio-top, 1 µL 100 µM P5-bottom, 1 µL 10xT4 DNA ligase buffer, and 7 µL 10 mM Tris-HCl pH8.0 in a 0.2 mL PCR tube. Adapters were heated to 95 °C for 5 minutes in a thermocycler and then slowly cooled to room temperature for at least 1 hour. Ligation reactions were prepared by combining 5 µL water, 79 µL end-repaired DNA sample, 10 µL 10xT4 DNA ligase buffer, 1 µL of denatured and annealed P5 adapter, and 5 µL of T4 DNA Ligase (400 units/µL NEB) in a 0.2-ml PCR tube on ice. Samples were mixed by gently tapping and incubated at 16 °C overnight (>18 hours).

Day 9: Bead size-selection, sonification, streptavidin pull-down, end repair, P7 ligation

**Purification of high-molecular-weight DNA with AMPure XP Beads.** AMPure XP bead aliquots were warmed to room temperature for at least 30 minutes, and tapped/vortexed for at least 1 minute to ensure complete resuspension and homogenization of beads. Ligation reactions were warmed to room temperature, and the volume was increased to 200 µL by adding 100 µL of 10 mM Tris-HCl pH8.0. 180 µL of AMPure beads (0.9 volumes beads) were added to the samples, and samples were mixed thoroughly by pipetting. The bead/sample mixture was incubated for 5 minutes at room temperature on a rotary mixer. High-molecular-weight DNA bound to beads was pelleted on a magnetic stand for 5 minutes. The supernatant was removed with a pipette and discarded. Beads were washed twice by dislodging the pellet with 400 µL of freshly prepared 80% ethanol (Pelleting on the magnetic stand and discarding ethanol after each wash). Residual ethanol was pipetted off while the beads were dried for 3-5 minutes (visible ethanol should be gone, but beads should remain dark and glossy). Samples were removed from the magnetic stand, resuspended in 140 µL of TE buffer, and incubated at room temperature for 5 minutes to elute high-molecular-weight DNA. Beads were pelleted on a magnetic stand for 5 minutes, and 136 µL of supernatant containing high-molecular-weight DNA was retained.

**DNA Sonication** 130 µL of the high-molecular-weight sample was fragmented to a target peak size of 350bp using a Covaris Focused-ultrasonicator M220, Covaris microTube-AFA Fiber Pre-Slit Snap Cap (Part.No. 520045) tubes, and the following duty cycle: Peak incident power 75; Duty factor 10; Cycle per burst 200; Treatment time 80 seconds. Fragment sizes were checked on a 1% TAE agarose gel and an Agilent Bioanalyzer instrument.

**Streptavidin Beads Pull-down** The sonicated DNA sample concentration was determined using a Qubit high-sensitivity dsDNA assay. Take 0.5 µg of sonicated DNA to dilute with TE to a final volume of 130 µL. DNA fragments that were ligated to P5 Illumina adapters were purified with a Dynabeads KilobaseBINDER Kit (Invitrogen). First, the kit was warmed to room temperature for >30 min. Second, 50 µL volume of Dynabeads was washed with 200 µL of the Binding Solution provided by the manufacturers, and resuspended in a second 200 µL volume of Binding Solution. The diluted 130 µL (0.5 µg) DNA sample was then added to the 200 µL Dynabeads suspended in the binding solution. Samples were mixed gently by inverting the tubes to avoid excessive bubbles. The samples were then incubated at room temperature for 1.5 hours on a rotating mixer at 10 rpm to maintain beads in suspension. The beads were washed twice with 500 µL of Washing Solution (2.0 M NaCl, 1 mM EDTA, 10 mM Tris-HCl pH7.5) and then twice with 400 µL of 10 mM Tris-HCl pH7.5. Samples were resuspended in 400 µL of 10 mM Tris-HCl pH7.5, and stored at 4 °C until the next step.

**End repair** DNA fragments were repaired with the NEBNext® End Repair Module. The 10 mM Tris-HCl pH 7.5 was removed and discarded, and then beads were resuspended in 100 µL of end-repair mix (85 μL of water, 10 μL of 10x NEBNext End Repair Reaction Buffer, and 5 μL of NEBNext End Repair Enzyme Mix, prepared on ice and mixed by pipetting). The bead mixture was incubated for 30 min at 20°C in a thermocycler with a heated lid set to its lowest setting. The beads were washed three times with 200 μL of 10 mM Tris-HCl pH7.5, and stored in 200 μL of 10mM Tris-HCl pH7.5 at 4°C until the next step.

**P7 Adapter ligation and Clean up** An annealing reaction was prepared by mixing 1 μL 100 μM P7-top, 1 μL 100 μM P7-bottom, 2 μL 10xT4 DNA ligase buffer, and 16 μL 10 mM Tris-HCl pH8.0 in a 0.2 mL PCR tube. Adapters were heated to 95°C for 5 minutes in a thermocycler and then slowly cooled to room temperature for at least 1 hour. A 99 μL ligation mixture volume for each sample was prepared by combining 88 μL water, 10 μL 10xT4 DNA ligase buffer, and 2 μL of denatured and annealed P7 adapter on ice. The ligation mixture was mixed thoroughly by pipetting. Then the 10 mM Tris-HCl pH7.5 storage solution was removed from the sample, and the sample was resuspended in 99 μL of ligation mixture. The reaction was pre-incubated on ice for 5 min. Last, 1 μL of T4 DNA Ligase (400 units/ul NEB) was added to the reaction, reactions were thoroughly mixed by pipetting, and incubated at room temperature overnight on a rotating mixer at 10 rpm.

Day 10: PCR

**PCR** Beads were washed twice with 200 μL Washing Solution (2.0 M NaCl, 1 mM EDTA, 10 mM Tris-HCl pH 7.5) and three times with 200 μL 10mM Tris-HCl pH7.5 buffer. The samples were stored in 200 μL of 10mM Tris-HCl pH7.5 at 4 °C until the PCR step. A 100 μL PCR mix was prepared for each sample by combining 75 μL of water, 20 μL of 5x Phusion HF buffer, 2 μL of dNTPs (10 mM each), 1 μL of 10 μM P5 indexing primer, 1 μL of 10 μM P7 indexing primer, and 1 μL of Phusion HSII Polymerase (2 U/μL) on ice. The storage buffer was then removed from the sample, and the sample was resuspended in the PCR mix. Samples were mixed thoroughly and transferred to a PCR thermocycler. The PCR program follows:

- 98 °C denaturation 30 seconds
- Cycle:
    - 98 °C denaturation 10 seconds
    - 65 °C annealing 20 seconds
    - 72 °C extension 20 seconds
- Repeat cycle 15 times
- 72 °C final extension for 5 minutes
- 4 °C hold

The cycle was interrupted at 15 minutes (~6 cycles) to resuspend the beads by mixing with a multichannel pipette.

**Purification of PCR products.** AMPure XP bead aliquots were warmed to room temperature for at least 30 minutes, and tapped/vortexed for at least 1 minute to ensure complete resuspension and homogenization of beads. The PCR reaction was warmed to room temperature, and Dyna-beads were pelleted on a magnetic stand for 5 minutes before the supernatant containing PCR products was retained. The PCR supernatant volume was increased to 100 μL (+/- 5 μL) with 10 mM Tris-HCl pH 8.0 if there was volume loss. 90 μL of AMPure XP beads were added to the PCR products (0.9 volume) and mixed thoroughly by gently pipetting. Samples were incubated at room temperature for 5 minutes, and DNA bound to AMPure beads were pelleted on a magnetic stand for 5 min. The supernatant was discarded, and the beads were washed twice with 200 μL of freshly prepared 80% ethanol. The bead pellet was removed from the magnet, dislodged into the 80% ethanol, and repelleted during each wash. The ethanol was removed, and the beads were dried for 5 minutes. Beads were resuspended in 30 μL of Tris-HCl pH8.0 buffer, incubated at room temperature for 5 min, repelleted on the magnetic stand for 5 min, and 28 μL of supernatant containing the final library was retained.

**Library quantification and sequencing.** Library concentration and purity were assessed using Qubit (Invitrogen), Microvolume UV-absorbance spectroscopy (Nanodrop, ThermoFisher Scientific), qPCR, and Agilent Bioanalyzer analysis - or equivalent technologies. Libraries were sequenced at the Pennsylvania State University Huck Genomics Core Facility on the Illumina NextSeq 2000 using a NextSeq 2000 P1 150-nucleotide paired read kit for all but the HEK293T sample. The HEK239T sample used a NextSeq 2000 P3 100-nucleotide paired read kit.

### Supplemental figures

##### Figure S1. Spearman's rank correlation coefficients between PDAL-Seq technical sequencing replicates in differently sized genomic windows

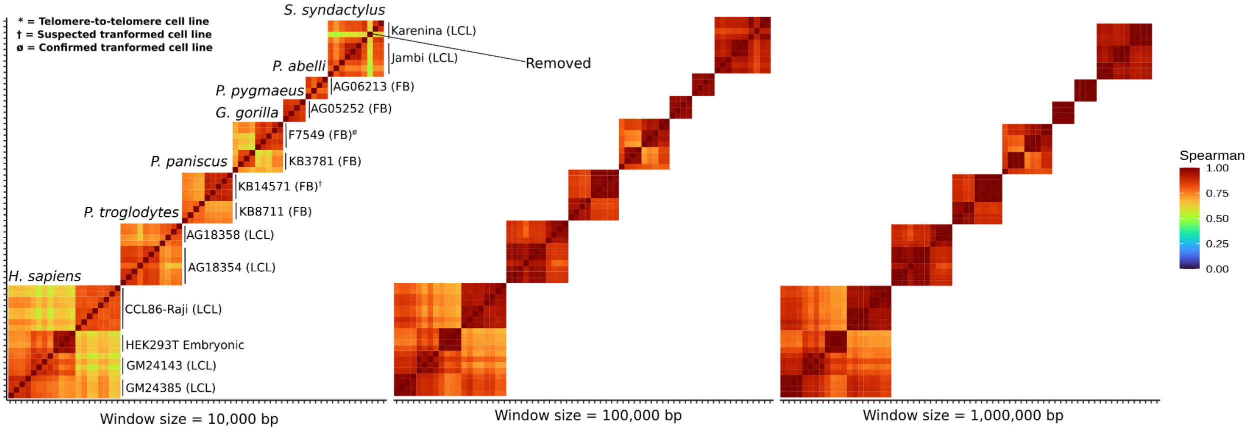

**Figure S2 Comparison of PDAL-Seq read density with non-B DNA motif density for each cell line**

Chromosomes with centromeres marked with grey circles (top), coverage by human HSat2 and HSat3 satellite arrays (as annotated by RepeatMasker) in 1-Mbp windows, coverage by non-B DNA motifs in 1-Mbp windows (middle), and number of PDAL-Seq reads per Mbp (bottom). Plots share an x-axis that corresponds to chromosomes, colored left to right by chromosome. A 99% winsorization transformation was applied to reduce the effect of spurious outliers on non-B motifs and read densities. **(A)** human; **(B)** chimpanzee; **(C)** bonobo; **(D)** gorilla; **(E)** Bornean orangutan; **(F)** Sumatran orangutan; **(G)** siamang.

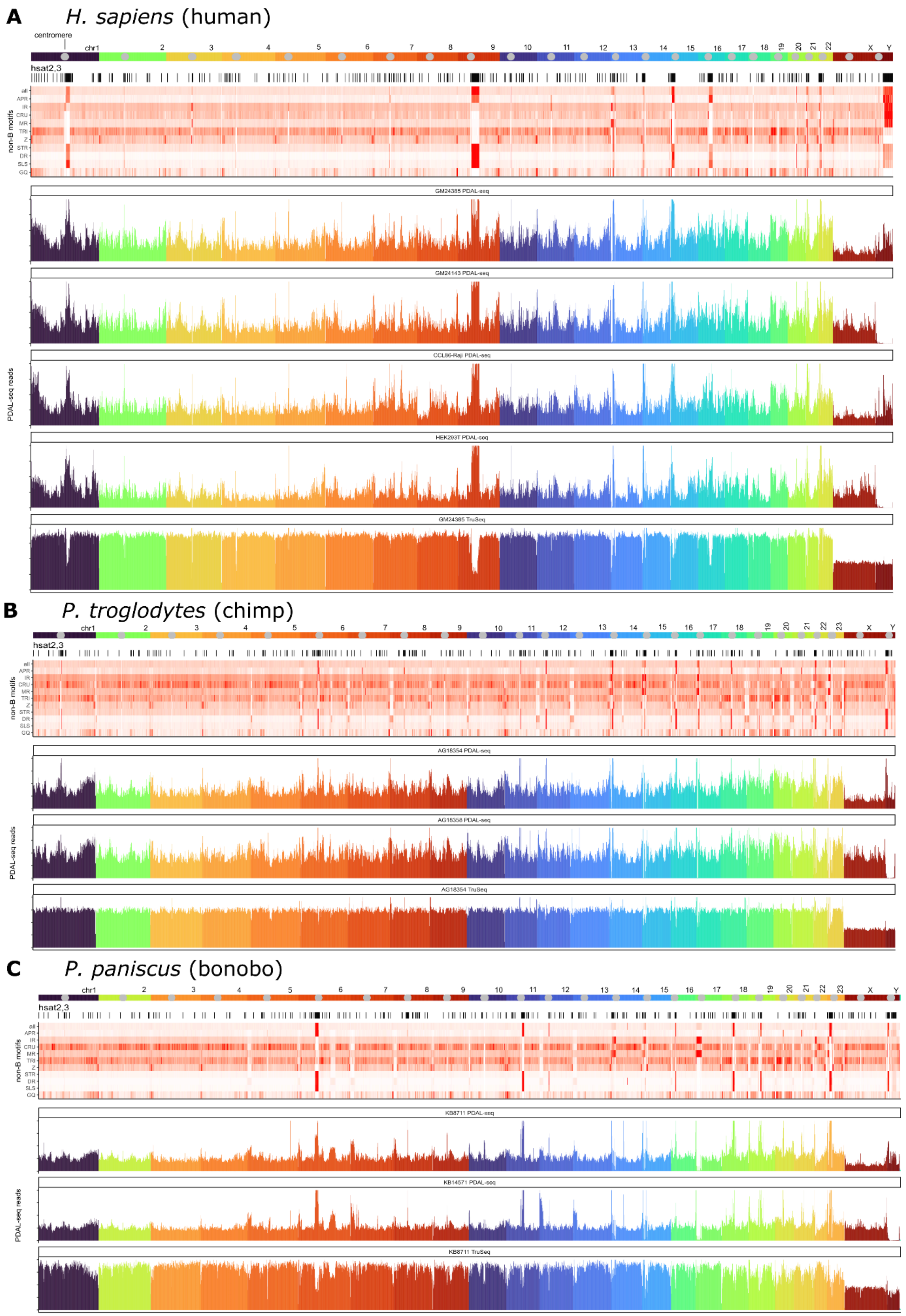
A
H. sapiens (human)
centromere
chr1
hsat2,3
non-B motifs
all
APR
IR
CRU
MR
TRI
Z
STR
DR
SLS
GQ
GM24385 PDAL-seq
GM24143 PDAL-seq
CCL86-Raji PDAL-seq
HEK293T PDAL-seq
GM24385 TruSeq
PDAL-seq reads
B
P. troglodytes (chimp)
AG18354 PDAL-seq
AG18358 PDAL-seq
AG18354 TruSeq
C
P. paniscus (bonobo)
KB8711 PDAL-seq
KB14571 PDAL-seq
KB8711 TruSeq

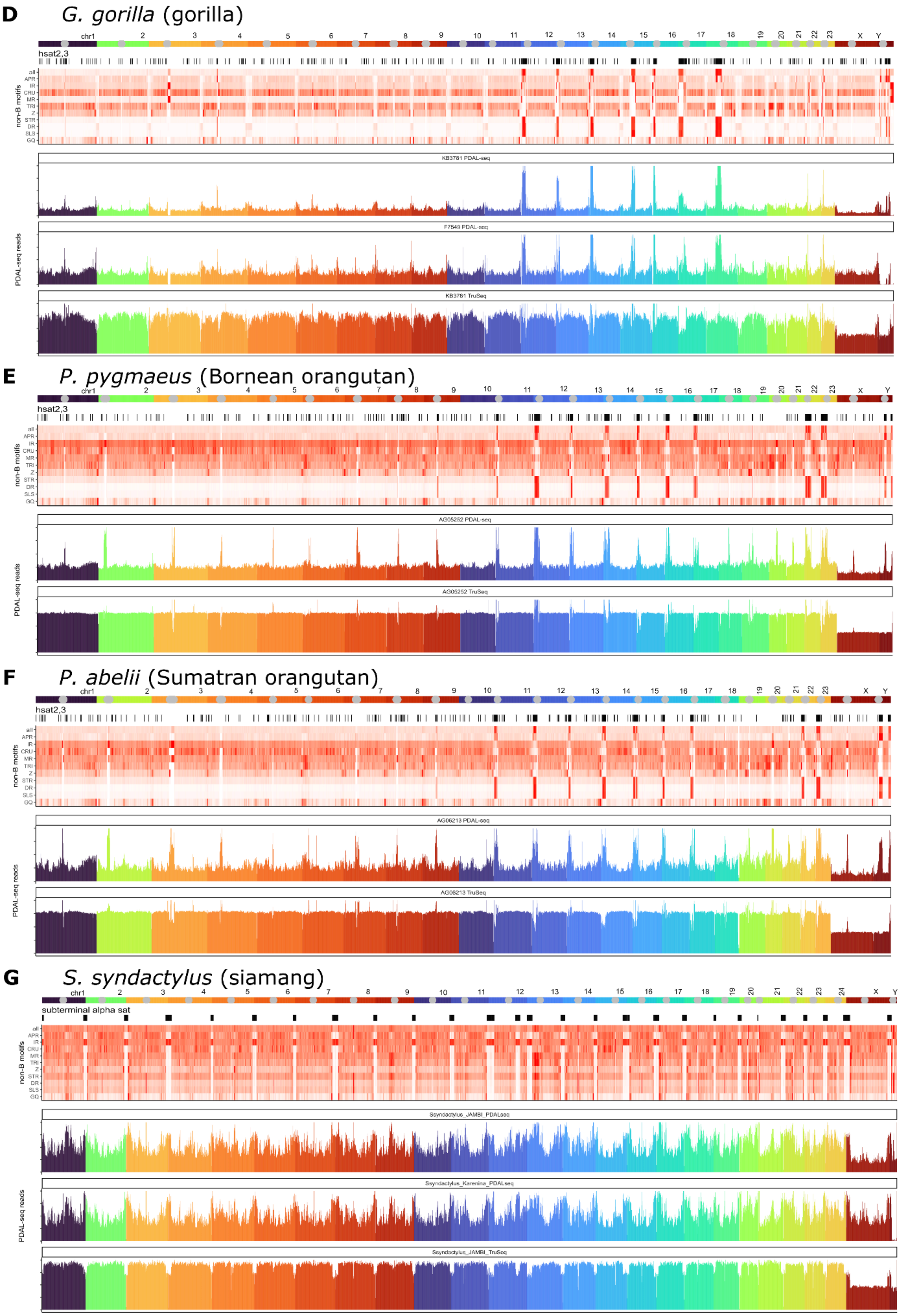
D
G. gorilla (gorilla)
hsat2,3
non-B motifs
KB3781 PDAL-seq
F7549 PDAL-seq
KB3781 TruSeq
PDAL-seq reads
E
P. pygmaeus (Bornean orangutan)
hsat2,3
non-B motifs
AG05252 PDAL-seq
AG05252 TruSeq
PDAL-seq reads
F
P. abelii (Sumatran orangutan)
hsat2,3
non-B motifs
AG06213 PDAL-seq
AG06213 TruSeq
PDAL-seq reads
G
S. syndactylus (siamang)
subterminal alpha sat
non-B motifs
Ssyndactylus_JAMBI_PDALseq
Ssyndactylus_Karenina_PDALseq
Ssyndactylus_JAMBI_TruSeq
PDAL-seq reads
all
APR
IR
CRU
MR
TRI
Z
STR
DR
SLS
GQ

**Figure S3. PDAL-Seq read density versus non-B DNA motif density for each cell line, separated by non-B DNA motif**

Coverage by non-B DNA motifs in 1-Mbp windows (left) versus the number of PDAL-Seq reads per Mbp (bottom). A 99% winsorization transformation was applied to reduce the effect of spurious outliers on non-B motifs and read densities. Blue and green points correspond to windows from chrX and chrY, respectively. **(A)** human; **(B)** chimpanzee; **(C)** bonobo; **(D)** gorilla; **(E)** Bornean orangutan; **(F)** Sumatran orangutan; **(G)** siamang.

**A** *H. sapiens*

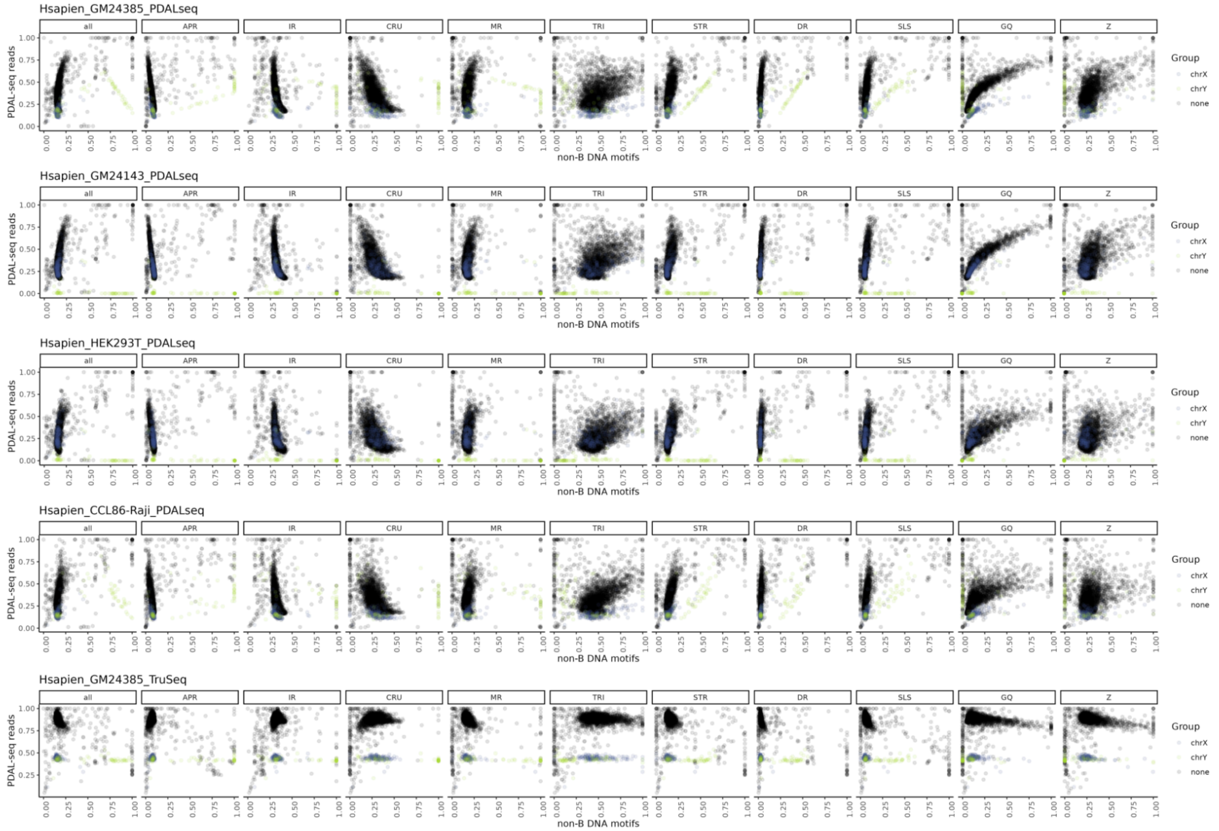


**B** *P. troglodytes*

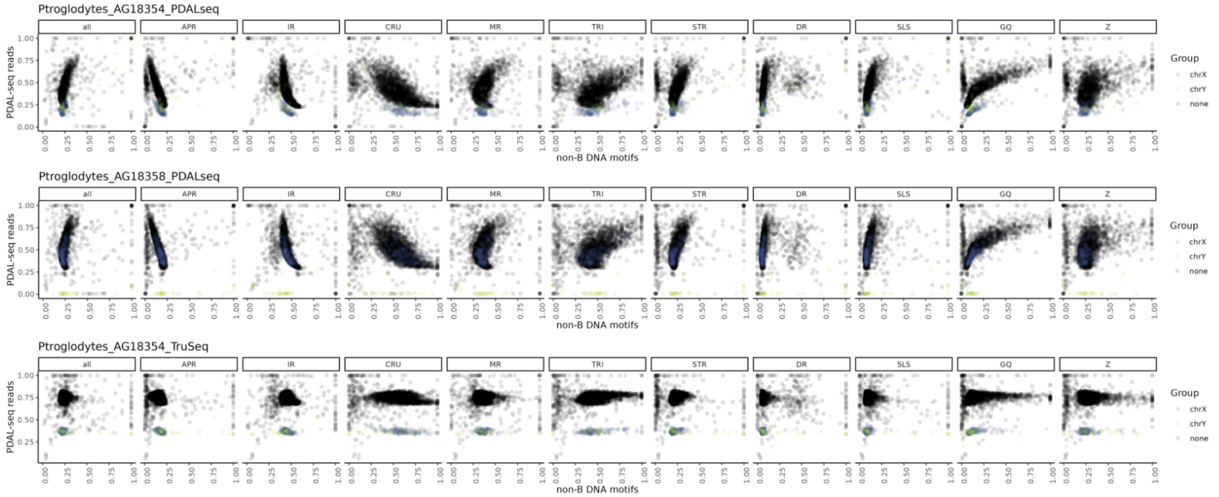

**C** *P. paniscus*

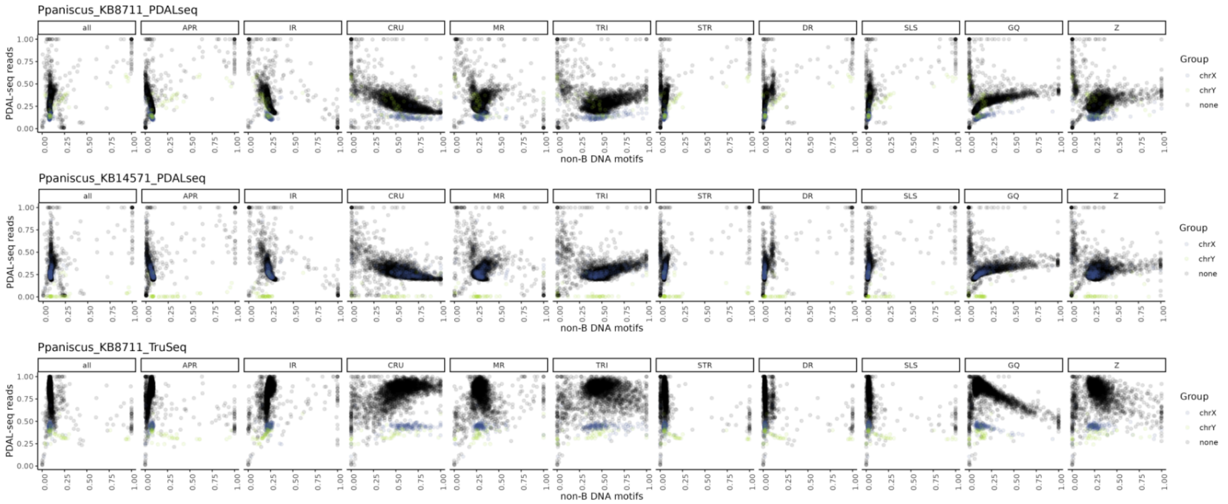


**D** *G. gorilla*

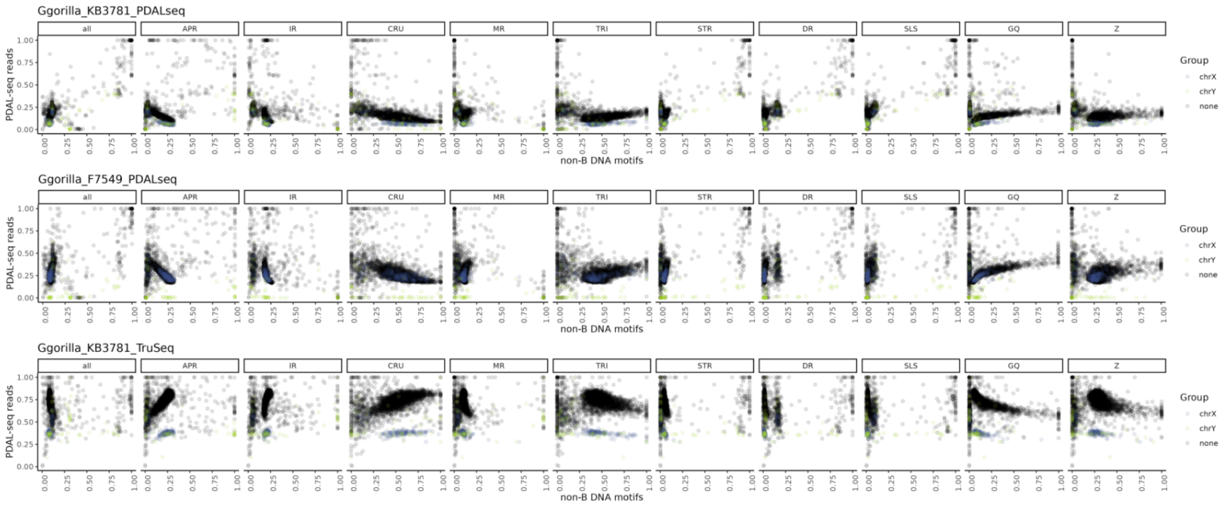


**E** *P. pygmaeus*

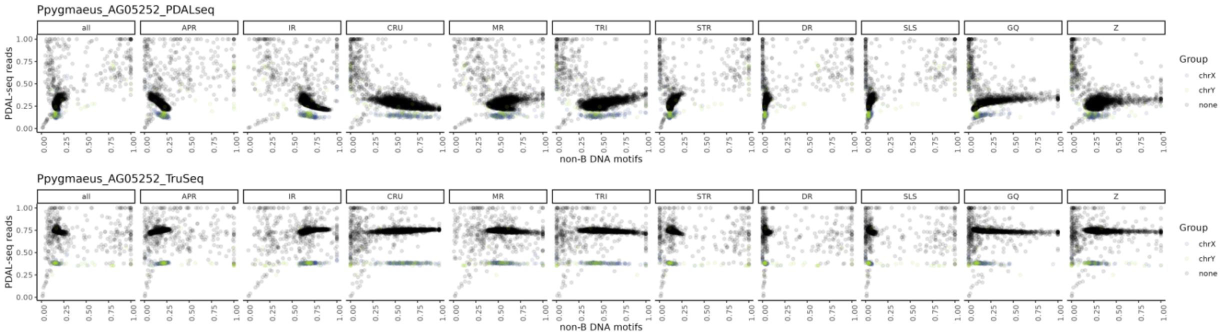

**F** *P. abelii*

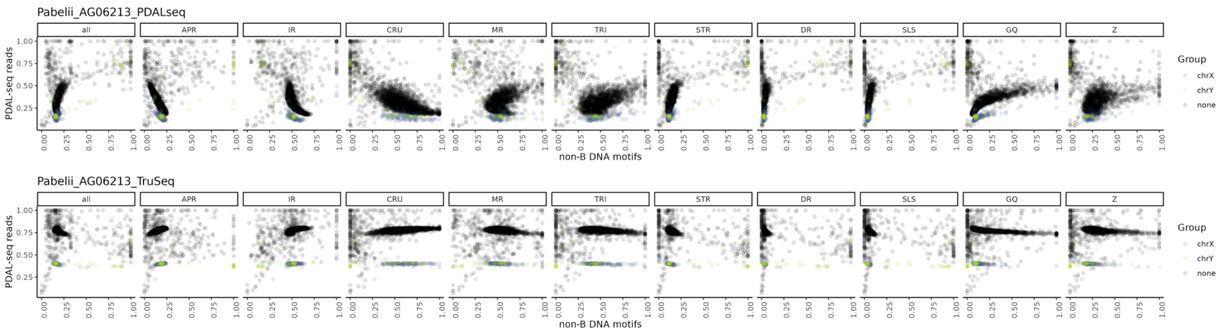


**G** *S. syndactylus*

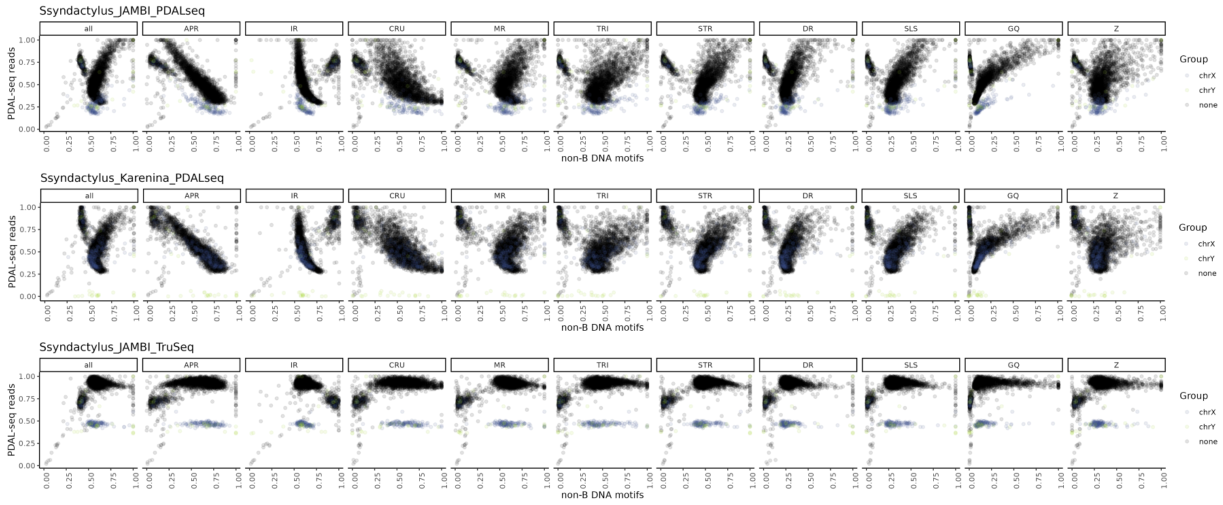

**Figure S4. Poisson generalized linear modeling (GLM) to identify the relationship between ssDNA versus non-B DNA motifs in 1-kbp windows corresponding to uniquely mapping regions of the genome**

The y-axis is the GLM parameter values for non-B DNA motifs in global fits. Bars and error bars represent the mean and standard error of 14 cell lines for PDAL-Seq and 7 cell lines for TruSeq. Sets correspond to "*dedup with GC"* ∈ {GC, APR, CRU, TRI, DR, G4, Z} and "*dedup no GC"* ∈ {APR, CRU, TRI, DR, G4, Z} (see Methods).

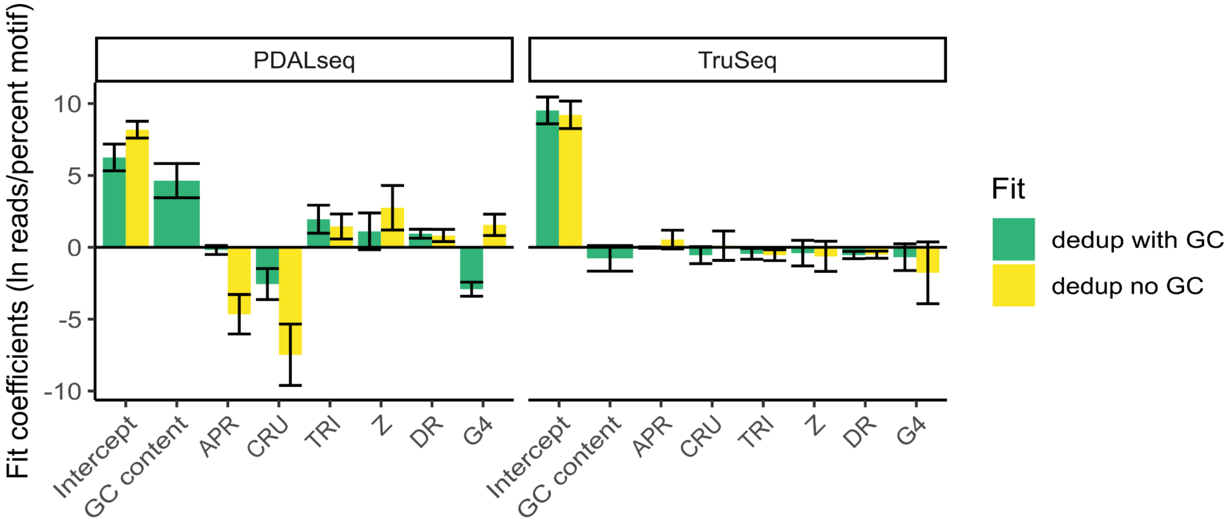

**Figure S5. Comparison of PDAL-Seq to 2 published G4 ChiP-Seq, 14 G4 CUT&Tag, and 6 R-loop CUT&Tag datasets in the HEK293T cell line.**
**(A)** Heatmap showing Spearman's rank correlation coefficients between reads mapping to 1-Mbp windows in different experiments. Data compiled from [12–17]. **(B)** Genome-wide comparison of a representative G4 CUT&Tag dataset, our PDAL-Seq data set, and a representative R-loop CUT&Tag dataset. Plots share an x-axis which corresponds to chromosomes chr1 to chr22 , chrX, and chrY according to the ideogram colored left to right by chromosome. A 99% winsorization transformation was applied to reduce the effect of spurious outliers on non-B motifs and read densities.

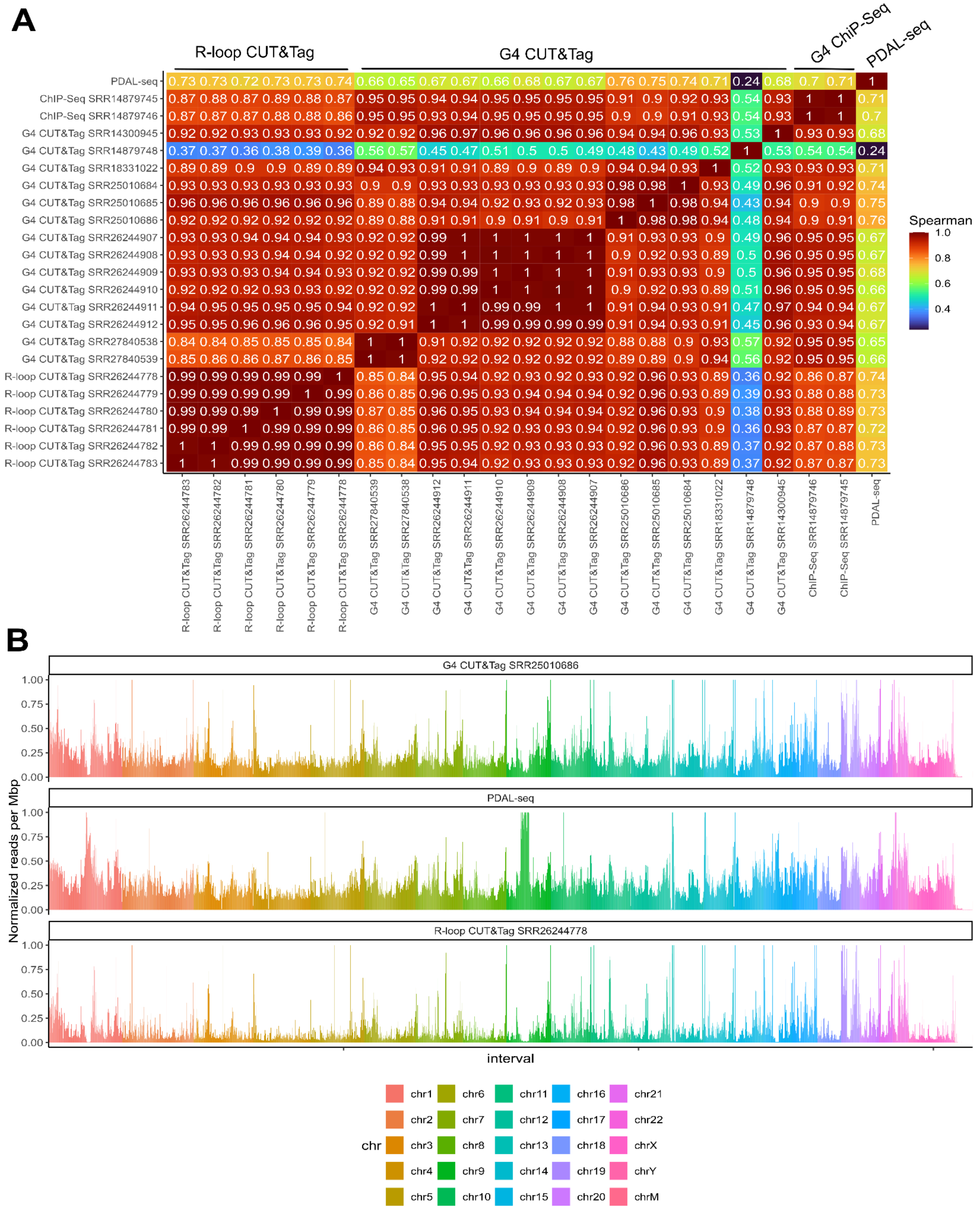
A
R-loop CUT&Tag
G4 CUT&Tag
G4 ChIP-Seq
PDAL-seq
Spearman
B
G4 CUT&Tag SRR25010686
PDAL-seq
R-loop CUT&Tag SRR26244778
Normalized reads per Mbp
interval
chr

**Figure S6. Principal component analysis of PDAL-Seq reads in alignment blocks**
Each point represents a cell line and sequencing method (PDAL-Seq or TruSeq) combination. Alignments were parsed according to different groups of species.

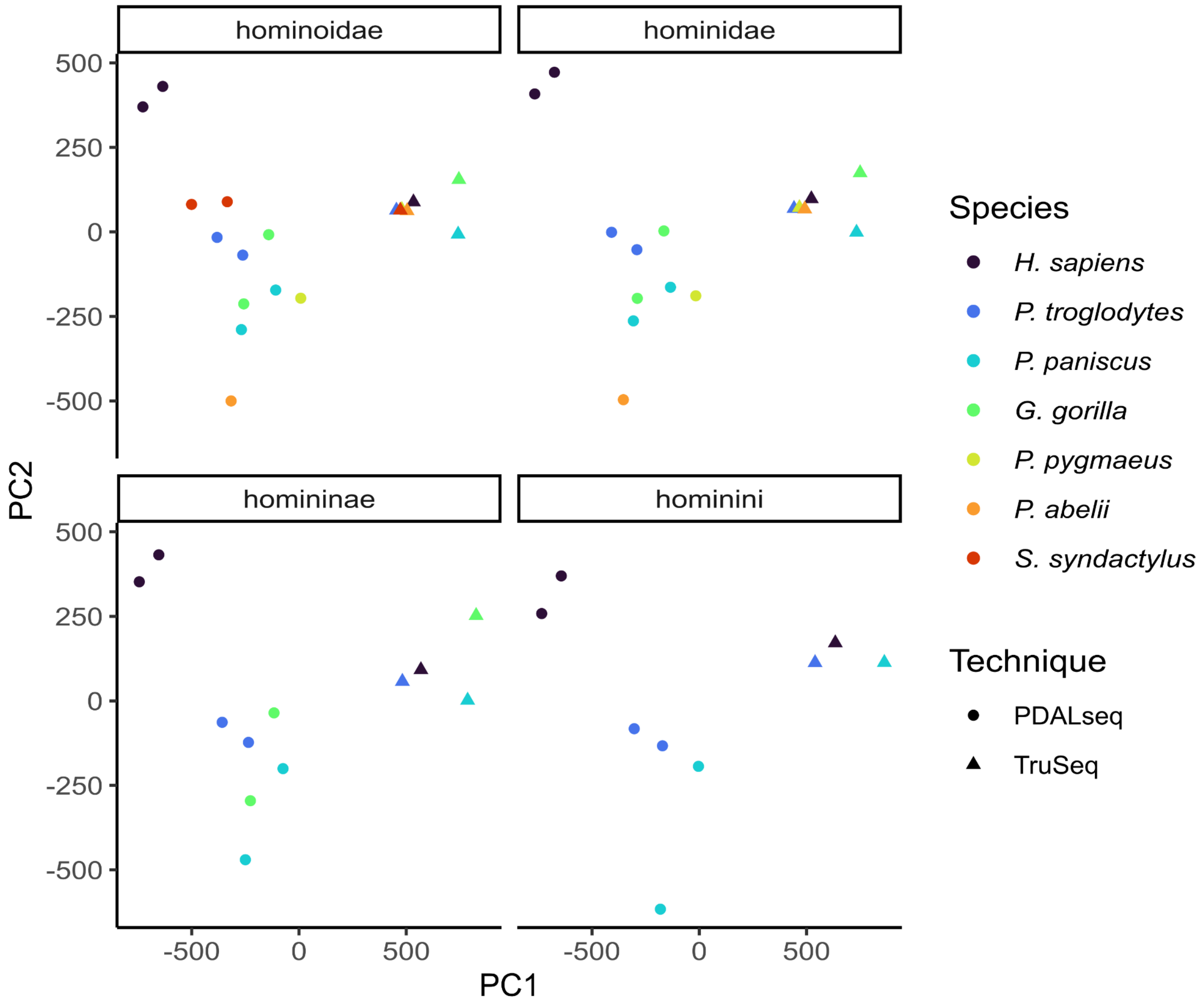

**Figure S7. PDAL-Seq signal in syntenic regions of the genome is correlated between ape species cell lines**

Spearman's rank correlation between read mapping levels in Hominoidae 8-way alignment blocks between different experiments (i.e., a cell line with either TruSeq or PDAL-Seq protocol applied to it). Fill corresponds to the correlation coefficient between the experiment on the x-axis and the experiment on the y-axis. The x-axis matches the y-axis, going bottom to top and left to right, respectively: GM24385-HG002 (human male), GM24143-HG004 (human female), AG18354 (chimpanzee male), AG18358 (chimpanzee female), KB8711 (bonobo male), KB14571 (bonobo female), KB37811 (gorilla male), F7549 (gorilla female), AG05252 (Bornean orangutan), AG06213 (Sumatran orangutan), Jambi (Siamang male), Karenina (Siamang female). The data are in Table S9.

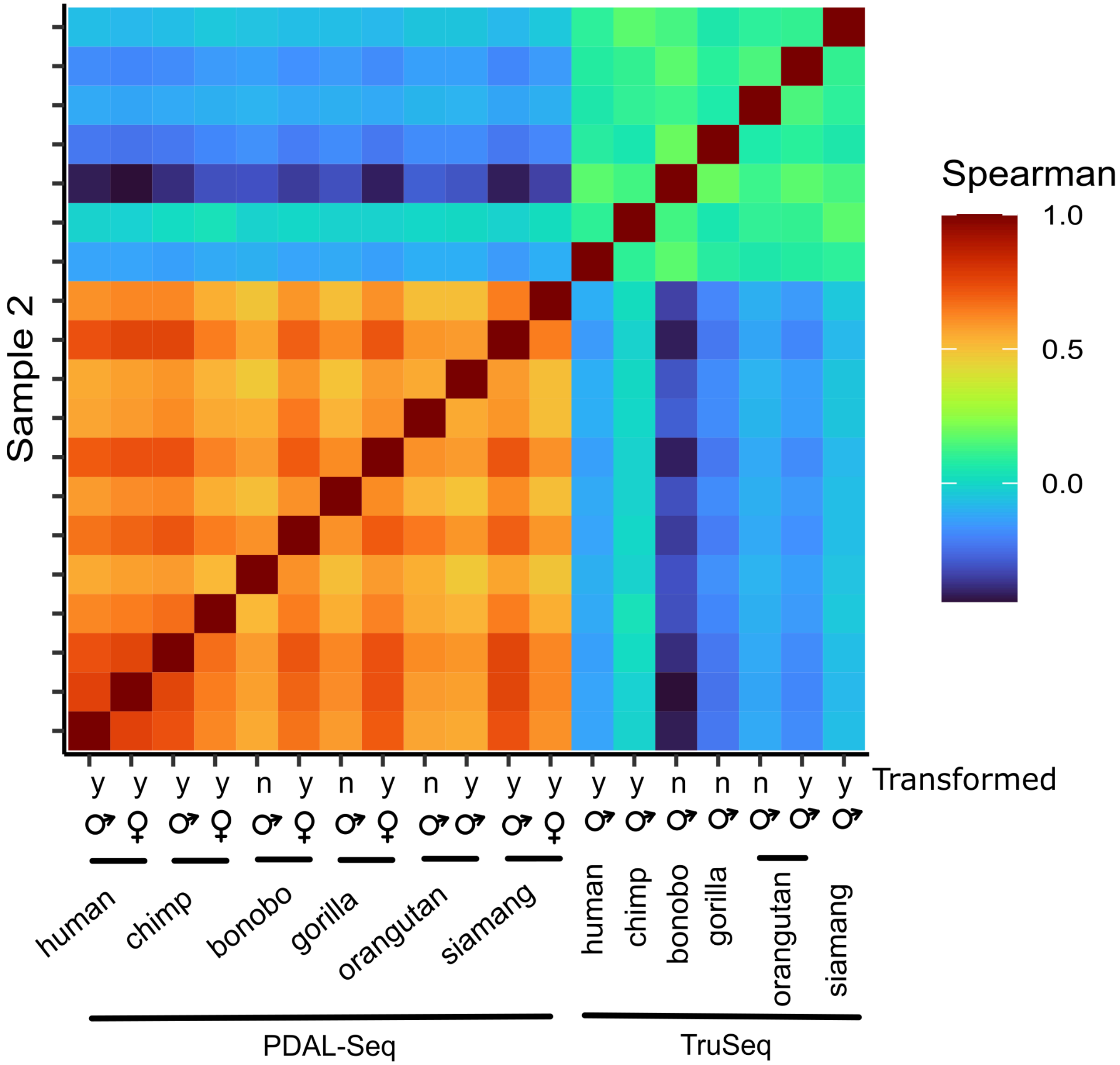

**Figure S8. Pairwise Spearman's rank correlation coefficients from Figure S8 for different types of comparisons**

*P*-values represent the result of a non-parametric Kruskal-Wallis test to determine the probability that the variance between groups could be explained by random sampling of the same population.

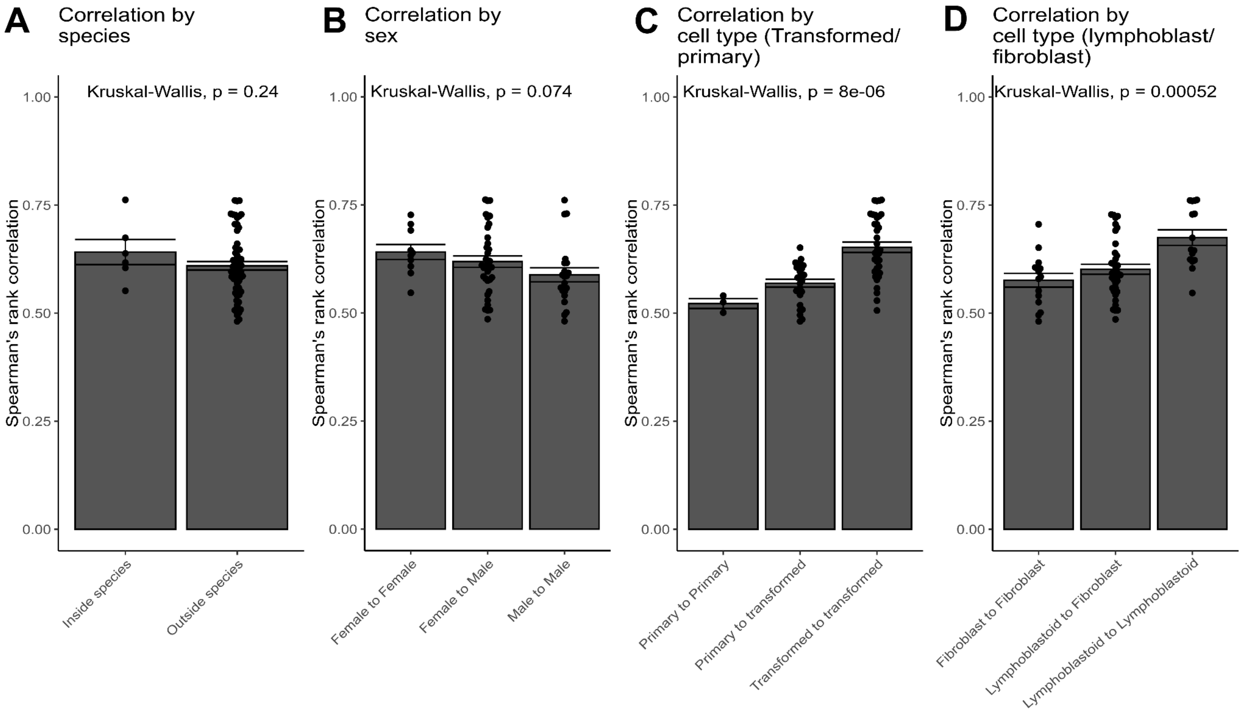

**Figure S9. Scree plots to determine state numbers for the analysis of PDAL-Seq data in ape alignments**

**(A)** PDAL-Seq data in alignments parsed using different species groups were clustered using a K-means clustering algorithm. **(B)** PDAL-Seq data in alignments parsed using different species groups were fit to a Multivariate Gaussian Hidden Markov Model and Bayesian Information Criterion (BIC) was determined for each fit.

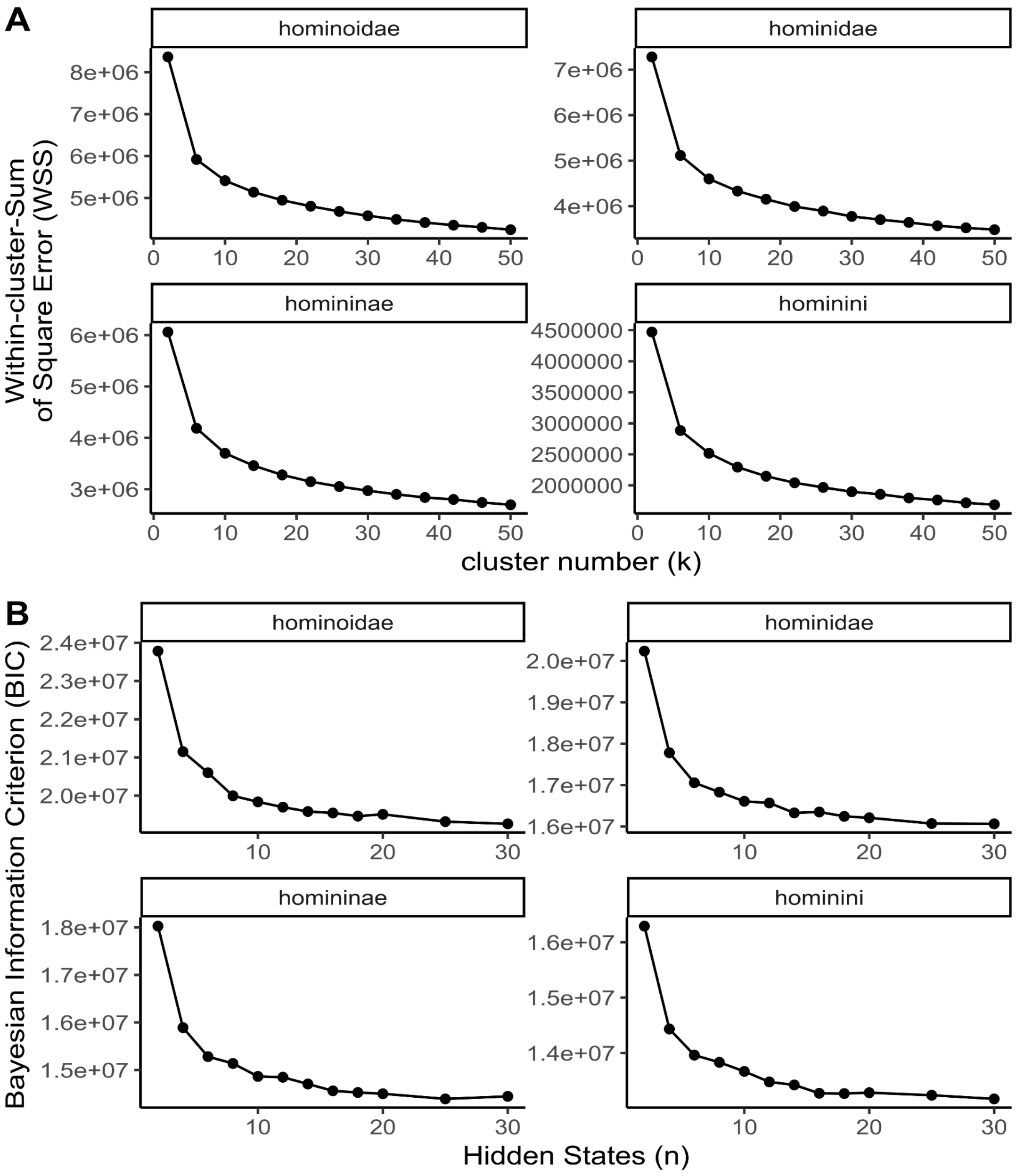

**Figure S10. Partitioning of variance as a function of hidden state number for Multivariate Gaussian Hidden Markov Model for the Hominoidae species group**

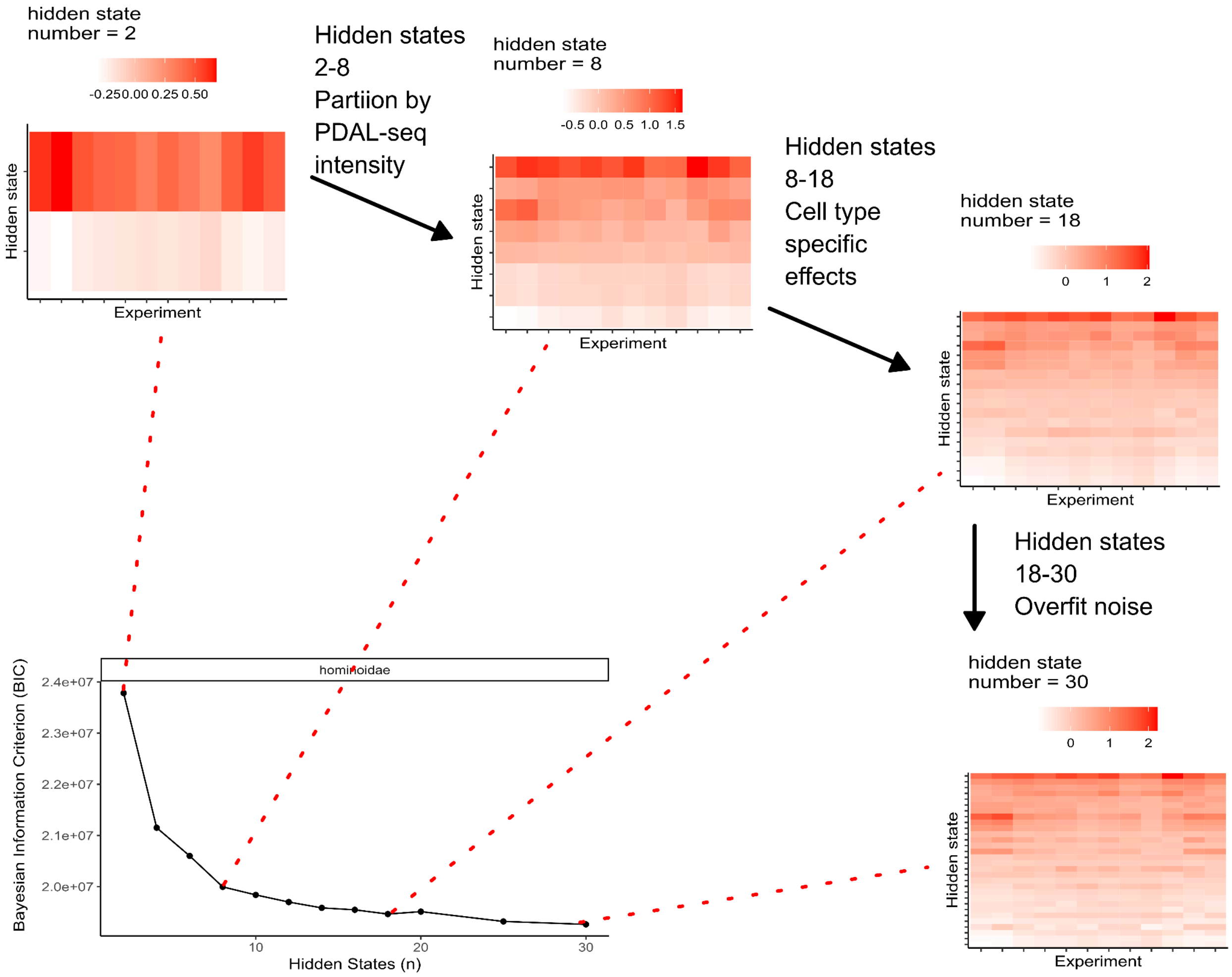

**Figure S11. Transition probability matrix for the 8-state Multivariate Gaussian Hidden Markov Model presented in Figure 3**

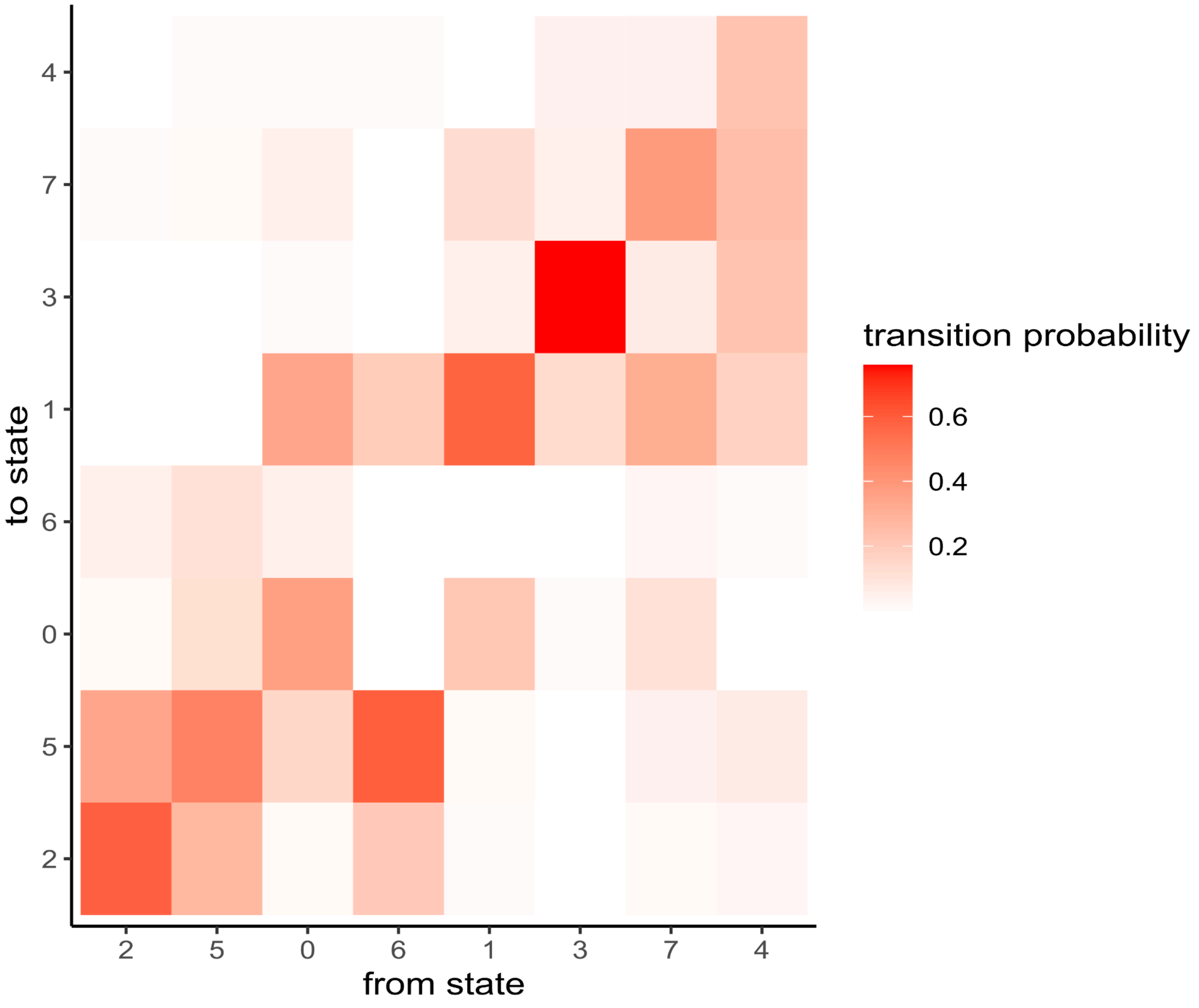

**Figure S12. Mean PDAL-Seq signal upstream and downstream of transcription start sites (TSSs)**
"All" contains all ~20,000 human TSSs. The remaining panels contain TSSs that overlap with ape PDAL-Seq states determined with MG-HMM presented in Figure 3.

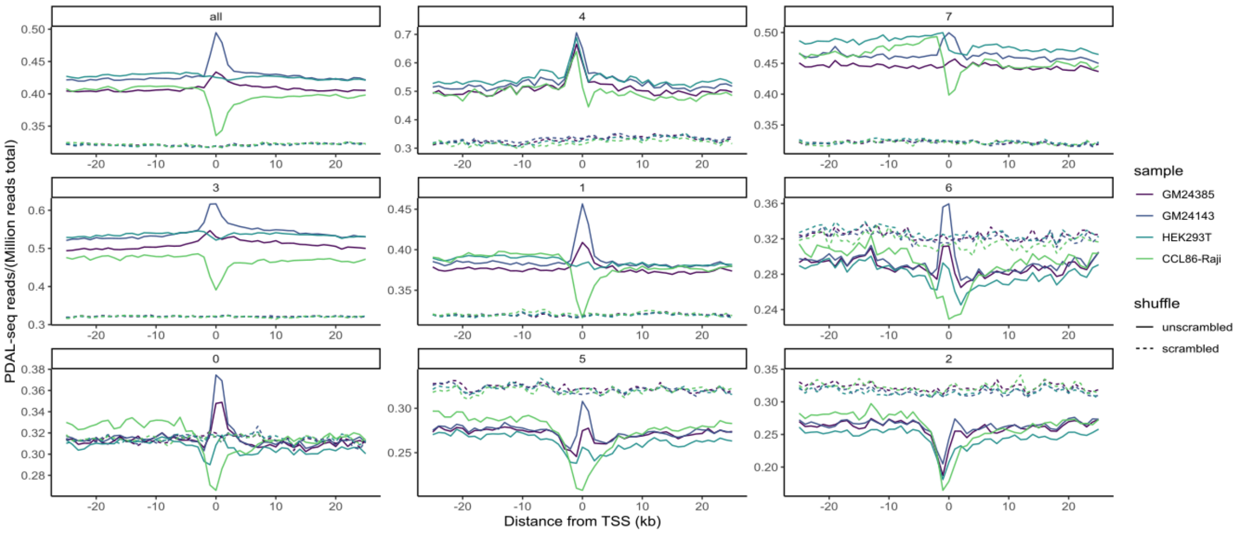

**Figure S13. Scree plots to determine state numbers for the analysis of PDAL-Seq data in the human genome**

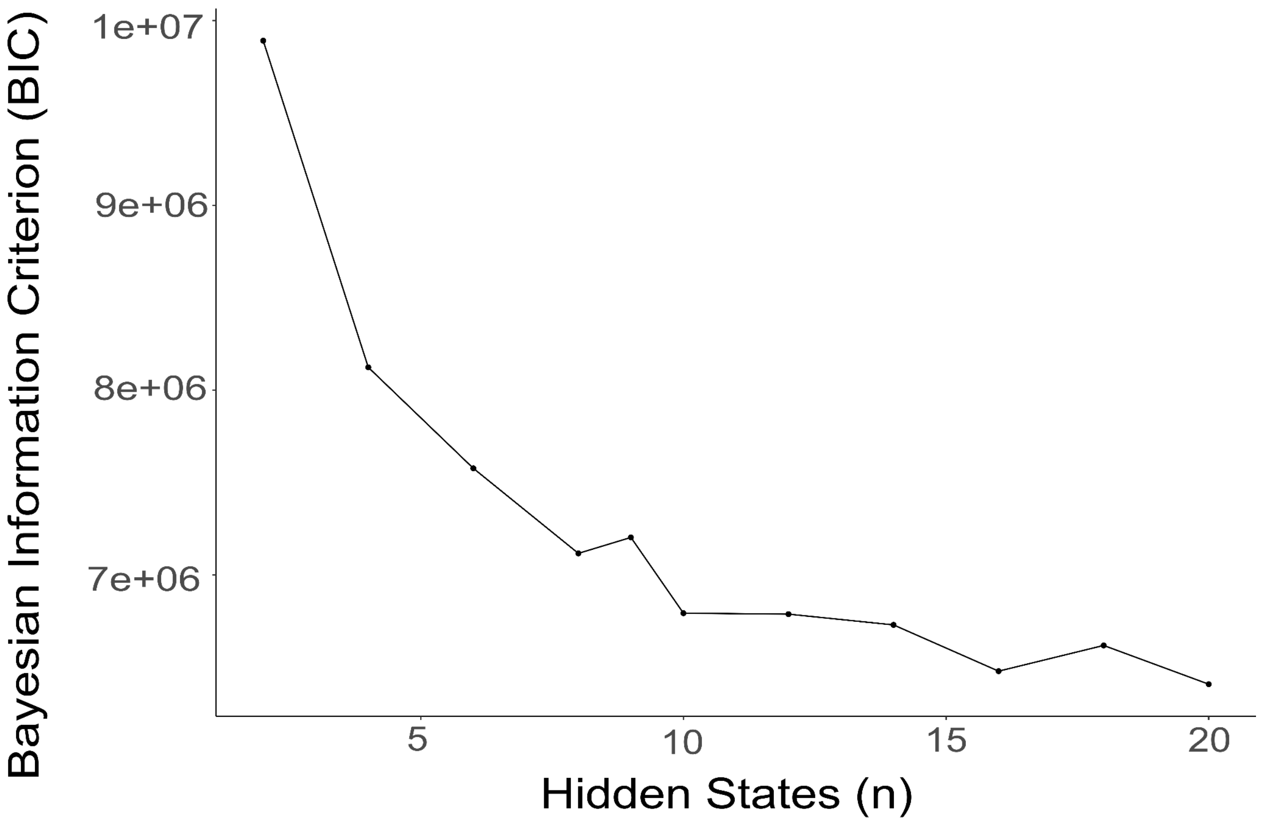

**Figure S14. Enrichment of repeat families in human PDAL-Seq MV-HMM models**
Enrichment corresponds to the observed number of annotations overlapping with a state divided by the number that would be observed by random chance. Repeat sub-families in this plot are members of the families in Figure 5D.

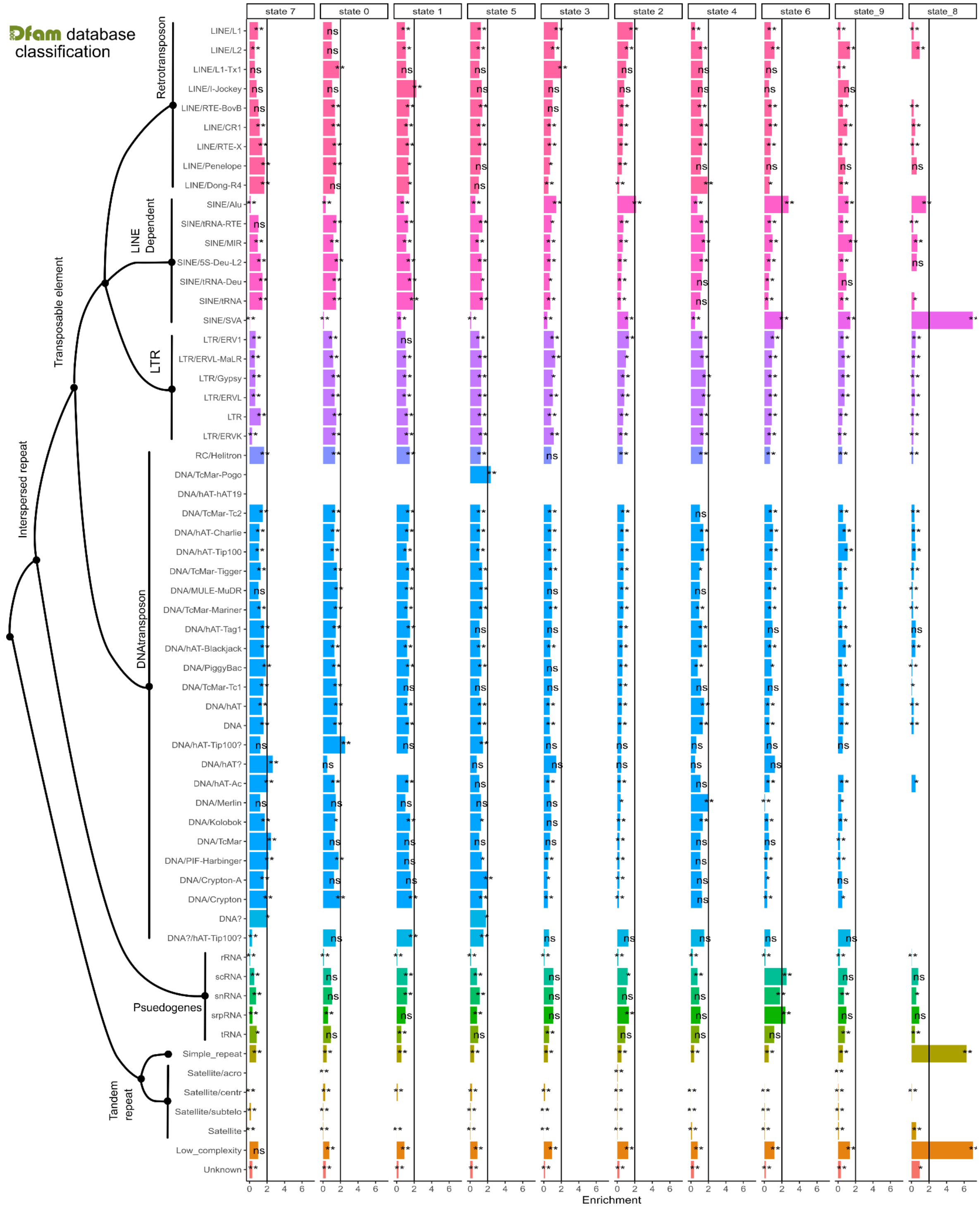

**Figure S15. Principal component analysis of PDAL-Seq reads in alignment blocks where each point represents a cell line x sequencing method (either PDAL-Seq or TruSeq) combination**

Alignments were parsed for the Hominoidae group.

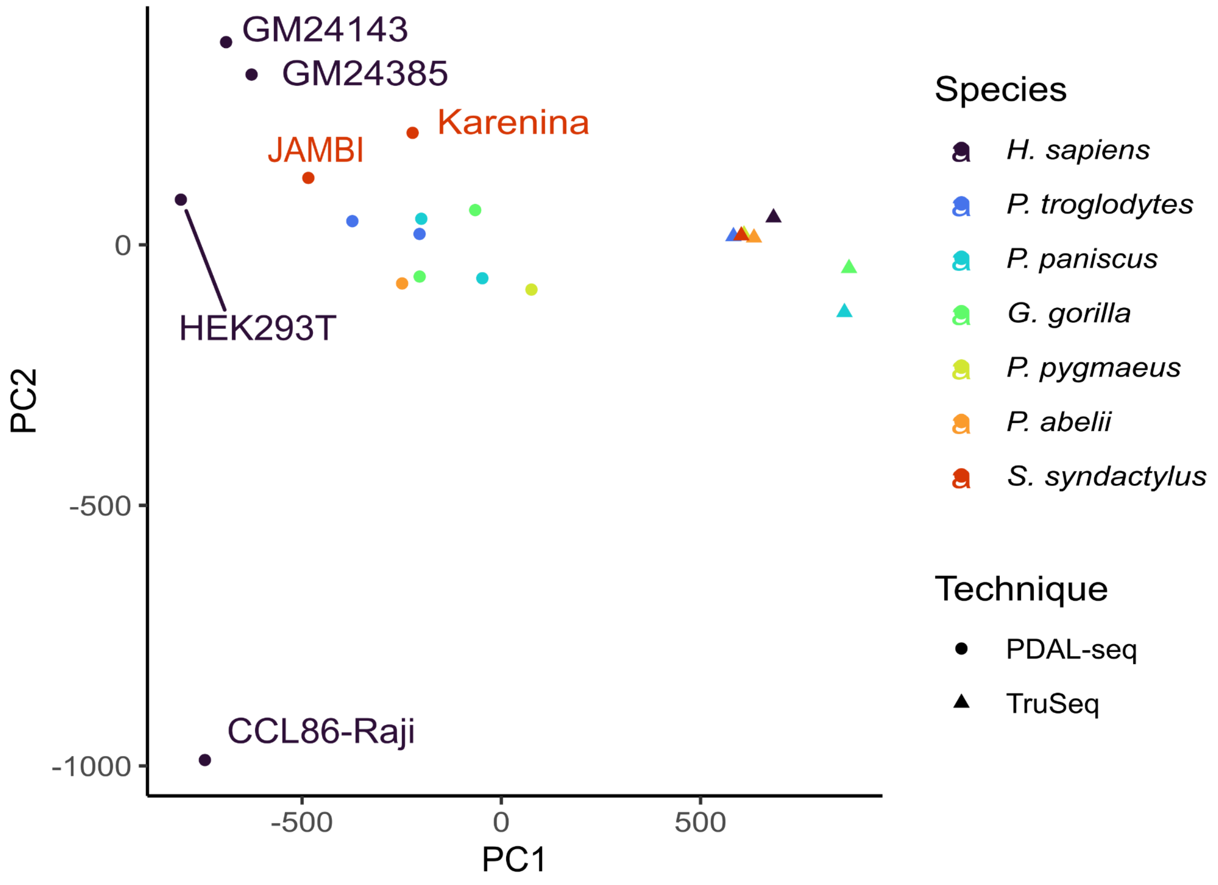

**Figure S16. Enrichment of PDAL-Seq reads in human functional genomic element annotations illustrates dysregulation of ssDNA distribution in the cancerous Raji cell line**
Enrichment is the number of PDAL-Seq reads observed in an annotation divided by the number of PDAL-Seq reads mapping to the same annotation that were randomly assigned to genomic loci. Significance levels: “ns” is $p \geq 0.05$, “*” is $p < 0.05$, and “**” is $p < 0.01$. P-values are determined by permutation testing the null hypothesis that the same number of reads would be observed by selecting random genomic loci ont the same chromosome with the same annotation length (see Methods).

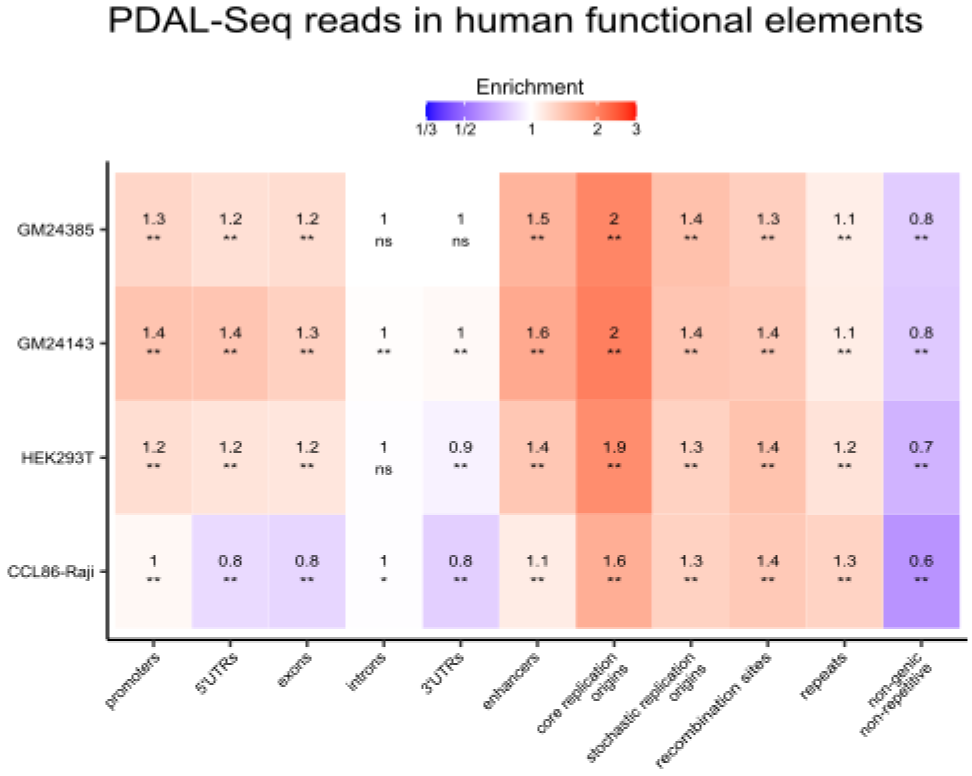

**Figure S17. Enrichment of PDAL-Seq and control TruSeq reads in ape satellite arrays.** Enrichment is the number of PDAL-Seq reads observed in an annotation divided by the number of PDAL-Seq reads mapping to the same annotation that were randomly assigned to genomic loci. Significance levels: "ns" is $p \geq 0.05$, "*" is $p < 0.05$, and "**" is $p < 0.01$. P-values are determined by permutation testing the null hypothesis that the same number of reads would be observed by selecting random genomic loci ont the same chromosome with the same annotation length (see Methods).

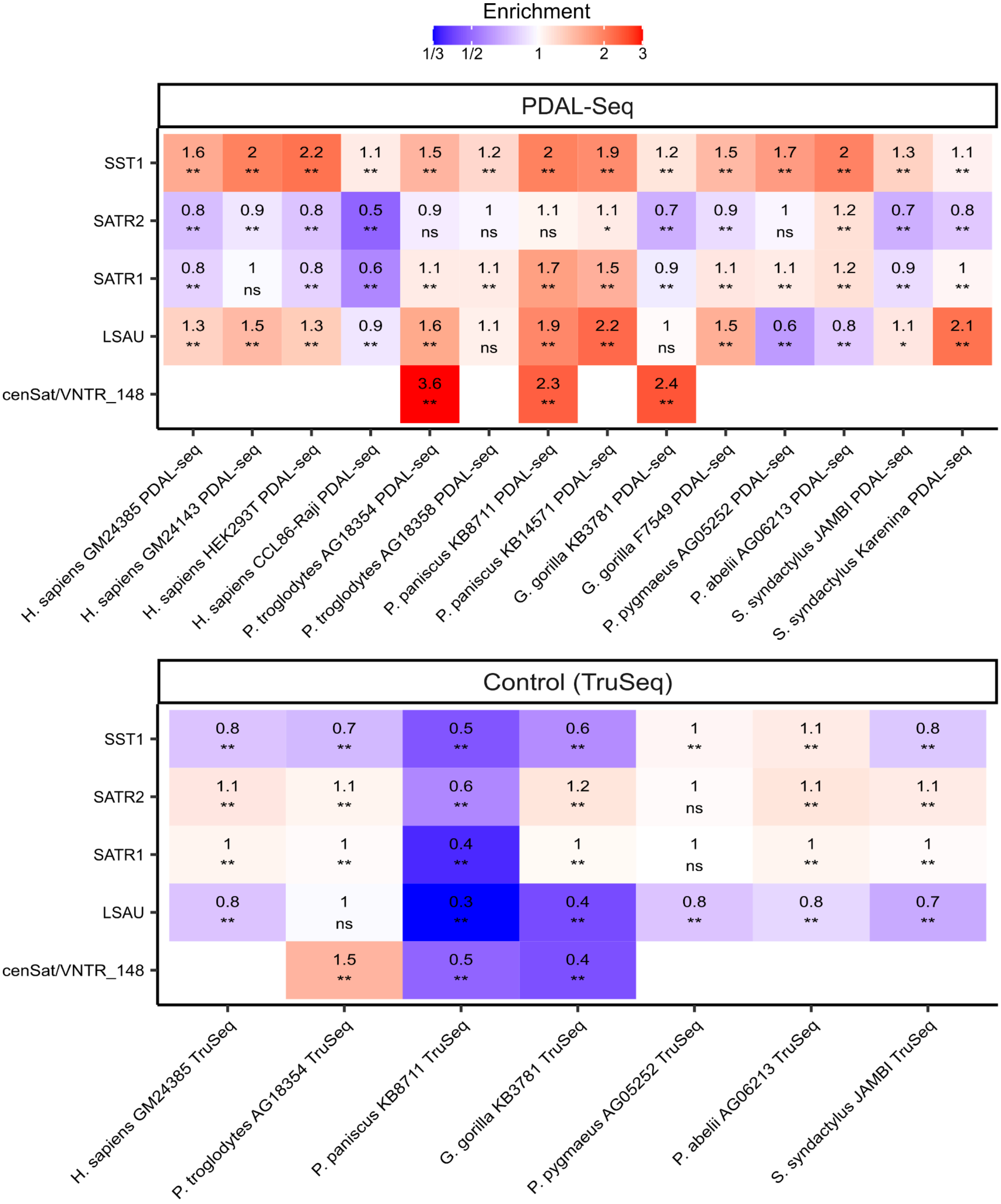


PDAL-Seq

| | H. sapiens GM24385 PDAL-seq | H. sapiens GM24143 PDAL-seq | H. sapiens HEK293T PDAL-seq | H. sapiens CCL86-Raji PDAL-seq | P. troglodytes AG18354 PDAL-seq | P. troglodytes AG18358 PDAL-seq | P. paniscus KB8711 PDAL-seq | P. paniscus KB14571 PDAL-seq | G. gorilla KB3781 PDAL-seq | G. gorilla F7549 PDAL-seq | P. pygmaeus AG05252 PDAL-seq | P. abelii AG06213 PDAL-seq | S. syndactylus JAMBI PDAL-seq | S. syndactylus Karenina PDAL-seq |
|---|---|---|---|---|---|---|---|---|---|---|---|---|---|---|
| SST1 | 1.6 ** | 2 ** | 2.2 ** | 1.1 ** | 1.5 ** | 1.2 ** | 2 ** | 1.9 ** | 1.2 ** | 1.5 ** | 1.7 ** | 2 ** | 1.3 ** | 1.1 ** |
| SATR2 | 0.8 ** | 0.9 ** | 0.8 ** | 0.5 ** | 0.9 ns | 1 ns | 1.1 ns | 1.1 * | 0.7 ** | 0.9 ** | 1 ns | 1.2 ** | 0.7 ** | 0.8 ** |
| SATR1 | 0.8 ** | 1 ns | 0.8 ** | 0.6 ** | 1.1 ** | 1.1 ** | 1.7 ** | 1.5 ** | 0.9 ** | 1.1 ** | 1.1 ** | 1.2 ** | 0.9 ** | 1 ** |
| LSAU | 1.3 ** | 1.5 ** | 1.3 ** | 0.9 ** | 1.6 ** | 1.1 ns | 1.9 ** | 2.2 ** | 1 ns | 1.5 ** | 0.6 ** | 0.8 ** | 1.1 * | 2.1 ** |
| cenSat/VNTR_148 | | | | | 3.6 ** | | 2.3 ** | | 2.4 ** | | | | | |

Control (TruSeq)

| | H. sapiens GM24385 TruSeq | P. troglodytes AG18354 TruSeq | P. paniscus KB8711 TruSeq | G. gorilla KB3781 TruSeq | P. pygmaeus AG05252 TruSeq | P. abelii AG06213 TruSeq | S. syndactylus JAMBI TruSeq |
|---|---|---|---|---|---|---|---|
| SST1 | 0.8 ** | 0.7 ** | 0.5 ** | 0.6 ** | 1 ** | 1.1 ** | 0.8 ** |
| SATR2 | 1.1 ** | 1.1 ** | 0.6 ** | 1.2 ** | 1 ns | 1.1 ** | 1.1 ** |
| SATR1 | 1 ** | 1 ** | 0.4 ** | 1 ** | 1 ns | 1 ** | 1 ** |
| LSAU | 0.8 ** | 1 ns | 0.3 ** | 0.4 ** | 0.8 ** | 0.8 ** | 0.7 ** |
| cenSat/VNTR_148 | | 1.5 ** | 0.5 ** | 0.4 ** | | | |

**Figure S18. Analysis of PDAL-Seq reads and non-B DNA motifs in Siamang subterminal alpha satellites**

**(A-B)** K-means clustering of PDAL-Seq reads, IR motifs, and G4 motifs in 1-Mbp genomic windows for two Siamang cell lines reveals genomic loci with abnormally high PDAL-Seq read density, high IR density, and low G4 density. **(C)** These genomic loci closely correspond to subterminal alpha-satellite annotations.

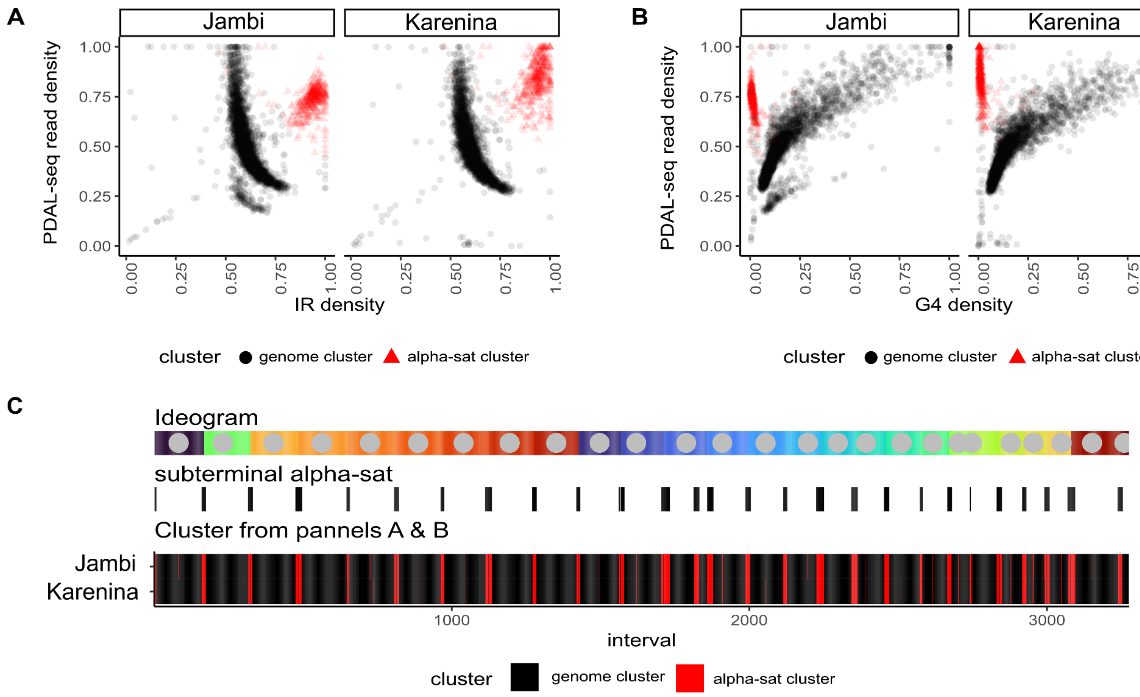

**Figure S19. Analysis of PDAL-Seq reads, DR motifs, and human Hsat2 and Hsat3 arrays in great ape chromosomes, organized by groups of homologous chromosomes**
Chromosomes are organized in homologous groups by human chromosomes, and data are shown in 1-Mbp windows. Top: the density of PDAL-Seq reads after a 99% winsorization normalization in primary male cell lines: *H. sapiens* GM24385, *P. troglodytes* AG18354*, P. paniscus* KB8711*, G. gorilla* KB37811*, P. pygmaeus* AG05252*, and P. abelii* AG06213. Middle:the coverage of DR repeats in a genomic window (bp divided by bp). Bottom: coverage by Hsat2,3 annotations from RepeatMasker.

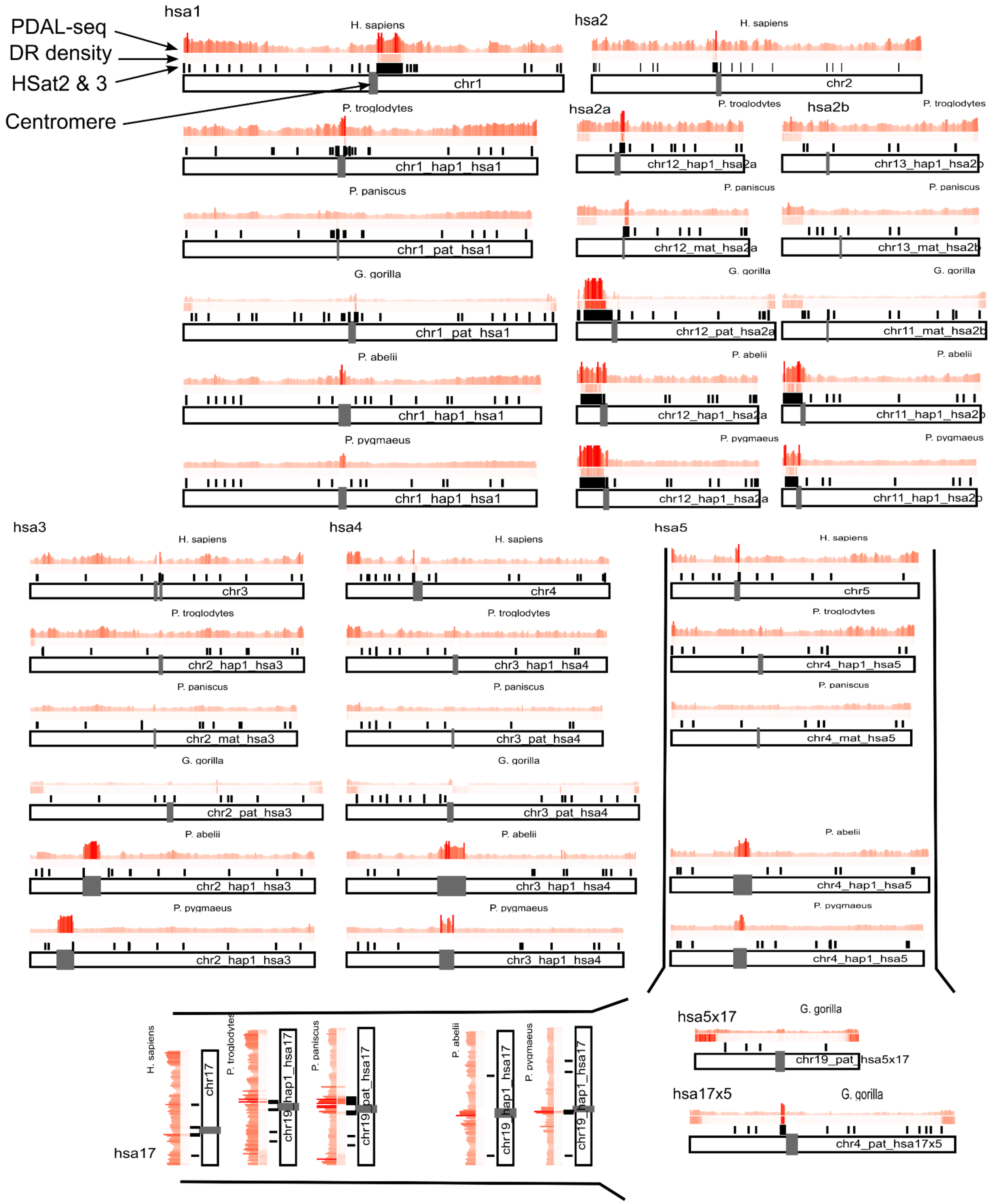
PDAL-seq
DR density
HSat2 & 3
Centromere
hsa1
hsa2
hsa2a
hsa2b
hsa3
hsa4
hsa5
hsa17
hsa5x17
hsa17x5
H. sapiens
P. troglodytes
P. paniscus
G. gorilla
P. abelii
P. pygmaeus
chr1
chr2
chr1_hap1_hsa1
chr1_pat_hsa1
chr12_hap1_hsa2a
chr13_hap1_hsa2b
chr12_mat_hsa2a
chr13_mat_hsa2b
chr12_pat_hsa2a
chr11_mat_hsa2b
chr11_hap1_hsa2b
chr3
chr4
chr5
chr2_hap1_hsa3
chr3_hap1_hsa4
chr4_hap1_hsa5
chr2_mat_hsa3
chr3_pat_hsa4
chr4_mat_hsa5
chr2_pat_hsa3
chr17
chr19_hap1_hsa17
chr19_pat_hsa17
chr19_pat_hsa5x17
chr4_pat_hsa17x5

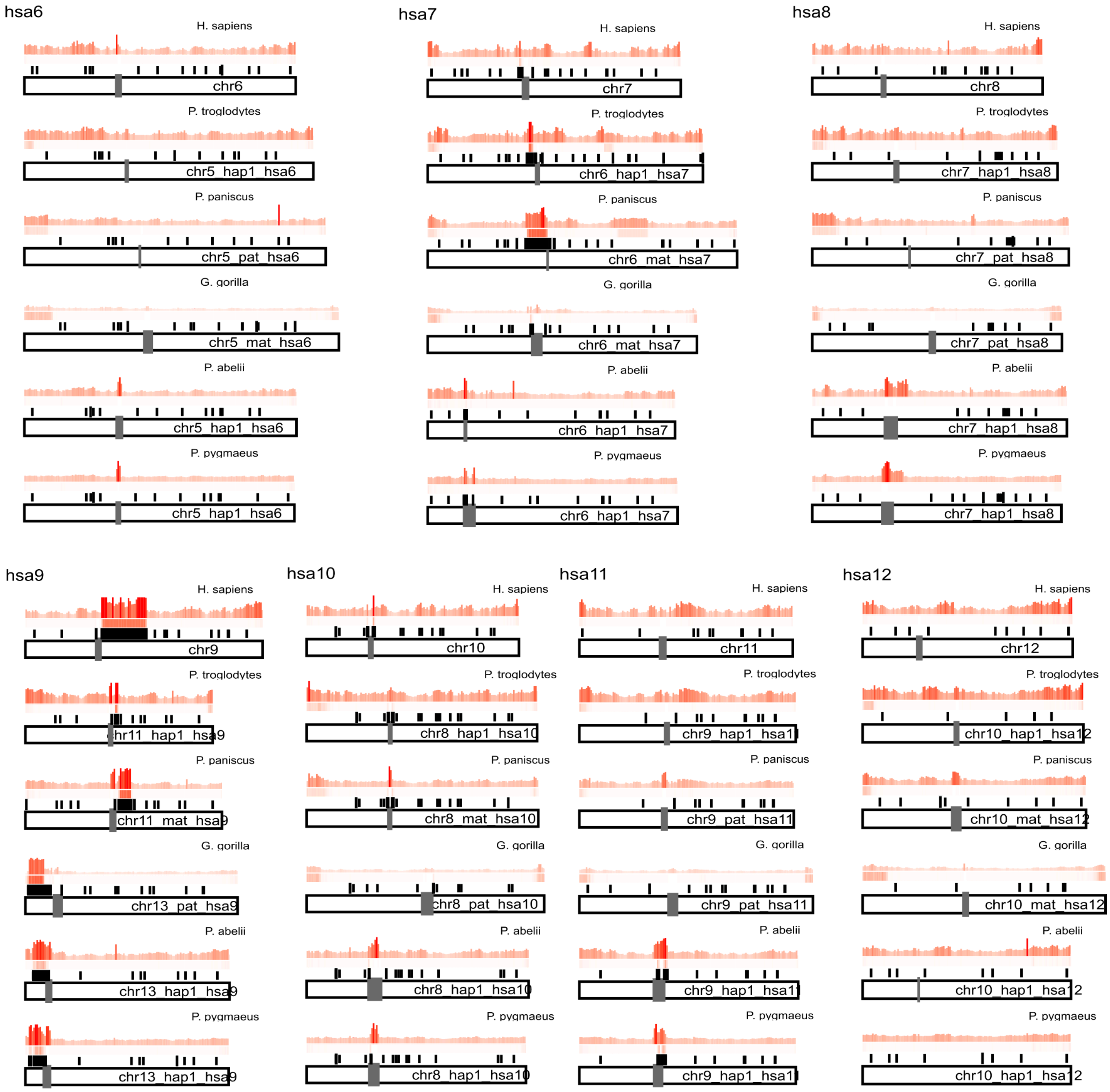
hsa6
H. sapiens
chr6
P. troglodytes
chr5 hap1 hsa6
P. paniscus
chr5 pat hsa6
G. gorilla
chr5 mat hsa6
P. abelii
chr5 hap1 hsa6
P. pygmaeus
chr5 hap1 hsa6
hsa7
H. sapiens
chr7
P. troglodytes
chr6 hap1 hsa7
P. paniscus
chr6 mat hsa7
G. gorilla
chr6 mat hsa7
P. abelii
chr6 hap1 hsa7
P. pygmaeus
chr6 hap1 hsa7
hsa8
H. sapiens
chr8
P. troglodytes
chr7 hap1 hsa8
P. paniscus
chr7 pat hsa8
G. gorilla
chr7 pat hsa8
P. abelii
chr7 hap1 hsa8
P. pygmaeus
chr7 hap1 hsa8
hsa9
H. sapiens
chr9
P. troglodytes
hr11 hap1 hsa9
P. paniscus
chr11 mat hsa9
G. gorilla
chr13 pat hsa9
P. abelii
chr13 hap1 hsa9
P. pygmaeus
chr13 hap1 hsa9
hsa10
H. sapiens
chr10
P. troglodytes
chr8 hap1 hsa10
P. paniscus
chr8 mat hsa10
G. gorilla
chr8 pat hsa10
P. abelii
chr8 hap1 hsa10
P. pygmaeus
chr8 hap1 hsa10
hsa11
H. sapiens
chr11
P. troglodytes
chr9 hap1 hsa11
P. paniscus
chr9 pat hsa11
G. gorilla
chr9 pat hsa11
P. abelii
chr9 hap1 hsa11
P. pygmaeus
chr9 hap1 hsa11
hsa12
H. sapiens
chr12
P. troglodytes
chr10 hap1 hsa12
P. paniscus
chr10 mat hsa12
G. gorilla
chr10 mat hsa12
P. abelii
chr10 hap1 hsa12
P. pygmaeus
chr10 hap1 hsa12

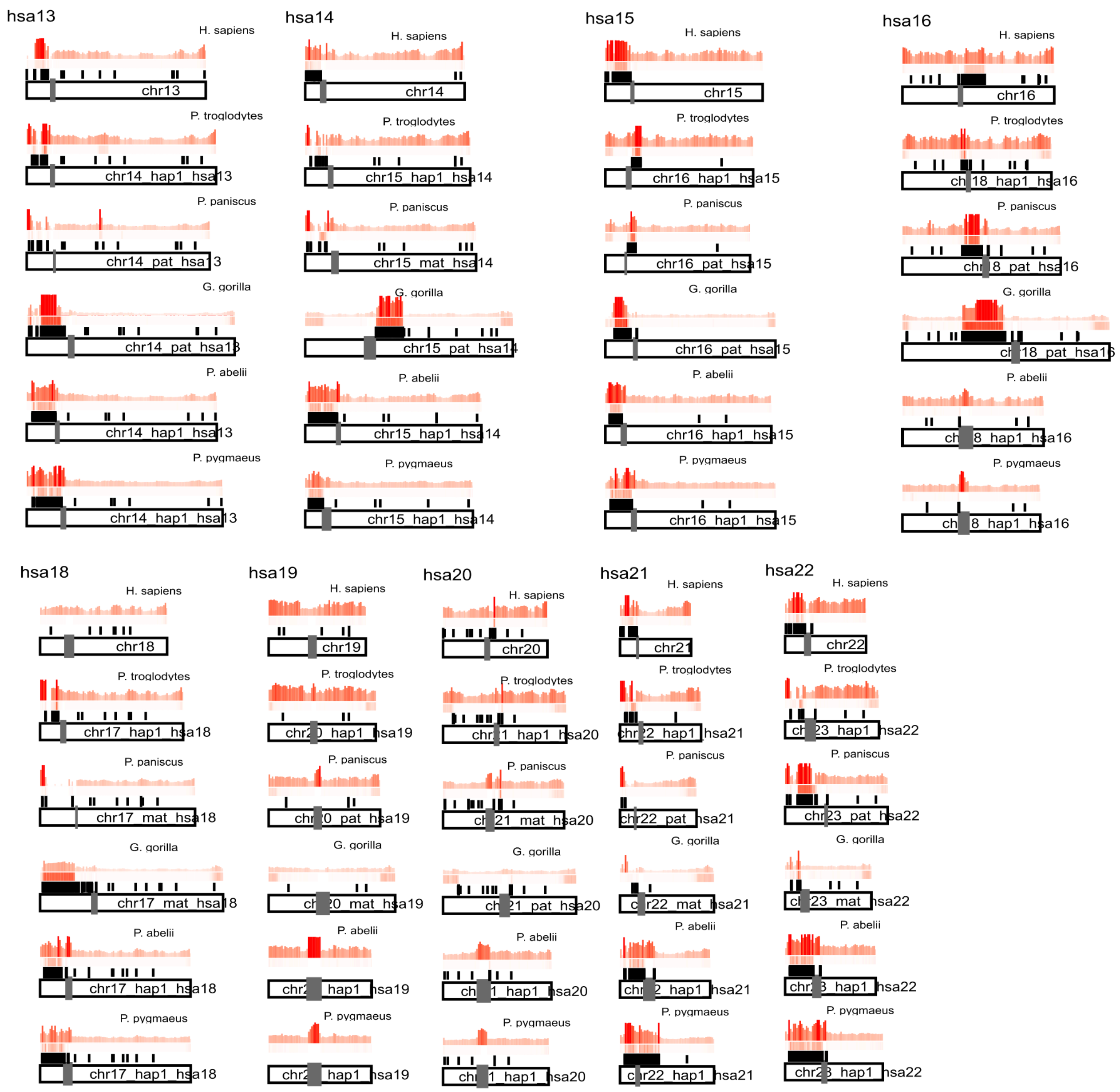
hsa13
H. sapiens
chr13
P. troglodytes
chr14_hap1_hsa13
P. paniscus
G. gorilla
P. abelii
chr14_hap1_hsa13
P. pygmaeus
hsa14
H. sapiens
chr14
P. troglodytes
chr15_hap1_hsa14
P. paniscus
G. gorilla
P. abelii
chr15_hap1_hsa14
P. pygmaeus
chr15_hap1_hsa14
hsa15
H. sapiens
chr15
P. troglodytes
chr16_hap1_hsa15
P. paniscus
chr16_pat_hsa15
G. gorilla
chr16_pat_hsa15
P. abelii
chr16_hap1_hsa15
P. pygmaeus
chr16_hap1_hsa15
hsa16
H. sapiens
chr16
P. troglodytes
P. paniscus
G. gorilla
P. abelii
P. pygmaeus
hsa18
H. sapiens
chr18
P. troglodytes
chr17_hap1_hsa18
P. paniscus
chr17_mat_hsa18
G. gorilla
chr17_mat_hsa18
P. abelii
chr17_hap1_hsa18
P. pygmaeus
chr17_hap1_hsa18
hsa19
H. sapiens
chr19
P. troglodytes
P. paniscus
G. gorilla
P. abelii
P. pygmaeus
hsa20
H. sapiens
chr20
P. troglodytes
P. paniscus
G. gorilla
P. abelii
P. pygmaeus
hsa21
H. sapiens
chr21
P. troglodytes
P. paniscus
G. gorilla
P. abelii
P. pygmaeus
hsa22
H. sapiens
chr22
P. troglodytes
P. paniscus
G. gorilla
P. abelii
P. pygmaeus

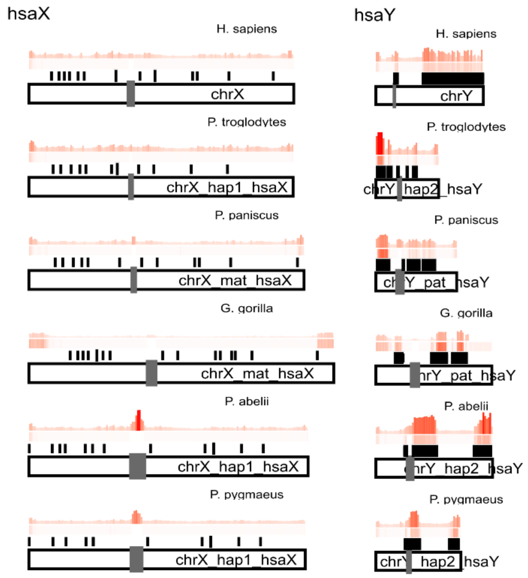
hsaX
H. sapiens
chrX
P. troglodytes
chrX_hap1_hsaX
P. paniscus
chrX_mat_hsaX
G. gorilla
chrX_mat_hsaX
P. abelii
chrX_hap1_hsaX
P. pygmaeus
chrX_hap1_hsaX
hsaY
H. sapiens
chrY
P. troglodytes
chrY_hap2_hsaY
P. paniscus
chrY_pat_hsaY
G. gorilla
chrY_pat_hsaY
P. abelii
chrY_hap2_hsaY
P. pygmaeus
chrY_hap2_hsaY

**Figure S20. Native PAGE and Size exclusion chromatography - multi-angle light scattering (SEC-MALS) analysis of Hsat3 $(AATGG)_n$ repeats *in vitro***

**(A)** Native PAGE gel analysis of $(AATGG)_n$ conformations. 67 pmol of AATGG monomer was separated on a 15% TBE PAGE gel supplemented with 10 mM KCL following renaturation in either 100 mM KCl 140 mM LiCl or 140 mM LiCl buffer. Bands were visualized with SYBR Gold and red pixels are oversaturated. **(B)** Multi-angle light scattering analysis (MALS) of $(AATGG)_n$ sequence peaks in the SEC-UV traces from Figure 6F. Molecular weights were estimated using MALS signal and either UV absorbance or refraction index as a concentration source. Relative molecular weights for each chromatogram were calculated by dividing the molecular weight of each peak by the molecular weight of the smallest peak.

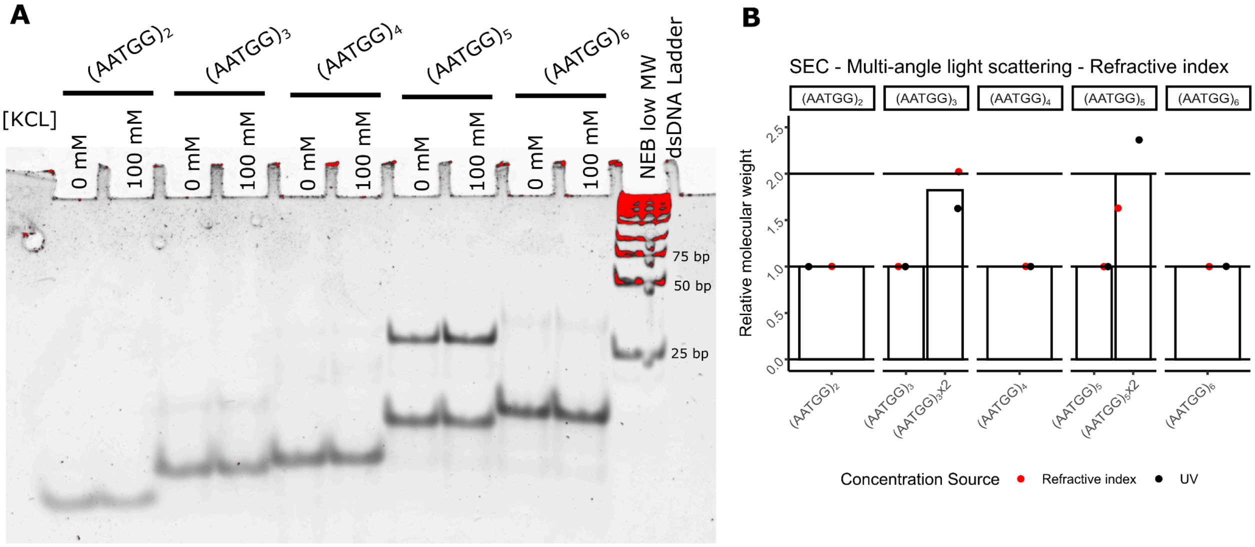


**Figure S21. An example of S1 digestion test gel**

Reaction products were run on a 0.6x TAE agarose gel and imaged using Gel Red fluorescence emission. The molecular weight ladder is dsDNA.

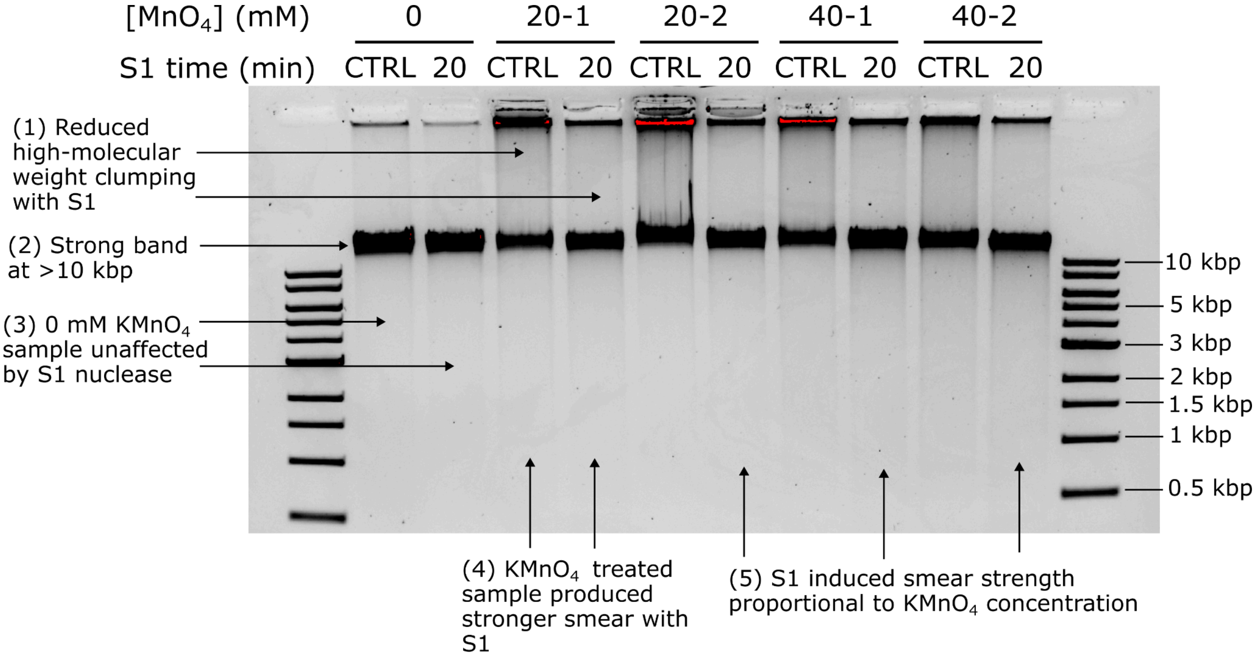
