## Supplemental tables for "Comparative analysis of single-stranded and non-canonical DNA formation in human and other ape cells with telomere-to-telomere genomes"

**Table S1: Summary statistics for mapping of PDAL-Seq and TruSeq sequencing reads to T2T genomes by technical replicate.**

Definitions:

Raw fastq reads: Number of reads in the original dataset

Trimmed fastq reads: Number of reads after read trimming by fastp

Primary mapped reads: Number of primary MAPPING locations. Note the secondary mapping locations for multi-mapping reads are not included in this dataset.

Supplementary mapped reads: Number of supplemental MAPPING locations due to chimeric alignments.

Uniquely mapped reads: Reads where the primary mapping location is 10x better than possible secondary mapping locations.

deduplicated: reads after removing PCR duplicates

high MAPQ: MAPQ >= 20 reads

| Species | Bio sample | Technique | Raw fastq reads | Trimmed fastq reads | Mapped primary reads | Mapped supplementary reads | Mapped deduplicated primary reads | Mapped deduplicated supplementary reads | Mapped deduplicated high MapQ primary reads | Mapped deduplicated high MapQ supplementary reads |
|---|---|---|---|---|---|---|---|---|---|---|
| H. sapiens | GM24385 | PDAL-Seq | 21143412 | 19178537 | 18857706 | 263566 | 12914825 | 185522 | 10662878 | 101616 |
| H. sapiens | GM24385 | PDAL-Seq | 22295005 | 20223469 | 19862004 | 391838 | 13421160 | 271275 | 11001354 | 147670 |
| H. sapiens | GM24385 | PDAL-Seq | 19200743 | 16742578 | 16393817 | 301457 | 11515671 | 216467 | 9390703 | 117690 |
| H. sapiens | GM24385 | PDAL-Seq | 26091710 | 23251804 | 22723458 | 440063 | 16006887 | 313811 | 12874576 | 174971 |
| H. sapiens | GM24143 | PDAL-Seq | 26036563 | 24875757 | 23598530 | 378052 | 17451439 | 287833 | 13741496 | 99810 |
| H. sapiens | GM24143 | PDAL-Seq | 23630188 | 23053180 | 22205100 | 124912 | 17454879 | 100739 | 14954775 | 37737 |
| H. sapiens | GM24143 | PDAL-Seq | 21530059 | 20741308 | 19828434 | 327950 | 14868602 | 251925 | 11706870 | 82214 |
| H. sapiens | GM24143 | PDAL-Seq | 28878696 | 27854333 | 26814903 | 191057 | 18981465 | 142603 | 16545905 | 49301 |
| H. sapiens | HEK293T | PDAL-Seq | 107189356 | 102227687 | 95765582 | 1536302 | 56841969 | 907571 | 40260198 | 323400 |
| H. sapiens | HEK293T | PDAL-Seq | 87523707 | 84338117 | 81509214 | 1378030 | 54883162 | 937251 | 41051961 | 403940 |
| H. sapiens | HEK293T | PDAL-Seq | 105703510 | 101047401 | 96357344 | 1569860 | 63150478 | 1032740 | 47745404 | 482336 |
| H. sapiens | HEK293T | PDAL-Seq | 113042174 | 107016574 | 102853182 | 927496 | 50791464 | 447546 | 38900399 | 164336 |
| H. sapiens | CCL86-Raji | PDAL-Seq | 9969066 | 9937546 | 9883949 | 1081447 | 7579308 | 838201 | 5891328 | 425432 |
| H. sapiens | CCL86-Raji | PDAL-Seq | 9527744 | 9497599 | 9443567 | 703252 | 7324869 | 555142 | 5830927 | 260794 |
| H. sapiens | CCL86-Raji | PDAL-Seq | 10536814 | 10496590 | 10448056 | 1020258 | 7706635 | 768802 | 6059076 | 373451 |
| H. sapiens | CCL86-Raji | PDAL-Seq | 11740482 | 11702043 | 11628992 | 1090931 | 8685133 | 825059 | 6857703 | 429392 |
| H. sapiens | CCL86-Raji | PDAL-Seq | 10986001 | 10958158 | 10885409 | 591445 | 8389369 | 460847 | 6620062 | 233541 |
| H. sapiens | CCL86-Raji | PDAL-Seq | 11487785 | 11456886 | 11368412 | 641780 | 8682264 | 497649 | 7025557 | 259413 |
| H. sapiens | CCL86-Raji | PDAL-Seq | 10578563 | 10532151 | 10487763 | 1088945 | 7613449 | 809580 | 6222376 | 407381 |
| H. sapiens | CCL86-Raji | PDAL-Seq | 9489844 | 9447884 | 9395726 | 810976 | 7157253 | 630282 | 5772489 | 307142 |

| | | | | | | | | | | |
|---|---|---|---|---|---|---|---|---|---|---|
| P. troglodytes | AG18354 | PDAL-Seq | 25424981 | 22470661 | 21316772 | 552009 | 15096685 | 394225 | 10869686 | 148651 |
| P. troglodytes | AG18354 | PDAL-Seq | 30327651 | 26278696 | 24765532 | 298226 | 17287689 | 211935 | 12803345 | 69503 |
| P. troglodytes | AG18354 | PDAL-Seq | 32664407 | 28450425 | 26477094 | 296395 | 19333953 | 220416 | 13952530 | 55317 |
| P. troglodytes | AG18354 | PDAL-Seq | 31560018 | 29662604 | 29306662 | 428568 | 20208716 | 299206 | 15262229 | 145947 |
| P. troglodytes | AG18354 | PDAL-Seq | 31632412 | 30435541 | 30281476 | 119181 | 19482161 | 81291 | 15308000 | 28585 |
| P. troglodytes | AG18354 | PDAL-Seq | 29083369 | 29033038 | 26187915 | 153649 | 18203114 | 110224 | 14375363 | 43166 |
| P. troglodytes | AG18354 | PDAL-Seq | 31814352 | 29959513 | 29707630 | 172157 | 20218972 | 121008 | 16031099 | 50497 |
| P. troglodytes | AG18358 | PDAL-Seq | 23808807 | 21607206 | 21224797 | 484297 | 15290783 | 354700 | 12072210 | 172420 |
| P. troglodytes | AG18358 | PDAL-Seq | 15442159 | 14008410 | 13841350 | 99306 | 10267980 | 77070 | 8511049 | 25792 |
| P. troglodytes | AG18358 | PDAL-Seq | 27584909 | 26178606 | 25954038 | 195069 | 18459181 | 144779 | 15448178 | 56908 |
| P. troglodytes | AG18358 | PDAL-Seq | 29170130 | 27917593 | 27766130 | 208178 | 18864940 | 149269 | 15678498 | 58085 |
| P. paniscus | KB8711 | PDAL-Seq | 26385672 | 26045302 | 25974051 | 124128 | 17219650 | 95062 | 13437053 | 18801 |
| P. paniscus | KB8711 | PDAL-Seq | 21299434 | 21250503 | 20975919 | 99659 | 13982952 | 76406 | 10916244 | 15450 |
| P. paniscus | KB8711 | PDAL-Seq | 22996246 | 22941573 | 22684745 | 118004 | 9325246 | 72213 | 6278924 | 9885 |
| P. paniscus | KB8711 | PDAL-Seq | 24647918 | 24428522 | 24380180 | 122353 | 12582167 | 85142 | 8970726 | 12632 |
| P. paniscus | KB14571 | PDAL-Seq | 92205907 | 91949428 | 89233535 | 585869 | 56693779 | 419141 | 42863823 | 62294 |
| P. paniscus | KB14571 | PDAL-Seq | 86425026 | 86193786 | 83106092 | 486989 | 51933341 | 343824 | 39418823 | 57649 |
| P. paniscus | KB14571 | PDAL-Seq | 84259514 | 84022498 | 80995420 | 599793 | 49887054 | 419827 | 37379728 | 63603 |
| P. paniscus | KB14571 | PDAL-Seq | 82140387 | 81903883 | 78575865 | 516841 | 51972802 | 378535 | 39532620 | 59266 |
| P. paniscus | KB14571 | PDAL-Seq | 101644173 | 101355821 | 94094603 | 735746 | 58836607 | 514065 | 43649696 | 84474 |
| G. gorilla | KB3781 | PDAL-Seq | 18143566 | 18110717 | 17820671 | 55494 | 13765851 | 46461 | 9856626 | 10113 |
| G. gorilla | KB3781 | PDAL-Seq | 31453015 | 31339307 | 25899828 | 235562 | 23360462 | 197187 | 13324284 | 23834 |
| G. gorilla | KB3781 | PDAL-Seq | 35395196 | 35263285 | 30468745 | 446059 | 27143378 | 386495 | 15112740 | 54805 |
| G. gorilla | KB3781 | PDAL-Seq | 22309952 | 22234577 | 18441980 | 265044 | 16382692 | 228874 | 8667243 | 23780 |
| G. gorilla | F7549 | PDAL-Seq | 64233173 | 55872457 | 54727553 | 369713 | 45095923 | 302156 | 31475628 | 82605 |
| G. gorilla | F7549 | PDAL-Seq | 76997291 | 66469818 | 65208637 | 325983 | 52375091 | 261695 | 36008256 | 39041 |
| G. gorilla | F7549 | PDAL-Seq | 50476552 | 50354366 | 48764160 | 209109 | 37958761 | 167987 | 26892334 | 25366 |
| G. gorilla | F7549 | PDAL-Seq | 76055255 | 63334907 | 61385292 | 550839 | 48742702 | 440866 | 29755993 | 49159 |
| G. gorilla | F7549 | PDAL-Seq | 68941177 | 68778864 | 56515211 | 513727 | 45686785 | 411284 | 28257854 | 66313 |

| | | | | | | | | | | |
|---|---|---|---|---|---|---|---|---|---|---|
| P. pygmaeus | AG05252 | PDAL-Seq | 117731524 | 106832385 | 105885766 | 487898 | 66717809 | 342826 | 50335561 | 65454 |
| P. pygmaeus | AG05252 | PDAL-Seq | 125288428 | 114002928 | 113315029 | 601322 | 55188665 | 365184 | 41083546 | 80756 |
| P. pygmaeus | AG05252 | PDAL-Seq | 110781905 | 110285690 | 107510091 | 628230 | 50091618 | 379830 | 35586618 | 70837 |
| P. pygmaeus | AG05252 | PDAL-Seq | 118480934 | 117949100 | 112317592 | 639990 | 55029137 | 399924 | 40325716 | 71829 |
| P. abelii | AG06213 | PDAL-Seq | 41015157 | 36015132 | 33606499 | 109730 | 21706844 | 75095 | 16272833 | 12775 |
| P. abelii | AG06213 | PDAL-Seq | 8909181 | 8046635 | 7570980 | 30506 | 5115661 | 22199 | 3723623 | 3848 |
| P. abelii | AG06213 | PDAL-Seq | 23612687 | 21245802 | 19815128 | 110063 | 12858224 | 77063 | 9180989 | 14203 |
| P. abelii | AG06213 | PDAL-Seq | 22205741 | 20154658 | 19346121 | 105805 | 12880376 | 76228 | 9060168 | 13198 |
| S. syndactylus | JAMBI | PDAL-Seq | 34620783 | 29564628 | 26979809 | 69384 | 19883648 | 50536 | 13649454 | 16226 |
| S. syndactylus | JAMBI | PDAL-Seq | 16457669 | 14324008 | 13016753 | 46920 | 9929683 | 34976 | 6485569 | 11599 |
| S. syndactylus | JAMBI | PDAL-Seq | 31923753 | 27609022 | 25361035 | 121526 | 18778009 | 87111 | 12954467 | 39673 |
| S. syndactylus | JAMBI | PDAL-Seq | 38565391 | 32947291 | 30236492 | 501852 | 27199935 | 454347 | 19483901 | 120315 |
| S. syndactylus | JAMBI | PDAL-Seq | 39301447 | 34264123 | 30546764 | 543603 | 27309458 | 489011 | 19556418 | 150788 |
| S. syndactylus | JAMBI | PDAL-Seq | 36699383 | 31406906 | 28286501 | 435575 | 25411487 | 395656 | 17430129 | 102112 |
| S. syndactylus | Karenina | PDAL-Seq | 21434356 | 20397064 | 20062748 | 176723 | 13482731 | 124165 | 9739514 | 36710 |
| S. syndactylus | Karenina | PDAL-Seq | 2682163 | 2525001 | 2479528 | 18093 | 1667329 | 12939 | 1191396 | 3598 |
| S. syndactylus | Karenina | PDAL-Seq | 23108285 | 21332548 | 20882836 | 125466 | 14209183 | 89884 | 10256751 | 25143 |
| S. syndactylus | Karenina | PDAL-Seq | 35673605 | 33332290 | 32716286 | 178781 | 21860861 | 126662 | 15956718 | 35198 |
| H. sapiens | GM24385 | TruSeq | 35124510 | 34159459 | 34069971 | 92284 | 33523173 | 80315 | 30197583 | 23543 |
| H. sapiens | GM24385 | TruSeq | 33487605 | 32550953 | 32463369 | 86050 | 31867292 | 76150 | 28671727 | 22846 |
| H. sapiens | GM24385 | TruSeq | 34512030 | 33661528 | 33624808 | 65960 | 33072267 | 56050 | 29853430 | 11851 |
| P. troglodytes | AG18354 | TruSeq | 48644096 | 48469293 | 48260468 | 47263 | 45445333 | 40315 | 39869169 | 7623 |
| P. troglodytes | AG18354 | TruSeq | 45924352 | 45736460 | 45542305 | 42523 | 44008070 | 36737 | 38678401 | 6864 |
| P. troglodytes | AG18354 | TruSeq | 47390720 | 47215883 | 47009720 | 44616 | 44838281 | 36733 | 39382400 | 7772 |
| P. troglodytes | AG18354 | TruSeq | 40517632 | 40297668 | 40117805 | 39569 | 38949055 | 35045 | 34141700 | 5886 |
| P. troglodytes | AG18354 | TruSeq | 44015616 | 43817147 | 43627852 | 39618 | 41970869 | 34146 | 36817181 | 6106 |
| P. troglodytes | AG18354 | TruSeq | 42369024 | 42162749 | 41982232 | 38458 | 40496523 | 33538 | 35505229 | 6047 |
| P. troglodytes | AG18354 | TruSeq | 45293568 | 45064930 | 44887425 | 43273 | 40307381 | 36314 | 35170692 | 6203 |
| P. troglodytes | AG18354 | TruSeq | 45727744 | 45500685 | 45323334 | 43268 | 40571369 | 36252 | 35401785 | 6474 |

| | | | | | | | | | | |
|---|---|---|---|---|---|---|---|---|---|---|
| P. paniscus | KB8711 | TruSeq | 319637420 | 318961109 | 315356545 | 2057799 | 303673461 | 1768988 | 266874988 | 370996 |
| G. gorilla | KB3781 | TruSeq | 35086293 | 34278528 | 34262989 | 42345 | 33618059 | 37999 | 26922096 | 4819 |
| P. pygmaeus | AG05252 | TruSeq | 46874624 | 46718503 | 46692103 | 48799 | 44003321 | 42079 | 38491523 | 8474 |
| P. pygmaeus | AG05252 | TruSeq | 43933696 | 43770441 | 43739115 | 42686 | 42323686 | 37307 | 37132907 | 7778 |
| P. pygmaeus | AG05252 | TruSeq | 45465600 | 45312657 | 45284571 | 45474 | 43273433 | 38722 | 37924895 | 8672 |
| P. pygmaeus | AG05252 | TruSeq | 37953536 | 37770814 | 37724764 | 37444 | 36726430 | 33415 | 32219572 | 6625 |
| P. pygmaeus | AG05252 | TruSeq | 41181184 | 41017036 | 40981183 | 38419 | 39528472 | 33543 | 34665385 | 7210 |
| P. pygmaeus | AG05252 | TruSeq | 39632896 | 39462627 | 39423107 | 37091 | 38149329 | 32509 | 33470090 | 6829 |
| P. pygmaeus | AG05252 | TruSeq | 43548672 | 43346917 | 43309940 | 43521 | 39257261 | 36923 | 34250251 | 7181 |
| P. pygmaeus | AG05252 | TruSeq | 44294144 | 44091664 | 44055629 | 44836 | 39805239 | 37790 | 34724787 | 7642 |
| P. abelii | AG06213 | TruSeq | 45670400 | 45514892 | 41064699 | 50423 | 38278506 | 41948 | 33067636 | 7985 |
| P. abelii | AG06213 | TruSeq | 43565056 | 43396579 | 39469497 | 44785 | 37905300 | 37932 | 32849185 | 7237 |
| P. abelii | AG06213 | TruSeq | 44507136 | 44352201 | 40137637 | 47228 | 38055799 | 38498 | 32943232 | 8275 |
| P. abelii | AG06213 | TruSeq | 37273600 | 37077248 | 34084963 | 38759 | 32970804 | 33468 | 28577235 | 6252 |
| P. abelii | AG06213 | TruSeq | 41058304 | 40880765 | 37355224 | 40843 | 35757061 | 34499 | 30981626 | 6707 |
| P. abelii | AG06213 | TruSeq | 39321600 | 39140539 | 35915820 | 39538 | 34473998 | 33532 | 29873756 | 6522 |
| P. abelii | AG06213 | TruSeq | 41074688 | 40865328 | 37679852 | 43323 | 33481355 | 35679 | 28796959 | 6429 |
| P. abelii | AG06213 | TruSeq | 41164800 | 40957098 | 37788586 | 43660 | 33497427 | 35934 | 28807666 | 6835 |
| S. syndactylus | JAMBI | TruSeq | 40689664 | 40526814 | 40063968 | 59106 | 35837316 | 49706 | 30352271 | 19962 |
| S. syndactylus | JAMBI | TruSeq | 39690240 | 39531555 | 39076332 | 57743 | 35370179 | 49115 | 30001011 | 19727 |
| S. syndactylus | JAMBI | TruSeq | 46178304 | 45942889 | 45431322 | 69251 | 42553999 | 59864 | 36185454 | 23752 |
| S. syndactylus | JAMBI | TruSeq | 46604288 | 46363777 | 45847625 | 70420 | 42903774 | 60789 | 36463191 | 23855 |
| S. syndactylus | JAMBI | TruSeq | 47636480 | 47416533 | 46879527 | 70328 | 43698426 | 60558 | 37154140 | 24162 |
| S. syndactylus | JAMBI | TruSeq | 45703168 | 45475188 | 44969801 | 67788 | 42209792 | 58835 | 35857767 | 23378 |
| S. syndactylus | JAMBI | TruSeq | 41680896 | 41406205 | 40914060 | 64845 | 39486124 | 57742 | 33682185 | 22044 |
| S. syndactylus | JAMBI | TruSeq | 39149568 | 38905371 | 38446729 | 60351 | 36950405 | 53566 | 31516397 | 20602 |

| Stat | Technique | ________ | Raw fastq reads (million) | Trimmed fastq reads | Mapped primary reads (million) | Mapped supplementary reads (million) | Mapped deduplicated primary reads (million) | Mapped deduplicated supplementary reads (million) | Mapped deduplicated high MapQ primary reads (million) | Mapped deduplicated high MapQ supplementary reads (million) |
|---|---|---|---|---|---|---|---|---|---|---|
| Mean | PDAL-Seq | | 42.233 | 39.920 | 38.154 | 0.435 | 25.319 | 0.315 | 18.560 | 0.110 |

| | | | | | | | | | | |
|---|---|---|---|---|---|---|---|---|---|---|
| Min | PDAL-Seq | | 2.682 | 2.525 | 2.480 | 0.018 | 1.667 | 0.013 | 1.191 | 0.004 |
| Max | PDAL-Seq | | 125.288 | 117.949 | 113.315 | 1.570 | 66.718 | 1.033 | 50.336 | 0.482 |
| % | PDAL-Seq | | **100.0%** | **94.5%** | **90.3%** | **1.0%** | **60.0%** | **0.7%** | **43.9%** | **0.3%** |
| Mean | TruSeq | | 49.7725 | 49.4897 | 48.4558 | 0.1051 | 45.9146 | 0.0903 | 39.9318 | 0.0211 |
| Min | TruSeq | | 33.4876 | 32.5510 | 32.4634 | 0.0371 | 31.8673 | 0.0325 | 26.9221 | 0.0048 |
| Max | TruSeq | | 319.6374 | 318.9611 | 315.3565 | 2.0578 | 303.6735 | 1.7690 | 266.8750 | 0.3710 |
| % | TruSeq | | **100.0%** | **99.4%** | **97.4%** | **0.2%** | **92.2%** | **0.2%** | **80.2%** | **0.0%** |

**Table S2:** Summary statistics for mamping of PDAL-Seq and TruSeq datasets to T2T genomes.

Column descriptions:

MapQ: weather or not the reads were filtered for MAPQ >20

Primary mapped reads: Number of primary MAPPING locations. Note secondary mapping locations are not included in this dataset.

Supplemental mapped reads: Number of supplemental MAPPING locations due to chimeric alignments.

Uniquely mapped reads: Reads where the primary mapping location is 10x better than possible secondary mapping locations.

| Species | Bio sample | Technique | MapQ | Primary mapped reads | Supplemental mapped reads | **Uniquely mapped reads** | Total | % **Fraction uniquely mapped reads** |
|---|---|---|---|---|---|---|---|---|
| H. sapiens | GM24385 | PDAL-Seq | All reads | 53858543 | 987075 | 44323013 | 54845618 | **81%** |
| H. sapiens | GM24143 | PDAL-Seq | All reads | 68756385 | 783100 | 56922429 | 69539485 | **82%** |
| H. sapiens | HEK293T | PDAL-Seq | All reads | 225667073 | 3325108 | 166750480 | 228992181 | **73%** |
| H. sapiens | CCL86-Raji | PDAL-Seq | All reads | 63138280 | 5385562 | 53106397 | 68523842 | **78%** |
| P. troglodytes | AG18354 | PDAL-Seq | All reads | 129831290 | 1438305 | 98172868 | 131269595 | **75%** |
| P. troglodytes | AG18358 | PDAL-Seq | All reads | 62882884 | 725818 | 51929908 | 63608702 | **82%** |
| P. paniscus | KB8711 | PDAL-Seq | All reads | 53110015 | 328823 | 39675862 | 53438838 | **74%** |
| P. paniscus | KB14571 | PDAL-Seq | All reads | 269323583 | 2075392 | 203168087 | 271398975 | **75%** |
| G. gorilla | KB3781 | PDAL-Seq | All reads | 80652383 | 859017 | 46956867 | 81511400 | **58%** |
| G. gorilla | F7549 | PDAL-Seq | All reads | 191900501 | 1416001 | 125544191 | 193316502 | **65%** |
| P. pygmaeus | AG05252 | PDAL-Seq | All reads | 227027229 | 1487764 | 167584703 | 228514993 | **73%** |
| P. abelii | AG06213 | PDAL-Seq | All reads | 52561105 | 250585 | 38298364 | 52811690 | **73%** |
| S. syndactylus | JAMBI | PDAL-Seq | All reads | 128512220 | 1511637 | 88781394 | 130023857 | **68%** |
| S. syndactylus | Karenina | PDAL-Seq | All reads | 51220104 | 353650 | 37234990 | 51573754 | **72%** |
| H. sapiens | GM24385 | TruSeq | All reads | 98462732 | 212515 | 88806866 | 98675247 | **90%** |

| | | | | | | | | |
|---|---|---|---|---|---|---|---|---|
| P. troglodytes | AG18354 | TruSeq | All reads | 336586881 | 289080 | 295326387 | 336875961 | **88%** |
| P. paniscus | KB8711 | TruSeq | All reads | 303673461 | 1768988 | 269673801 | 305442449 | **88%** |
| G. gorilla | KB3781 | TruSeq | All reads | 33618059 | 37999 | 26968190 | 33656058 | **80%** |
| P. pygmaeus | AG05252 | TruSeq | All reads | 323067171 | 292288 | 283198411 | 323359459 | **88%** |
| P. abelii | AG06213 | TruSeq | All reads | 284420250 | 291490 | 246190748 | 284711740 | **86%** |
| S. syndactylus | JAMBI | TruSeq | All reads | 319010015 | 450175 | 271630526 | 319460190 | **85%** |
| H. sapiens | GM24385 | PDAL-Seq | High MapQ | 43929511 | 541947 | 44220193 | 44471458 | **99%** |
| H. sapiens | GM24143 | PDAL-Seq | High MapQ | 56949046 | 269062 | 56815929 | 57218108 | **99%** |
| H. sapiens | HEK293T | PDAL-Seq | High MapQ | 167957962 | 1374012 | 166355130 | 169331974 | **98%** |
| H. sapiens | CCL86-Raji | PDAL-Seq | High MapQ | 50279518 | 2696546 | 52597604 | 52976064 | **99%** |
| P. troglodytes | AG18354 | PDAL-Seq | High MapQ | 98602252 | 541666 | 97958253 | 99143918 | **99%** |
| P. troglodytes | AG18358 | PDAL-Seq | High MapQ | 51709935 | 313205 | 51814951 | 52023140 | **100%** |
| P. paniscus | KB8711 | PDAL-Seq | High MapQ | 39602947 | 56768 | 39603439 | 39659715 | **100%** |
| P. paniscus | KB14571 | PDAL-Seq | High MapQ | 202844690 | 327286 | 202744069 | 203171976 | **100%** |
| G. gorilla | KB3781 | PDAL-Seq | High MapQ | 46960893 | 112532 | 46757298 | 47073425 | **99%** |
| G. gorilla | F7549 | PDAL-Seq | High MapQ | 125497731 | 237118 | 125195102 | 125734849 | **100%** |
| P. pygmaeus | AG05252 | PDAL-Seq | High MapQ | 167331441 | 288876 | 167348048 | 167620317 | **100%** |
| P. abelii | AG06213 | PDAL-Seq | High MapQ | 38237613 | 44024 | 38227628 | 38281637 | **100%** |
| S. syndactylus | JAMBI | PDAL-Seq | High MapQ | 89559938 | 440713 | 88501881 | 90000651 | **98%** |
| S. syndactylus | Karenina | PDAL-Seq | High MapQ | 37144379 | 100649 | 37166397 | 37245028 | **100%** |
| H. sapiens | GM24385 | TruSeq | High MapQ | 88722740 | 58240 | 88739086 | 88780980 | **100%** |

| | | | | | | | | |
|---|---|---|---|---|---|---|---|---|
| P. troglodytes | AG18354 | TruSeq | High MapQ | 294966557 | 52975 | 294970121 | 295019532 | **100%** |
| P. paniscus | KB8711 | TruSeq | High MapQ | 266874988 | 370996 | 267090092 | 267245984 | **100%** |
| G. gorilla | KB3781 | TruSeq | High MapQ | 26922096 | 4819 | 26920861 | 26926915 | **100%** |
| P. pygmaeus | AG05252 | TruSeq | High MapQ | 282879410 | 60411 | 282893904 | 282939821 | **100%** |
| P. abelii | AG06213 | TruSeq | High MapQ | 245897295 | 56242 | 245911309 | 245953537 | **100%** |
| S. syndactylus | JAMBI | TruSeq | High MapQ | 271212416 | 177482 | 271332964 | 271389898 | **100%** |
| Summary | Mean | PDAL-Seq | All reads | 118460113.9 | 1494845.5 | 87032110.93 | 119954959.4 | **73%** |
| Summary | Min | PDAL-Seq | All reads | 51220104 | 250585 | 37234990 | 51573754 | **58%** |
| Summary | Max | PDAL-Seq | All reads | 269323583 | 5385562 | 203168087 | 271398975 | **82%** |
| Summary | Mean | TruSeq | All reads | 242691224.1 | 477505 | 211684989.9 | 243168729.1 | **86%** |
| Summary | Min | TruSeq | All reads | 33618059 | 37999 | 26968190 | 33656058 | **80%** |
| Summary | Max | TruSeq | All reads | 336586881 | 1768988 | 295326387 | 336875961 | **90%** |
| Summary | Mean | PDAL-Seq | High MapQ | 86900561.14 | 524600.2857 | 86807565.86 | 87425161.43 | **99%** |
| Summary | Min | PDAL-Seq | High MapQ | 37144379 | 44024 | 37166397 | 37245028 | **98%** |
| Summary | Max | PDAL-Seq | High MapQ | 202844690 | 2696546 | 202744069 | 203171976 | **100%** |
| Summary | Mean | TruSeq | High MapQ | 211067928.9 | 111595 | 211122619.6 | 211179523.9 | **100%** |
| Summary | Min | TruSeq | High MapQ | 26922096 | 4819 | 26920861 | 26926915 | **100%** |
| Summary | Max | TruSeq | High MapQ | 294966557 | 370996 | 294970121 | 295019532 | **100%** |

**Table S3:** Multi-mapping regions of great ape genomes. Unique mapping percentages are calculated as the number of reads that have a primary alignment position with a score 10x better than their secondary mapping position divided by the total number of reads.

| Species | Cell line | Technique | Total length | Multimapping regions length | Genome in multimapping regions | Percent unique mapping reads in unique mapping region |
|---|---|---|---|---|---|---|
| Mean | | | | | 18.9% | 91% |
| H. sapiens | GM24385 | PDAL-Seq | 3117292070 | 481257732 | 15.4% | 92% |
| H. sapiens | GM24143 | PDAL-Seq | 3117292070 | 481257732 | 15.4% | 89% |
| H. sapiens | HEK293T | PDAL-Seq | 3117292070 | 481257732 | 15.4% | 80% |
| H. sapiens | CCL86-Raji | PDAL-Seq | 3117292070 | 481257732 | 15.4% | 93% |
| P. troglodytes | AG18354 | PDAL-Seq | 3177739762 | 539679762 | 17.0% | 85% |
| P. troglodytes | AG18358 | PDAL-Seq | 3177739762 | 539679762 | 17.0% | 95% |
| P. paniscus | KB8711 | PDAL-Seq | 3244888708 | 556528708 | 17.2% | 96% |
| P. paniscus | KB14571 | PDAL-Seq | 3244888708 | 556528708 | 17.2% | 88% |
| G. gorilla | KB3781 | PDAL-Seq | 3545834224 | 915464224 | 25.8% | 72% |
| G. gorilla | F7549 | PDAL-Seq | 3545834224 | 915464224 | 25.8% | 85% |
| P. pygmaeus | AG05252 | PDAL-Seq | 3220935163 | 577226452 | 17.9% | 92% |
| P. abelii | AG06213 | PDAL-Seq | 3259853530 | 627093530 | 19.2% | 86% |
| S. syndactylus | JAMBI | PDAL-Seq | 3262887708 | 715267708 | 21.9% | 76% |
| S. syndactylus | Karenina | PDAL-Seq | 3262887708 | 715267708 | 21.9% | 92% |
| H. sapiens | GM24385 | TruSeq | 3117292070 | 481257732 | 15.4% | 99% |
| P. troglodytes | AG18354 | TruSeq | 3177739762 | 539679762 | 17.0% | 99% |
| G. gorilla | KB3781 | TruSeq | 3545834224 | 915464224 | 25.8% | 99% |
| P. pygmaeus | AG05252 | TruSeq | 3220935163 | 577226452 | 17.9% | 99% |
| P. abelii | AG06213 | TruSeq | 3259853530 | 627093530 | 19.2% | 88% |
| S. syndactylus | JAMBI | TruSeq | 3262887708 | 715267708 | 21.9% | 98% |
| P. paniscus | KB8711 | TruSeq | 3244888708 | 556528708 | 17.2% | 98% |

**Table S4:** McFadden's R2 coefficient for Poisson Generalized Linear Models (GLM) fit to PDAL-Seq reads versus sets of non-B DNA motifs as predictors, in uniquely-mapping 1-kbp windows.

Column descriptions:

Fit- The parameters used in each GLM: "dedup with GC" includes k ∈ {GC, APR, CRU, TRI, DR, GQ, Z} and "dup no GC" contains l ∈ {APR, CRU, TRI, DR, GQ, Z}

PDAL-Seq data 1 kbp unique mapping windows

| Species | Cell line | Technique | Fit | # mcfadden_r2 |
|---|---|---|---|---|
| H. sapiens | GM24143 | PDAL-Seq | dedup with GC | 0.65 |
| H. sapiens | HEK293T | PDAL-Seq | dedup with GC | 0.56 |
| H. sapiens | CCL86-Raji | PDAL-Seq | dedup with GC | 0.29 |
| H. sapiens | GM24385 | PDAL-Seq | dedup with GC | 0.59 |
| P. troglodytes | AG18358 | PDAL-Seq | dedup with GC | 0.46 |
| P. troglodytes | AG18354 | PDAL-Seq | dedup with GC | 0.58 |
| P. paniscus | KB14571 | PDAL-Seq | dedup with GC | 0.44 |
| P. paniscus | KB8711 | PDAL-Seq | dedup with GC | 0.37 |
| G. gorilla | F7549 | PDAL-Seq | dedup with GC | 0.60 |
| G. gorilla | KB3781 | PDAL-Seq | dedup with GC | 0.40 |
| P. pygmaeus | AG05252 | PDAL-Seq | dedup with GC | 0.35 |
| P. abelii | AG06213 | PDAL-Seq | dedup with GC | 0.39 |
| S. syndactylus | JAMBI | PDAL-Seq | dedup with GC | 0.60 |
| S. syndactylus | Karenina | PDAL-Seq | dedup with GC | 0.50 |
| H. sapiens | GM24143 | PDAL-Seq | dedup no GC | 0.11 |
| H. sapiens | HEK293T | PDAL-Seq | dedup no GC | 0.10 |
| H. sapiens | CCL86-Raji | PDAL-Seq | dedup no GC | 0.07 |
| H. sapiens | GM24385 | PDAL-Seq | dedup no GC | 0.10 |
| P. troglodytes | AG18358 | PDAL-Seq | dedup no GC | 0.06 |
| P. troglodytes | AG18354 | PDAL-Seq | dedup no GC | 0.11 |
| P. paniscus | KB14571 | PDAL-Seq | dedup no GC | 0.05 |
| P. paniscus | KB8711 | PDAL-Seq | dedup no GC | 0.05 |
| G. gorilla | F7549 | PDAL-Seq | dedup no GC | 0.08 |
| G. gorilla | KB3781 | PDAL-Seq | dedup no GC | 0.05 |

| | | | | |
|---|---|---|---|---|
| P. pygmaeus | AG05252 | PDAL-Seq | dedup no GC | 0.06 |
| P. abelii | AG06213 | PDAL-Seq | dedup no GC | 0.04 |
| S. syndactylus | JAMBI | PDAL-Seq | dedup no GC | 0.10 |
| S. syndactylus | Karenina | PDAL-Seq | dedup no GC | 0.06 |
| TruSeq data 1 kbp unique mapping windows | | | | |
| Species | Bio_sample | Technique | Fit | mcfadden_r2 |
| | | mean | dedup with GC (set j) | 0.13 |
| | | mean | dedup no GC (set l) | 0.07 |
| H. sapiens | GM24385 | TruSeq | dedup with GC | 0.08539426 |
| P. troglodytes | AG18354 | TruSeq | dedup with GC | 0.03701044 |
| P. paniscus | KB8711 | TruSeq | dedup with GC | 0.40394369 |
| G. gorilla | KB3781 | TruSeq | dedup with GC | 0.14334882 |
| P. pygmaeus | AG05252 | TruSeq | dedup with GC | 0.07293 |
| P. abelii | AG06213 | TruSeq | dedup with GC | 0.10697006 |
| S. syndactylus | JAMBI | TruSeq | dedup with GC | 0.04659567 |
| H. sapiens | GM24385 | TruSeq | dedup no GC | 0.06728106 |
| P. troglodytes | AG18354 | TruSeq | dedup no GC | 0.03701044 |
| P. paniscus | KB8711 | TruSeq | dedup no GC | 0.1938509 |
| G. gorilla | KB3781 | TruSeq | dedup no GC | 0.05437807 |
| P. pygmaeus | AG05252 | TruSeq | dedup no GC | 0.05983437 |
| P. abelii | AG06213 | TruSeq | dedup no GC | 0.06589109 |
| S. syndactylus | JAMBI | TruSeq | dedup no GC | 0.03927525 |

**Table S5:** Parameters of the Poisson generalized linear model (GLM). GLMs fit to 1 Mbp genomic windows are on the left and GLMs fit to 1 kbp windows in uniquely mapping portions of the genome are on the right.

Fit: Predictors included in GLMs. Sets correspond to "dedup with GC" j ∈ {GC, APR, CRU, TRI, DR, G4, Z} and "dedup no GC" l ∈ {APR, CRU, TRI, DR, G4, Z}

Estimate: Coefficient for each predictor generated by the fit

Std..Error: Standard error for each predictor generated by the fit

z.value: value for f-test used to test the null hypothesis that the same variance would be observed given a parameter of zero.

Pr...z...: p-value for f-test used to test the null hypothesis that the same variance would be observed given a parameter of zero.

vif: Variance inflation factor

| 1 kbp unique mapping windows | | | | | | | | | |
|---|---|---|---|---|---|---|---|---|---|
| Species | Bio_sample | Technique | Fit | Parameter | Estimate | Std..Error | z.value | Pr...z.. | vif |
| G. gorilla | F7549 | PDAL-Seq | dedup no GC | Intercept | 8.94833451 | 8.25E-06 | 1085145.29 | 0 | NA |
| G. gorilla | F7549 | PDAL-Seq | dedup no GC | APR | -3.5823164 | 0.00077097 | -4646.478 | 0 | 1.00286015 |
| G. gorilla | F7549 | PDAL-Seq | dedup no GC | CRU | -5.8056961 | 0.00119928 | -4840.9982 | 0 | 1.02986779 |
| G. gorilla | F7549 | PDAL-Seq | dedup no GC | TRI | 0.70355221 | 0.00057716 | 1218.9977 | 0 | 1.13714897 |
| G. gorilla | F7549 | PDAL-Seq | dedup no GC | Z | 2.5501079 | 0.00058784 | 4338.09379 | 0 | 1.11006774 |
| G. gorilla | F7549 | PDAL-Seq | dedup no GC | DR | 0.29535615 | 0.00016136 | 1830.39566 | 0 | 1.54927737 |
| G. gorilla | F7549 | PDAL-Seq | dedup no GC | G4 | 1.76950337 | 0.00018443 | 9594.40809 | 0 | 1.29267365 |
| P. pygmaeus | AG05252 | TruSeq | dedup no GC | Intercept | 9.6631385 | 6.03E-06 | 1602525.93 | 0 | NA |
| P. pygmaeus | AG05252 | TruSeq | dedup no GC | APR | 0.12993959 | 0.00051144 | 254.066278 | 0 | 1.00334078 |
| P. pygmaeus | AG05252 | TruSeq | dedup no GC | CRU | -0.5988403 | 0.00074648 | -802.21765 | 0 | 1.04209755 |
| P. pygmaeus | AG05252 | TruSeq | dedup no GC | TRI | -0.458719 | 0.00049148 | -933.33472 | 0 | 1.08224022 |
| P. pygmaeus | AG05252 | TruSeq | dedup no GC | Z | 0.2174565 | 0.00053094 | 409.565577 | 0 | 1.06122361 |
| P. pygmaeus | AG05252 | TruSeq | dedup no GC | DR | -0.41392 | 0.00011783 | -3512.7655 | 0 | 1.17268073 |
| P. pygmaeus | AG05252 | TruSeq | dedup no GC | G4 | -0.5580845 | 0.00018719 | -2981.3732 | 0 | 1.0295329 |
| P. abelii | AG06213 | TruSeq | dedup no GC | Intercept | 9.52765547 | 6.48E-06 | 1470841.7 | 0 | NA |
| P. abelii | AG06213 | TruSeq | dedup no GC | APR | 0.2848428 | 0.0005498 | 518.080884 | 0 | 1.00320549 |
| P. abelii | AG06213 | TruSeq | dedup no GC | CRU | -0.4667639 | 0.00079564 | -586.65377 | 0 | 1.05829405 |
| P. abelii | AG06213 | TruSeq | dedup no GC | TRI | -0.468237 | 0.00052974 | -883.904 | 0 | 1.10478867 |
| P. abelii | AG06213 | TruSeq | dedup no GC | Z | 0.08172184 | 0.00057897 | 141.149909 | 0 | 1.07542377 |
| P. abelii | AG06213 | TruSeq | dedup no GC | DR | -0.4369234 | 0.0001481 | -2950.1755 | 0 | 1.2421062 |
| P. abelii | AG06213 | TruSeq | dedup no GC | G4 | -0.7636828 | 0.00020618 | -3703.9131 | 0 | 1.03518098 |

| | | | | | | | | | |
|---|---|---|---|---|---|---|---|---|---|
| H. sapiens | GM24143 | PDAL-Seq | dedup no GC | Intercept | 7.92554082 | 1.34E-05 | 591327.889 | 0 | NA |
| H. sapiens | GM24143 | PDAL-Seq | dedup no GC | APR | -6.8250837 | 0.00133357 | -5117.9181 | 0 | 1.0029107 |
| H. sapiens | GM24143 | PDAL-Seq | dedup no GC | CRU | -9.562548 | 0.00208801 | -4579.739 | 0 | 1.01939616 |
| H. sapiens | GM24143 | PDAL-Seq | dedup no GC | TRI | 0.45759405 | 0.00079133 | 578.257408 | 0 | 1.10790747 |
| H. sapiens | GM24143 | PDAL-Seq | dedup no GC | Z | 4.50745282 | 0.00077054 | 5849.76922 | 0 | 1.12436394 |
| H. sapiens | GM24143 | PDAL-Seq | dedup no GC | DR | 0.74204346 | 0.00021099 | 3517.01847 | 0 | 2.12827912 |
| H. sapiens | GM24143 | PDAL-Seq | dedup no GC | G4 | 2.64799289 | 0.00023386 | 11322.7587 | 0 | 1.88708849 |
| H. sapiens | GM24385 | TruSeq | dedup no GC | Intercept | 8.53024089 | 1.08E-05 | 789849.351 | 0 | NA |
| H. sapiens | GM24385 | TruSeq | dedup no GC | APR | 0.42674317 | 0.00090416 | 471.977158 | 0 | 1.00372249 |
| H. sapiens | GM24385 | TruSeq | dedup no GC | CRU | 0.77216899 | 0.00122967 | 627.946015 | 0 | 1.07037083 |
| H. sapiens | GM24385 | TruSeq | dedup no GC | TRI | -1.005903 | 0.00088088 | -1141.936 | 0 | 1.11578393 |
| H. sapiens | GM24385 | TruSeq | dedup no GC | Z | -1.3060955 | 0.00106936 | -1221.3757 | 0 | 1.06309373 |
| H. sapiens | GM24385 | TruSeq | dedup no GC | DR | -1.0040698 | 0.00026935 | -3727.738 | 0 | 1.22737188 |
| H. sapiens | GM24385 | TruSeq | dedup no GC | G4 | -1.0424347 | 0.00035636 | -2925.2401 | 0 | 1.0233127 |
| H. sapiens | HEK293T | PDAL-Seq | dedup no GC | Intercept | 8.67925103 | 9.12E-06 | 951749.419 | 0 | NA |
| H. sapiens | HEK293T | PDAL-Seq | dedup no GC | APR | -6.7655379 | 0.00090812 | -7450.0568 | 0 | 1.00259779 |
| H. sapiens | HEK293T | PDAL-Seq | dedup no GC | CRU | -9.6501136 | 0.00136644 | -7062.2341 | 0 | 1.02237612 |
| H. sapiens | HEK293T | PDAL-Seq | dedup no GC | TRI | 1.6019555 | 0.00046137 | 3472.17377 | 0 | 1.1410866 |
| H. sapiens | HEK293T | PDAL-Seq | dedup no GC | Z | 4.15360208 | 0.00048996 | 8477.37186 | 0 | 1.13113621 |
| H. sapiens | HEK293T | PDAL-Seq | dedup no GC | DR | 1.33796235 | 0.00012696 | 10538.6284 | 0 | 1.93181716 |
| H. sapiens | HEK293T | PDAL-Seq | dedup no GC | G4 | 1.93863929 | 0.00015583 | 12440.6862 | 0 | 1.68029219 |
| P. abelii | AG06213 | PDAL-Seq | dedup no GC | Intercept | 7.61886408 | 1.59E-05 | 478449.922 | 0 | NA |
| P. abelii | AG06213 | PDAL-Seq | dedup no GC | APR | -4.5447306 | 0.00151685 | -2996.1651 | 0 | 1.00231026 |
| P. abelii | AG06213 | PDAL-Seq | dedup no GC | CRU | -6.205359 | 0.00229121 | -2708.3311 | 0 | 1.03115234 |
| P. abelii | AG06213 | PDAL-Seq | dedup no GC | TRI | 1.33923128 | 0.00104026 | 1287.40019 | 0 | 1.15226151 |
| P. abelii | AG06213 | PDAL-Seq | dedup no GC | Z | 1.82253602 | 0.00106176 | 1716.51987 | 0 | 1.10782727 |
| P. abelii | AG06213 | PDAL-Seq | dedup no GC | DR | 0.73588492 | 0.00027781 | 2648.9163 | 0 | 1.54637196 |
| P. abelii | AG06213 | PDAL-Seq | dedup no GC | G4 | 1.23867451 | 0.00035193 | 3519.65794 | 0 | 1.29076654 |

| | | | | | | | | | |
|---|---|---|---|---|---|---|---|---|---|
| H. sapiens | CCL86-Raji | PDAL-Seq | dedup no GC | Intercept | 7.89164887 | 1.35E-05 | 582508.631 | 0 | NA |
| H. sapiens | CCL86-Raji | PDAL-Seq | dedup no GC | APR | -4.2435309 | 0.0012883 | -3293.8909 | 0 | 1.00244362 |
| H. sapiens | CCL86-Raji | PDAL-Seq | dedup no GC | CRU | -11.716029 | 0.00204612 | -5725.9707 | 0 | 1.02210213 |
| H. sapiens | CCL86-Raji | PDAL-Seq | dedup no GC | TRI | 3.66191988 | 0.00058343 | 6276.55627 | 0 | 1.23213768 |
| H. sapiens | CCL86-Raji | PDAL-Seq | dedup no GC | Z | 3.41286463 | 0.00072054 | 4736.51652 | 0 | 1.11967716 |
| H. sapiens | CCL86-Raji | PDAL-Seq | dedup no GC | DR | 1.72938596 | 0.00017442 | 9914.84311 | 0 | 1.55640727 |
| H. sapiens | CCL86-Raji | PDAL-Seq | dedup no GC | G4 | 0.40218262 | 0.00027266 | 1475.05969 | 0 | 1.26865918 |
| P. troglodytes | AG18358 | PDAL-Seq | dedup no GC | Intercept | 7.87953178 | 1.41E-05 | 560603.13 | 0 | NA |
| P. troglodytes | AG18358 | PDAL-Seq | dedup no GC | APR | -4.0291769 | 0.00130798 | -3080.4508 | 0 | 1.00320735 |
| P. troglodytes | AG18358 | PDAL-Seq | dedup no GC | CRU | -6.72789 | 0.00199468 | -3372.9231 | 0 | 1.03213842 |
| P. troglodytes | AG18358 | PDAL-Seq | dedup no GC | TRI | 1.79109731 | 0.00096279 | 1860.32102 | 0 | 1.11668929 |
| P. troglodytes | AG18358 | PDAL-Seq | dedup no GC | Z | 2.58213021 | 0.00099604 | 2592.3896 | 0 | 1.09254939 |
| P. troglodytes | AG18358 | PDAL-Seq | dedup no GC | DR | 0.66358832 | 0.00025241 | 2629.06016 | 0 | 1.48838545 |
| P. troglodytes | AG18358 | PDAL-Seq | dedup no GC | G4 | 1.44424411 | 0.00031067 | 4648.8771 | 0 | 1.26356087 |
| S. syndactylus | JAMBI | TruSeq | dedup no GC | Intercept | 9.66336843 | 6.18E-06 | 1562623.4 | 0 | NA |
| S. syndactylus | JAMBI | TruSeq | dedup no GC | APR | 0.10358023 | 0.00052588 | 196.965019 | 0 | 1.00356133 |
| S. syndactylus | JAMBI | TruSeq | dedup no GC | CRU | -0.7634667 | 0.0007869 | -970.22203 | 0 | 1.04139892 |
| S. syndactylus | JAMBI | TruSeq | dedup no GC | TRI | -0.3928755 | 0.00047566 | -825.96554 | 0 | 1.13504797 |
| S. syndactylus | JAMBI | TruSeq | dedup no GC | Z | 0.16069149 | 0.000583 | 275.627216 | 0 | 1.06746834 |
| S. syndactylus | JAMBI | TruSeq | dedup no GC | DR | -0.4586417 | 0.00015444 | -2969.7699 | 0 | 1.24766339 |
| S. syndactylus | JAMBI | TruSeq | dedup no GC | G4 | -0.6351166 | 0.00019703 | -3223.3814 | 0 | 1.0287918 |
| P. pygmaeus | AG05252 | PDAL-Seq | dedup no GC | Intercept | 9.09089205 | 7.74E-06 | 1174103.57 | 0 | NA |
| P. pygmaeus | AG05252 | PDAL-Seq | dedup no GC | APR | -2.3573055 | 0.00069867 | -3373.9739 | 0 | 1.00281554 |
| P. pygmaeus | AG05252 | PDAL-Seq | dedup no GC | CRU | -5.0541882 | 0.00103302 | -4892.6444 | 0 | 1.03174244 |
| P. pygmaeus | AG05252 | PDAL-Seq | dedup no GC | TRI | 1.20539528 | 0.00053319 | 2260.74044 | 0 | 1.09706095 |
| P. pygmaeus | AG05252 | PDAL-Seq | dedup no GC | Z | 0.14123402 | 0.00058018 | 243.430983 | 0 | 1.05720446 |
| P. pygmaeus | AG05252 | PDAL-Seq | dedup no GC | DR | 1.00501217 | 0.00010998 | 9138.12315 | 0 | 1.22791271 |
| P. pygmaeus | AG05252 | PDAL-Seq | dedup no GC | G4 | 0.04791778 | 0.00019549 | 245.119983 | 0 | 1.09022282 |

| | | | | | | | | | |
|---|---|---|---|---|---|---|---|---|---|
| P. paniscus | KB14571 | PDAL-Seq | dedup no GC | Intercept | 9.26310381 | 7.07E-06 | 1310060.52 | 0 | NA |
| P. paniscus | KB14571 | PDAL-Seq | dedup no GC | APR | -3.2112933 | 0.00064634 | -4968.4282 | 0 | 1.00292231 |
| P. paniscus | KB14571 | PDAL-Seq | dedup no GC | CRU | -4.7624563 | 0.0009355 | -5090.8118 | 0 | 1.0447357 |
| P. paniscus | KB14571 | PDAL-Seq | dedup no GC | TRI | 2.14534478 | 0.00049554 | 4329.26364 | 0 | 1.12013315 |
| P. paniscus | KB14571 | PDAL-Seq | dedup no GC | Z | 1.09934886 | 0.00057705 | 1905.10259 | 0 | 1.0726033 |
| P. paniscus | KB14571 | PDAL-Seq | dedup no GC | DR | 0.50683015 | 0.0001388 | 3651.56707 | 0 | 1.38139375 |
| P. paniscus | KB14571 | PDAL-Seq | dedup no GC | G4 | 0.92808773 | 0.00017402 | 5333.17611 | 0 | 1.16723698 |
| P. troglodytes | AG18354 | PDAL-Seq | dedup no GC | Intercept | 8.24952356 | 1.15E-05 | 717865.942 | 0 | NA |
| P. troglodytes | AG18354 | PDAL-Seq | dedup no GC | APR | -4.6067334 | 0.00108913 | -4229.7244 | 0 | 1.00307779 |
| P. troglodytes | AG18354 | PDAL-Seq | dedup no GC | CRU | -7.6467472 | 0.00163532 | -4675.9869 | 0 | 1.03103451 |
| P. troglodytes | AG18354 | PDAL-Seq | dedup no GC | TRI | 1.54118709 | 0.00074175 | 2077.76279 | 0 | 1.11253653 |
| P. troglodytes | AG18354 | PDAL-Seq | dedup no GC | Z | 3.91376297 | 0.00064914 | 6029.16272 | 0 | 1.13629043 |
| P. troglodytes | AG18354 | PDAL-Seq | dedup no GC | DR | 1.17753251 | 0.00018251 | 6451.91448 | 0 | 1.64934954 |
| P. troglodytes | AG18354 | PDAL-Seq | dedup no GC | G4 | 1.59395399 | 0.00022887 | 6964.41225 | 0 | 1.37211768 |
| G. gorilla | KB3781 | PDAL-Seq | dedup no GC | Intercept | 7.75423396 | 1.49E-05 | 519493.212 | 0 | NA |
| G. gorilla | KB3781 | PDAL-Seq | dedup no GC | APR | -3.7169864 | 0.00140029 | -2654.44 | 0 | 1.00269411 |
| G. gorilla | KB3781 | PDAL-Seq | dedup no GC | CRU | -6.4982495 | 0.00215613 | -3013.8419 | 0 | 1.03183043 |
| G. gorilla | KB3781 | PDAL-Seq | dedup no GC | TRI | 1.67703454 | 0.00096755 | 1733.27475 | 0 | 1.16252682 |
| G. gorilla | KB3781 | PDAL-Seq | dedup no GC | Z | 1.68361961 | 0.00108766 | 1547.93071 | 0 | 1.09320559 |
| G. gorilla | KB3781 | PDAL-Seq | dedup no GC | DR | 0.68752786 | 0.00027282 | 2520.06742 | 0 | 1.51985585 |
| G. gorilla | KB3781 | PDAL-Seq | dedup no GC | G4 | 1.3071699 | 0.00033711 | 3877.62511 | 0 | 1.26371146 |
| G. gorilla | KB3781 | TruSeq | dedup no GC | Intercept | 7.33522539 | 1.98E-05 | 371010.52 | 0 | NA |
| G. gorilla | KB3781 | TruSeq | dedup no GC | APR | 1.25305345 | 0.00163437 | 766.690762 | 0 | 1.00324595 |
| G. gorilla | KB3781 | TruSeq | dedup no GC | CRU | 1.70477382 | 0.00214455 | 794.931337 | 0 | 1.09435462 |
| G. gorilla | KB3781 | TruSeq | dedup no GC | TRI | -1.1481033 | 0.00163839 | -700.75231 | 0 | 1.11497518 |
| G. gorilla | KB3781 | TruSeq | dedup no GC | Z | -2.5460068 | 0.00206021 | -1235.8012 | 0 | 1.05796351 |
| G. gorilla | KB3781 | TruSeq | dedup no GC | DR | -0.6441764 | 0.00049759 | -1294.6015 | 0 | 1.24349322 |
| G. gorilla | KB3781 | TruSeq | dedup no GC | G4 | -2.7224753 | 0.00073962 | -3680.9341 | 0 | 1.01396823 |

| | | | | | | | | | |
|---|---|---|---|---|---|---|---|---|---|
| P. paniscus | KB8711 | PDAL-Seq | dedup no GC | Intercept | 7.65203036 | 1.57E-05 | 488837.058 | 0 | NA |
| P. paniscus | KB8711 | PDAL-Seq | dedup no GC | APR | -4.0258291 | 0.00146256 | -2752.5996 | 0 | 1.00281151 |
| P. paniscus | KB8711 | PDAL-Seq | dedup no GC | CRU | -6.0906983 | 0.00219688 | -2772.4329 | 0 | 1.03536651 |
| P. paniscus | KB8711 | PDAL-Seq | dedup no GC | TRI | 1.81177943 | 0.00108448 | 1670.64968 | 0 | 1.11575835 |
| P. paniscus | KB8711 | PDAL-Seq | dedup no GC | Z | 0.94934847 | 0.00127972 | 741.839336 | 0 | 1.0655808 |
| P. paniscus | KB8711 | PDAL-Seq | dedup no GC | DR | 0.57846657 | 0.00029623 | 1952.7372 | 0 | 1.5357648 |
| P. paniscus | KB8711 | PDAL-Seq | dedup no GC | G4 | 1.7745245 | 0.0003395 | 5226.90347 | 0 | 1.33646043 |
| H. sapiens | GM24385 | PDAL-Seq | dedup no GC | Intercept | 7.70471989 | 1.50E-05 | 512961.309 | 0 | NA |
| H. sapiens | GM24385 | PDAL-Seq | dedup no GC | APR | -6.1764403 | 0.00147236 | -4194.9371 | 0 | 1.00286297 |
| H. sapiens | GM24385 | PDAL-Seq | dedup no GC | CRU | -10.7524 | 0.00236876 | -4539.2455 | 0 | 1.01823029 |
| H. sapiens | GM24385 | PDAL-Seq | dedup no GC | TRI | -0.0590138 | 0.00094093 | -62.718902 | 0 | 1.09380499 |
| H. sapiens | GM24385 | PDAL-Seq | dedup no GC | Z | 5.23515879 | 0.0007612 | 6877.49903 | 0 | 1.17016328 |
| H. sapiens | GM24385 | PDAL-Seq | dedup no GC | DR | 1.04987355 | 0.00022391 | 4688.78434 | 0 | 1.87823541 |
| H. sapiens | GM24385 | PDAL-Seq | dedup no GC | G4 | 2.05521985 | 0.00026978 | 7618.00485 | 0 | 1.60973333 |
| S. syndactylus | JAMBI | PDAL-Seq | dedup no GC | Intercept | 8.28367855 | 1.14E-05 | 723883.907 | 0 | NA |
| S. syndactylus | JAMBI | PDAL-Seq | dedup no GC | APR | -6.2467846 | 0.00113333 | -5511.8631 | 0 | 1.0029585 |
| S. syndactylus | JAMBI | PDAL-Seq | dedup no GC | CRU | -7.4512561 | 0.00175566 | -4244.139 | 0 | 1.017964 |
| S. syndactylus | JAMBI | PDAL-Seq | dedup no GC | TRI | 0.95405695 | 0.00070403 | 1355.13265 | 0 | 1.18375612 |
| S. syndactylus | JAMBI | PDAL-Seq | dedup no GC | Z | 4.40464546 | 0.00055822 | 7890.46136 | 0 | 1.28122763 |
| S. syndactylus | JAMBI | PDAL-Seq | dedup no GC | DR | 0.9404346 | 0.0002082 | 4517.03305 | 0 | 1.84915956 |
| S. syndactylus | JAMBI | PDAL-Seq | dedup no GC | G4 | 2.17967187 | 0.0002361 | 9232.04347 | 0 | 1.36601056 |
| P. troglodytes | AG18354 | TruSeq | dedup no GC | Intercept | 9.71253063 | 5.90E-06 | 1645160.15 | 0 | NA |
| P. troglodytes | AG18354 | TruSeq | dedup no GC | APR | -0.0642178 | 0.00049947 | -128.57277 | 0 | 1.00375125 |
| P. troglodytes | AG18354 | TruSeq | dedup no GC | CRU | -0.8411129 | 0.00070467 | -1193.6185 | 0 | 1.06392714 |
| P. troglodytes | AG18354 | TruSeq | dedup no GC | TRI | -0.1672152 | 0.00047764 | -350.08802 | 0 | 1.10070243 |
| P. troglodytes | AG18354 | TruSeq | dedup no GC | Z | 0.03320113 | 0.00055563 | 59.7536024 | 0 | 1.06523187 |
| P. troglodytes | AG18354 | TruSeq | dedup no GC | DR | -0.4184566 | 0.00013592 | -3078.7069 | 0 | 1.23334265 |
| P. troglodytes | AG18354 | TruSeq | dedup no GC | G4 | -0.3960495 | 0.00018025 | -2197.1684 | 0 | 1.04072536 |

| | | | | | | | | | |
|---|---|---|---|---|---|---|---|---|---|
| S. syndactylus | Karenina | PDAL-Seq | dedup no GC | Intercept | 7.58528121 | 1.66E-05 | 458201.142 | 0 | NA |
| S. syndactylus | Karenina | PDAL-Seq | dedup no GC | APR | -4.851467 | 0.00158506 | -3060.7397 | 0 | 1.00319605 |
| S. syndactylus | Karenina | PDAL-Seq | dedup no GC | CRU | -6.8079847 | 0.00255884 | -2660.5723 | 0 | 1.01831231 |
| S. syndactylus | Karenina | PDAL-Seq | dedup no GC | TRI | 1.50400767 | 0.0010966 | 1371.51936 | 0 | 1.18198617 |
| S. syndactylus | Karenina | PDAL-Seq | dedup no GC | Z | 2.04248993 | 0.00127137 | 1606.52463 | 0 | 1.09096388 |
| S. syndactylus | Karenina | PDAL-Seq | dedup no GC | DR | 0.08454476 | 0.00035153 | 240.505154 | 0 | 1.60708971 |
| S. syndactylus | Karenina | PDAL-Seq | dedup no GC | G4 | 2.50866304 | 0.00036022 | 6964.23337 | 0 | 1.31382873 |
| P. paniscus | KB8711 | TruSeq | dedup no GC | Intercept | 10.0790841 | 4.99E-06 | 2019784.96 | 0 | NA |
| P. paniscus | KB8711 | TruSeq | dedup no GC | APR | 1.63913969 | 0.00040704 | 4026.96021 | 0 | 1.00290601 |
| P. paniscus | KB8711 | TruSeq | dedup no GC | CRU | 0.99155409 | 0.00050629 | 1958.45456 | 0 | 1.12221634 |
| P. paniscus | KB8711 | TruSeq | dedup no GC | TRI | -0.2192384 | 0.00040557 | -540.56905 | 0 | 1.10481774 |
| P. paniscus | KB8711 | TruSeq | dedup no GC | Z | -1.0533343 | 0.00049221 | -2139.9993 | 0 | 1.06386234 |
| P. paniscus | KB8711 | TruSeq | dedup no GC | DR | -0.214063 | 0.00011894 | -1799.7197 | 0 | 1.2617849 |
| P. paniscus | KB8711 | TruSeq | dedup no GC | G4 | -6.3116287 | 0.00022745 | -27749.306 | 0 | 1.00731862 |
| G. gorilla | F7549 | PDAL-Seq | dedup with GC | Intercept | 7.39384538 | 4.54E-05 | 163032.721 | 0 | NA |
| G. gorilla | F7549 | PDAL-Seq | dedup with GC | APR | -0.132688 | 0.0007747 | -171.27652 | 0 | 1.02238255 |
| G. gorilla | F7549 | PDAL-Seq | dedup with GC | CRU | -2.1573556 | 0.00117248 | -1839.9963 | 0 | 1.03120705 |
| G. gorilla | F7549 | PDAL-Seq | dedup with GC | TRI | 0.86681945 | 0.00060101 | 1442.26236 | 0 | 1.10333837 |
| G. gorilla | F7549 | PDAL-Seq | dedup with GC | Z | 1.1044557 | 0.00059783 | 1847.45248 | 0 | 1.09940226 |
| G. gorilla | F7549 | PDAL-Seq | dedup with GC | DR | 0.56904414 | 0.00015027 | 3786.75311 | 0 | 1.22435845 |
| G. gorilla | F7549 | PDAL-Seq | dedup with GC | G4 | -2.1877738 | 0.00025494 | -8581.5401 | 0 | 1.41343962 |
| G. gorilla | F7549 | PDAL-Seq | dedup with GC | GC_content | 3.76572574 | 0.00010654 | 35347.2834 | 0 | 1.43010123 |
| P. pygmaeus | AG05252 | TruSeq | dedup with GC | Intercept | 9.75036291 | 3.24E-05 | 300598.428 | 0 | NA |
| P. pygmaeus | AG05252 | TruSeq | dedup with GC | APR | -0.0368102 | 0.00051532 | -71.431683 | 0 | 1.01742049 |
| P. pygmaeus | AG05252 | TruSeq | dedup with GC | CRU | -0.8000935 | 0.00075146 | -1064.7207 | 0 | 1.05266874 |
| P. pygmaeus | AG05252 | TruSeq | dedup with GC | TRI | -0.451821 | 0.00049076 | -920.65412 | 0 | 1.08333102 |
| P. pygmaeus | AG05252 | TruSeq | dedup with GC | Z | 0.29390542 | 0.00053158 | 552.886436 | 0 | 1.0641553 |
| P. pygmaeus | AG05252 | TruSeq | dedup with GC | DR | -0.424041 | 0.00011801 | -3593.2265 | 0 | 1.17974256 |

| | | | | | | | | | |
|---|---|---|---|---|---|---|---|---|---|
| P. pygmaeus | AG05252 | TruSeq | dedup with GC | G4 | -0.2899863 | 0.00020915 | -1386.502 | 0 | 1.31763763 |
| P. pygmaeus | AG05252 | TruSeq | dedup with GC | GC_content | -0.2176278 | 7.96E-05 | -2734.5101 | 0 | 1.31813016 |
| P. abelii | AG06213 | TruSeq | dedup with GC | Intercept | 9.69199365 | 3.48E-05 | 278151.628 | 0 | NA |
| P. abelii | AG06213 | TruSeq | dedup with GC | APR | -0.0332721 | 0.00055396 | -60.062238 | 0 | 1.01748199 |
| P. abelii | AG06213 | TruSeq | dedup with GC | CRU | -0.8325048 | 0.00080234 | -1037.5918 | 0 | 1.06898111 |
| P. abelii | AG06213 | TruSeq | dedup with GC | TRI | -0.4445216 | 0.00052837 | -841.31322 | 0 | 1.10744679 |
| P. abelii | AG06213 | TruSeq | dedup with GC | Z | 0.23495779 | 0.00057996 | 405.130925 | 0 | 1.07854662 |
| P. abelii | AG06213 | TruSeq | dedup with GC | DR | -0.4641435 | 0.00014851 | -3125.414 | 0 | 1.25775765 |
| P. abelii | AG06213 | TruSeq | dedup with GC | G4 | -0.249261 | 0.00022841 | -1091.3033 | 0 | 1.3282764 |
| P. abelii | AG06213 | TruSeq | dedup with GC | GC_content | -0.4105239 | 8.57E-05 | -4792.4498 | 0 | 1.31595116 |
| H. sapiens | GM24143 | PDAL-Seq | dedup with GC | Intercept | 5.15458663 | 7.37E-05 | 69955.8779 | 0 | NA |
| H. sapiens | GM24143 | PDAL-Seq | dedup with GC | APR | 0.08793091 | 0.00133023 | 66.1021987 | 0 | 1.02674611 |
| H. sapiens | GM24143 | PDAL-Seq | dedup with GC | CRU | -2.4498749 | 0.00202401 | -1210.4064 | 0 | 1.01994422 |
| H. sapiens | GM24143 | PDAL-Seq | dedup with GC | TRI | 1.36592958 | 0.00087693 | 1557.6205 | 0 | 1.07815994 |
| H. sapiens | GM24143 | PDAL-Seq | dedup with GC | Z | 1.89486817 | 0.00078812 | 2404.29343 | 0 | 1.08484313 |
| H. sapiens | GM24143 | PDAL-Seq | dedup with GC | DR | 1.02303301 | 0.00016673 | 6135.75938 | 0 | 1.16919755 |
| H. sapiens | GM24143 | PDAL-Seq | dedup with GC | G4 | -2.6971972 | 0.00034315 | -7860.1052 | 0 | 1.46814131 |
| H. sapiens | GM24143 | PDAL-Seq | dedup with GC | GC_content | 6.54520799 | 0.00016678 | 39245.5404 | 0 | 1.51765335 |
| H. sapiens | GM24385 | TruSeq | dedup with GC | Intercept | 8.725154 | 5.79E-05 | 150569.864 | 0 | NA |
| H. sapiens | GM24385 | TruSeq | dedup with GC | APR | 0.05738651 | 0.00091102 | 62.9914541 | 0 | 1.01764846 |
| H. sapiens | GM24385 | TruSeq | dedup with GC | CRU | 0.34920434 | 0.00124541 | 280.392462 | 0 | 1.08205866 |
| H. sapiens | GM24385 | TruSeq | dedup with GC | TRI | -0.9505617 | 0.00087967 | -1080.5856 | 0 | 1.118587 |
| H. sapiens | GM24385 | TruSeq | dedup with GC | Z | -1.1614967 | 0.00107135 | -1084.1445 | 0 | 1.06475988 |
| H. sapiens | GM24385 | TruSeq | dedup with GC | DR | -1.0197578 | 0.00026986 | -3778.8794 | 0 | 1.23838859 |
| H. sapiens | GM24385 | TruSeq | dedup with GC | G4 | -0.3908397 | 0.00039702 | -984.44003 | 0 | 1.32609148 |
| H. sapiens | GM24385 | TruSeq | dedup with GC | GC_content | -0.4878971 | 0.00014278 | -3417.0598 | 0 | 1.33104223 |
| H. sapiens | HEK293T | PDAL-Seq | dedup with GC | Intercept | 6.07242316 | 5.02E-05 | 120963.058 | 0 | NA |
| H. sapiens | HEK293T | PDAL-Seq | dedup with GC | APR | -0.3051704 | 0.00090635 | -336.70112 | 0 | 1.02588114 |

| | | | | | | | | | |
|---|---|---|---|---|---|---|---|---|---|
| H. sapiens | HEK293T | PDAL-Seq | dedup with GC | CRU | -2.5904354 | 0.00132668 | -1952.5673 | 0 | 1.02148009 |
| H. sapiens | HEK293T | PDAL-Seq | dedup with GC | TRI | 2.81414461 | 0.00051128 | 5504.06776 | 0 | 1.10634856 |
| H. sapiens | HEK293T | PDAL-Seq | dedup with GC | Z | 2.02744662 | 0.00049892 | 4063.66483 | 0 | 1.08756495 |
| H. sapiens | HEK293T | PDAL-Seq | dedup with GC | DR | 1.34354422 | 0.00010506 | 12788.2412 | 0 | 1.20755173 |
| H. sapiens | HEK293T | PDAL-Seq | dedup with GC | G4 | -2.896012 | 0.00023688 | -12225.867 | 0 | 1.43091712 |
| H. sapiens | HEK293T | PDAL-Seq | dedup with GC | GC_content | 6.17146139 | 0.00011409 | 54091.3699 | 0 | 1.48568011 |
| P. abelii | AG06213 | PDAL-Seq | dedup with GC | Intercept | 5.82805689 | 8.81E-05 | 66189.555 | 0 | NA |
| P. abelii | AG06213 | PDAL-Seq | dedup with GC | APR | -0.5099753 | 0.00152282 | -334.88926 | 0 | 1.02233781 |
| P. abelii | AG06213 | PDAL-Seq | dedup with GC | CRU | -1.7573917 | 0.00224095 | -784.2169 | 0 | 1.03041744 |
| P. abelii | AG06213 | PDAL-Seq | dedup with GC | TRI | 2.05535441 | 0.00111343 | 1845.97028 | 0 | 1.10473428 |
| P. abelii | AG06213 | PDAL-Seq | dedup with GC | Z | 0.16604352 | 0.00109562 | 151.552253 | 0 | 1.09476723 |
| P. abelii | AG06213 | PDAL-Seq | dedup with GC | DR | 0.891498 | 0.00025646 | 3476.1767 | 0 | 1.21341701 |
| P. abelii | AG06213 | PDAL-Seq | dedup with GC | G4 | -3.1731558 | 0.00050362 | -6300.7494 | 0 | 1.38244516 |
| P. abelii | AG06213 | PDAL-Seq | dedup with GC | GC_content | 4.32762046 | 0.00020585 | 21022.9477 | 0 | 1.41248515 |
| H. sapiens | CCL86-Raji | PDAL-Seq | dedup with GC | Intercept | 5.86285865 | 7.54E-05 | 77793.3301 | 0 | NA |
| H. sapiens | CCL86-Raji | PDAL-Seq | dedup with GC | APR | 0.42062841 | 0.00128701 | 326.826975 | 0 | 1.02403187 |
| H. sapiens | CCL86-Raji | PDAL-Seq | dedup with GC | CRU | -5.7984832 | 0.00203537 | -2848.8573 | 0 | 1.02212908 |
| H. sapiens | CCL86-Raji | PDAL-Seq | dedup with GC | TRI | 4.80279308 | 0.00062974 | 7626.57824 | 0 | 1.18102533 |
| H. sapiens | CCL86-Raji | PDAL-Seq | dedup with GC | Z | 2.11093931 | 0.00072716 | 2902.99575 | 0 | 1.09298977 |
| H. sapiens | CCL86-Raji | PDAL-Seq | dedup with GC | DR | 1.47269678 | 0.00016186 | 9098.64918 | 0 | 1.29127631 |
| H. sapiens | CCL86-Raji | PDAL-Seq | dedup with GC | G4 | -4.0974622 | 0.00042758 | -9582.9189 | 0 | 1.36071712 |
| H. sapiens | CCL86-Raji | PDAL-Seq | dedup with GC | GC_content | 4.87382223 | 0.00017504 | 27843.5636 | 0 | 1.42626434 |
| P. troglodytes | AG18358 | PDAL-Seq | dedup with GC | Intercept | 6.16091951 | 7.72E-05 | 79851.5451 | 0 | NA |
| P. troglodytes | AG18358 | PDAL-Seq | dedup with GC | APR | -0.2235332 | 0.00131174 | -170.40918 | 0 | 1.0232398 |
| P. troglodytes | AG18358 | PDAL-Seq | dedup with GC | CRU | -2.492373 | 0.00194801 | -1279.4433 | 0 | 1.03281149 |
| P. troglodytes | AG18358 | PDAL-Seq | dedup with GC | TRI | 2.04626492 | 0.00100081 | 2044.61002 | 0 | 1.0885107 |
| P. troglodytes | AG18358 | PDAL-Seq | dedup with GC | Z | 1.04643668 | 0.00102429 | 1021.62036 | 0 | 1.08570453 |
| P. troglodytes | AG18358 | PDAL-Seq | dedup with GC | DR | 0.78647563 | 0.00023485 | 3348.90534 | 0 | 1.19192375 |

| | | | | | | | | | |
|---|---|---|---|---|---|---|---|---|---|
| P. troglodytes | AG18358 | PDAL-Seq | dedup with GC | G4 | -2.9055183 | 0.00043985 | -6605.6287 | 0 | 1.40051578 |
| P. troglodytes | AG18358 | PDAL-Seq | dedup with GC | GC_content | 4.15977256 | 0.00018073 | 23017.0641 | 0 | 1.43141149 |
| S. syndactylus | JAMBI | TruSeq | dedup with GC | Intercept | 9.74321558 | 3.33E-05 | 292190.426 | 0 | NA |
| S. syndactylus | JAMBI | TruSeq | dedup with GC | APR | -0.0518701 | 0.00052983 | -97.899594 | 0 | 1.01818531 |
| S. syndactylus | JAMBI | TruSeq | dedup with GC | CRU | -0.9483528 | 0.00079169 | -1197.8897 | 0 | 1.05136906 |
| S. syndactylus | JAMBI | TruSeq | dedup with GC | TRI | -0.372292 | 0.00047537 | -783.16384 | 0 | 1.13666419 |
| S. syndactylus | JAMBI | TruSeq | dedup with GC | Z | 0.22277994 | 0.0005836 | 381.735727 | 0 | 1.06957927 |
| S. syndactylus | JAMBI | TruSeq | dedup with GC | DR | -0.4646803 | 0.0001546 | -3005.7043 | 0 | 1.25305239 |
| S. syndactylus | JAMBI | TruSeq | dedup with GC | G4 | -0.3793219 | 0.00022176 | -1710.534 | 0 | 1.32546096 |
| S. syndactylus | JAMBI | TruSeq | dedup with GC | GC_content | -0.1983919 | 8.15E-05 | -2434.8957 | 0 | 1.32954479 |
| P. pygmaeus | AG05252 | PDAL-Seq | dedup with GC | Intercept | 8.06551761 | 4.23E-05 | 190868.985 | 0 | NA |
| P. pygmaeus | AG05252 | PDAL-Seq | dedup with GC | APR | -0.251212 | 0.00069803 | -359.88814 | 0 | 1.01973793 |
| P. pygmaeus | AG05252 | PDAL-Seq | dedup with GC | CRU | -2.3890754 | 0.0010225 | -2336.4983 | 0 | 1.03692678 |
| P. pygmaeus | AG05252 | PDAL-Seq | dedup with GC | TRI | 1.5039461 | 0.00055271 | 2721.02075 | 0 | 1.07939871 |
| P. pygmaeus | AG05252 | PDAL-Seq | dedup with GC | Z | -0.7365169 | 0.00059047 | -1247.3351 | 0 | 1.05950916 |
| P. pygmaeus | AG05252 | PDAL-Seq | dedup with GC | DR | 1.04775225 | 0.00010748 | 9748.10391 | 0 | 1.13725266 |
| P. pygmaeus | AG05252 | PDAL-Seq | dedup with GC | G4 | -2.684911 | 0.00025985 | -10332.56 | 0 | 1.32569275 |
| P. pygmaeus | AG05252 | PDAL-Seq | dedup with GC | GC_content | 2.51129586 | 0.00010079 | 24916.5641 | 0 | 1.35008525 |
| P. paniscus | KB14571 | PDAL-Seq | dedup with GC | Intercept | 7.93795985 | 3.85E-05 | 206041.068 | 0 | NA |
| P. paniscus | KB14571 | PDAL-Seq | dedup with GC | APR | -0.3645266 | 0.00064986 | -560.93149 | 0 | 1.02135085 |
| P. paniscus | KB14571 | PDAL-Seq | dedup with GC | CRU | -1.6219101 | 0.00090739 | -1787.4494 | 0 | 1.04649765 |
| P. paniscus | KB14571 | PDAL-Seq | dedup with GC | TRI | 2.11223647 | 0.00049246 | 4289.12538 | 0 | 1.10386686 |
| P. paniscus | KB14571 | PDAL-Seq | dedup with GC | Z | -0.0798995 | 0.00058124 | -137.46352 | 0 | 1.07060464 |
| P. paniscus | KB14571 | PDAL-Seq | dedup with GC | DR | 0.61098721 | 0.00013338 | 4580.71025 | 0 | 1.20734075 |
| P. paniscus | KB14571 | PDAL-Seq | dedup with GC | G4 | -2.7533663 | 0.00023557 | -11688.302 | 0 | 1.38373716 |
| P. paniscus | KB14571 | PDAL-Seq | dedup with GC | GC_content | 3.2359991 | 9.14E-05 | 35417.6245 | 0 | 1.40731045 |
| P. troglodytes | AG18354 | PDAL-Seq | dedup with GC | Intercept | 6.31108611 | 6.31E-05 | 100068.084 | 0 | NA |
| P. troglodytes | AG18354 | PDAL-Seq | dedup with GC | APR | -0.1629279 | 0.0010906 | -149.39343 | 0 | 1.02421062 |

| | | | | | | | | | |
|---|---|---|---|---|---|---|---|---|---|
| P. troglodytes | AG18354 | PDAL-Seq | dedup with GC | CRU | -2.6968119 | 0.0015862 | -1700.1717 | 0 | 1.03021648 |
| P. troglodytes | AG18354 | PDAL-Seq | dedup with GC | TRI | 1.99202437 | 0.00077503 | 2570.25782 | 0 | 1.08397831 |
| P. troglodytes | AG18354 | PDAL-Seq | dedup with GC | Z | 2.36613963 | 0.00068043 | 3477.39528 | 0 | 1.11791698 |
| P. troglodytes | AG18354 | PDAL-Seq | dedup with GC | DR | 1.28866161 | 0.00016475 | 7821.88431 | 0 | 1.22966939 |
| P. troglodytes | AG18354 | PDAL-Seq | dedup with GC | G4 | -2.7249445 | 0.00033192 | -8209.6425 | 0 | 1.40862907 |
| P. troglodytes | AG18354 | PDAL-Seq | dedup with GC | GC_content | 4.65877401 | 0.00014639 | 31824.3205 | 0 | 1.43994833 |
| G. gorilla | KB3781 | PDAL-Seq | dedup with GC | Intercept | 6.14536032 | 8.24E-05 | 74595.4052 | 0 | NA |
| G. gorilla | KB3781 | PDAL-Seq | dedup with GC | APR | -0.1506501 | 0.00140668 | -107.09641 | 0 | 1.02227232 |
| G. gorilla | KB3781 | PDAL-Seq | dedup with GC | CRU | -2.5311884 | 0.00211032 | -1199.4328 | 0 | 1.03142727 |
| G. gorilla | KB3781 | PDAL-Seq | dedup with GC | TRI | 1.96502743 | 0.00099881 | 1967.36894 | 0 | 1.12431222 |
| G. gorilla | KB3781 | PDAL-Seq | dedup with GC | Z | 0.29435301 | 0.00110734 | 265.818713 | 0 | 1.08154368 |
| G. gorilla | KB3781 | PDAL-Seq | dedup with GC | DR | 0.85184127 | 0.00025409 | 3352.50593 | 0 | 1.22513484 |
| G. gorilla | KB3781 | PDAL-Seq | dedup with GC | G4 | -2.7824969 | 0.0004749 | -5859.1395 | 0 | 1.39333849 |
| G. gorilla | KB3781 | PDAL-Seq | dedup with GC | GC_content | 3.89714917 | 0.00019348 | 20142.5106 | 0 | 1.41971503 |
| G. gorilla | KB3781 | TruSeq | dedup with GC | Intercept | 8.01762035 | 0.00010668 | 75158.6279 | 0 | NA |
| G. gorilla | KB3781 | TruSeq | dedup with GC | APR | 0.00588706 | 0.00164723 | 3.57391887 | 0.00035168 | 1.015864 |
| G. gorilla | KB3781 | TruSeq | dedup with GC | CRU | 0.26498455 | 0.00222419 | 119.137691 | 0 | 1.10964203 |
| G. gorilla | KB3781 | TruSeq | dedup with GC | TRI | -0.9131882 | 0.00162948 | -560.41705 | 0 | 1.12495626 |
| G. gorilla | KB3781 | TruSeq | dedup with GC | Z | -2.0475412 | 0.00207006 | -989.12202 | 0 | 1.05875441 |
| G. gorilla | KB3781 | TruSeq | dedup with GC | DR | -0.7227901 | 0.00049788 | -1451.7311 | 0 | 1.2718147 |
| G. gorilla | KB3781 | TruSeq | dedup with GC | G4 | -0.3186347 | 0.00078561 | -405.58853 | 0 | 1.2890742 |
| G. gorilla | KB3781 | TruSeq | dedup with GC | GC_content | -1.7211804 | 0.00026624 | -6464.8859 | 0 | 1.29399351 |
| P. paniscus | KB8711 | PDAL-Seq | dedup with GC | Intercept | 6.02911773 | 8.51E-05 | 70847.9314 | 0 | NA |
| P. paniscus | KB8711 | PDAL-Seq | dedup with GC | APR | -0.3813536 | 0.00146963 | -259.48953 | 0 | 1.02251361 |
| P. paniscus | KB8711 | PDAL-Seq | dedup with GC | CRU | -2.2659407 | 0.00212399 | -1066.8317 | 0 | 1.03596665 |
| P. paniscus | KB8711 | PDAL-Seq | dedup with GC | TRI | 1.83379823 | 0.00107916 | 1699.28965 | 0 | 1.09458614 |
| P. paniscus | KB8711 | PDAL-Seq | dedup with GC | Z | -0.6660392 | 0.00129736 | -513.3797 | 0 | 1.06110671 |
| P. paniscus | KB8711 | PDAL-Seq | dedup with GC | DR | 0.81567822 | 0.00027413 | 2975.56229 | 0 | 1.19274252 |

| | | | | | | | | | |
|---|---|---|---|---|---|---|---|---|---|
| P. paniscus | KB8711 | PDAL-Seq | dedup with GC | G4 | -2.2208523 | 0.00047074 | -4717.8256 | 0 | 1.42168074 |
| P. paniscus | KB8711 | PDAL-Seq | dedup with GC | GC_content | 3.92949447 | 0.0001996 | 19686.852 | 0 | 1.43445725 |
| H. sapiens | GM24385 | PDAL-Seq | dedup with GC | Intercept | 5.05899436 | 8.31E-05 | 60883.5486 | 0 | NA |
| H. sapiens | GM24385 | PDAL-Seq | dedup with GC | APR | 0.26340585 | 0.00146914 | 179.292992 | 0 | 1.02625867 |
| H. sapiens | GM24385 | PDAL-Seq | dedup with GC | CRU | -3.6805535 | 0.00232514 | -1582.9361 | 0 | 1.01909673 |
| H. sapiens | GM24385 | PDAL-Seq | dedup with GC | TRI | 0.72891007 | 0.00101699 | 716.732825 | 0 | 1.07062173 |
| H. sapiens | GM24385 | PDAL-Seq | dedup with GC | Z | 3.10280417 | 0.00077253 | 4016.41664 | 0 | 1.12159287 |
| H. sapiens | GM24385 | PDAL-Seq | dedup with GC | DR | 1.09800636 | 0.00018625 | 5895.33591 | 0 | 1.18919077 |
| H. sapiens | GM24385 | PDAL-Seq | dedup with GC | G4 | -3.4202574 | 0.00042131 | -8118.1214 | 0 | 1.44438104 |
| H. sapiens | GM24385 | PDAL-Seq | dedup with GC | GC_content | 6.28205576 | 0.00018945 | 33159.8165 | 0 | 1.50876345 |
| S. syndactylus | JAMBI | PDAL-Seq | dedup with GC | Intercept | 5.96739336 | 6.43E-05 | 92842.2025 | 0 | NA |
| S. syndactylus | JAMBI | PDAL-Seq | dedup with GC | APR | -0.8407849 | 0.00113551 | -740.44693 | 0 | 1.02425364 |
| S. syndactylus | JAMBI | PDAL-Seq | dedup with GC | CRU | -1.6079367 | 0.00171721 | -936.36806 | 0 | 1.02084952 |
| S. syndactylus | JAMBI | PDAL-Seq | dedup with GC | TRI | 1.48925541 | 0.00073317 | 2031.24624 | 0 | 1.13838387 |
| S. syndactylus | JAMBI | PDAL-Seq | dedup with GC | Z | 2.75622184 | 0.00057567 | 4787.81704 | 0 | 1.23081962 |
| S. syndactylus | JAMBI | PDAL-Seq | dedup with GC | DR | 1.06329425 | 0.00018604 | 5715.30869 | 0 | 1.34909614 |
| S. syndactylus | JAMBI | PDAL-Seq | dedup with GC | G4 | -3.3548077 | 0.00034883 | -9617.3319 | 0 | 1.41756782 |
| S. syndactylus | JAMBI | PDAL-Seq | dedup with GC | GC_content | 5.52298848 | 0.00014761 | 37416.5342 | 0 | 1.46864703 |
| P. troglodytes | AG18354 | TruSeq | dedup with GC | Intercept | 9.71253031 | 3.17E-05 | 306706.288 | 0 | NA |
| P. troglodytes | AG18354 | TruSeq | dedup with GC | APR | -0.0642172 | 0.00050307 | -127.65013 | 0 | 1.01829424 |
| P. troglodytes | AG18354 | TruSeq | dedup with GC | CRU | -0.8411122 | 0.00070805 | -1187.9334 | 0 | 1.07413314 |
| P. troglodytes | AG18354 | TruSeq | dedup with GC | TRI | -0.1672152 | 0.00047771 | -350.03835 | 0 | 1.10101588 |
| P. troglodytes | AG18354 | TruSeq | dedup with GC | Z | 0.03320085 | 0.00055632 | 59.6795733 | 0 | 1.06785812 |
| P. troglodytes | AG18354 | TruSeq | dedup with GC | DR | -0.4184566 | 0.00013597 | -3077.5097 | 0 | 1.23430218 |
| P. troglodytes | AG18354 | TruSeq | dedup with GC | G4 | -0.3960505 | 0.00020477 | -1934.1261 | 0 | 1.34306004 |
| P. troglodytes | AG18354 | TruSeq | dedup with GC | GC_content | 7.91E-07 | 7.76E-05 | 0.01020251 | 0.99185971 | 1.33754788 |
| S. syndactylus | Karenina | PDAL-Seq | dedup with GC | Intercept | 5.49142874 | 9.21E-05 | 59596.668 | 0 | NA |
| S. syndactylus | Karenina | PDAL-Seq | dedup with GC | APR | -0.0825109 | 0.00158978 | -51.90081 | 0 | 1.02415999 |

| | | | | | | | | | |
|---|---|---|---|---|---|---|---|---|---|
| S. syndactylus | Karenina | PDAL-Seq | dedup with GC | CRU | -1.7741759 | 0.0025156 | -705.27078 | 0 | 1.02206946 |
| S. syndactylus | Karenina | PDAL-Seq | dedup with GC | TRI | 1.77016587 | 0.00114543 | 1545.41147 | 0 | 1.12730981 |
| S. syndactylus | Karenina | PDAL-Seq | dedup with GC | Z | 0.117714 | 0.00130877 | 89.9425 | 0 | 1.08178779 |
| S. syndactylus | Karenina | PDAL-Seq | dedup with GC | DR | 0.35693371 | 0.00032293 | 1105.31001 | 0 | 1.21248603 |
| S. syndactylus | Karenina | PDAL-Seq | dedup with GC | G4 | -2.8964187 | 0.00051237 | -5652.968 | 0 | 1.42807173 |
| S. syndactylus | Karenina | PDAL-Seq | dedup with GC | GC_content | 5.01810123 | 0.00021302 | 23556.8575 | 0 | 1.46566482 |
| P. paniscus | KB8711 | TruSeq | dedup with GC | Intercept | 10.9952627 | 2.71E-05 | 405187.213 | 0 | NA |
| P. paniscus | KB8711 | TruSeq | dedup with GC | APR | 0.02890149 | 0.00041031 | 70.4377537 | 0 | 1.01471281 |
| P. paniscus | KB8711 | TruSeq | dedup with GC | CRU | -1.0303235 | 0.00053254 | -1934.7176 | 0 | 1.14683134 |
| P. paniscus | KB8711 | TruSeq | dedup with GC | TRI | 0.10998058 | 0.00040415 | 272.129463 | 0 | 1.11596937 |
| P. paniscus | KB8711 | TruSeq | dedup with GC | Z | -0.4252296 | 0.00049651 | -856.44559 | 0 | 1.06461416 |
| P. paniscus | KB8711 | TruSeq | dedup with GC | DR | -0.2476139 | 0.00011878 | -2084.6257 | 0 | 1.28520846 |
| P. paniscus | KB8711 | TruSeq | dedup with GC | G4 | -2.7820692 | 0.00024024 | -11580.219 | 0 | 1.22066606 |
| P. paniscus | KB8711 | TruSeq | dedup with GC | GC_content | -2.3264851 | 6.84E-05 | -34008.01 | 0 | 1.24590811 |

**Table S6:** Homologuous relationships among ape chromosomes identified by alignment filtering criteria.

| H. sapiens (ref) | P. troglodytes | P. paniscus | G. gorilla | P. pygmaeus | P. abelii | S. syndactylus |
|---|---|---|---|---|---|---|
| chr1 | chr1_hap1_hsa1 | chr1_pat_hsa1 | chr1_pat_hsa1 | chr1_hap1_hsa1 | chr1_hap1_hsa1 | chr12_hap1, chr19_hap1 |
| chr2 | chr13_hap1_hsa2b, chr12_hap1_hsa2a | chr13_mat_hsa2b, chr12_mat_hsa2a | chr11_mat_hsa2b, chr12_pat_hsa2a | chr11_hap1_hsa2b, chr12_hap1_hsa2a | chr11_hap1_hsa2b, chr12_hap1_hsa2a | chr8_hap1, chr14_hap1, chr18_hap1, chr22_hap1 |
| chr3 | chr2_hap1_hsa3 | chr2_mat_hsa3 | chr2_pat_hsa3 | chr2_hap1_hsa3 | chr2_hap1_hsa3 | chr21_hap1, chr17_hap1, chr1_hap1 |
| chr4 | chr3_hap1_hsa4 | chr3_pat_hsa4 | chr3_pat_hsa4 | chr3_hap1_hsa4 | chr3_hap1_hsa4 | chr4_hap1, chr10_hap1, chr16_hap1 |
| chr5 | chr4_hap1_hsa5 | chr4_mat_hsa5 | chr4_pat_hsa17x5, chr19_pat_hsa5x17 | chr4_hap1_hsa5 | chr4_hap1_hsa5 | chr11_hap1, chr7_hap1, chr16_hap1 |
| chr6 | chr5_hap1_hsa6 | chr5_pat_hsa6 | chr5_mat_hsa6 | chr5_hap1_hsa6 | chr5_hap1_hsa6 | chr2_hap1, chr23_hap1 |
| chr7 | chr6_hap1_hsa7 | chr6_mat_hsa7 | chr6_mat_hsa7 | chr6_hap1_hsa7 | chr6_hap1_hsa7 | chr6_hap1, chr9_hap1, chr3_hap1 |
| chr8 | chr7_hap1_hsa8 | chr7_pat_hsa8 | chr7_pat_hsa8 | chr7_hap1_hsa8 | chr7_hap1_hsa8 | chr7_hap1, chr10_hap1 |
| chr9 | chr11_hap1_hsa9 | chr11_mat_hsa9 | chr13_pat_hsa9 | chr13_hap1_hsa9 | chr13_hap1_hsa9 | chr3_hap1, chr9_hap1 |
| chr10 | chr8_hap1_hsa10 | chr8_mat_hsa10 | chr8_pat_hsa10 | chr8_hap1_hsa10 | chr8_hap1_hsa10 | chr4_hap1, chr2_hap1 |
| chr11 | chr9_hap1_hsa11 | chr9_pat_hsa11 | chr9_pat_hsa11 | chr9_hap1_hsa11 | chr9_hap1_hsa11 | chr6_hap1, chr3_hap1 |
| chr12 | chr10_hap1_hsa12 | chr10_mat_hsa12 | chr10_mat_hsa12 | chr10_hap1_hsa12 | chr10_hap1_hsa12 | chr13_hap1, chr5_hap1, chr17_hap1 |
| chr13 | chr14_hap1_hsa13 | chr14_pat_hsa13 | chr14_pat_hsa13 | chr14_hap1_hsa13 | chr14_hap1_hsa13 | chr15_hap1 |

**Table S6:** Homologuous relationships among ape chromosomes identified by alignment filtering criteria.

| H. sapiens (ref) | P. troglodytes | P. paniscus | G. gorilla | P. pygmaeus | P. abelii | S. syndactylus |
|---|---|---|---|---|---|---|
| chr14 | chr15_hap1_hsa14 | chr15_mat_hsa14 | chr15_pat_hsa14 | chr15_hap1_hsa14 | chr15_hap1_hsa14 | chr8_hap1, chr9_hap1 |
| chr15 | chr16_hap1_hsa15 | chr16_pat_hsa15 | chr16_pat_hsa15 | chr16_hap1_hsa15 | chr16_hap1_hsa15 | chr5_hap1 |
| chr16 | chr18_hap1_hsa16 | chr18_pat_hsa16 | chr18_pat_hsa16 | chr18_hap1_hsa16 | chr18_hap1_hsa16 | chr11_hap1, chr14_hap1 |
| chr17 | chr19_hap1_hsa17 | chr19_pat_hsa17 | chr4_pat_hsa17x5, chr19_pat_hsa5x17 | chr19_hap1_hsa17 | chr19_hap1_hsa17 | chr20_hap1, chr14_hap1 |
| chr18 | chr17_hap1_hsa18 | chr17_mat_hsa18 | chr17_mat_hsa18 | chr17_hap1_hsa18 | chr17_hap1_hsa18 | chr1_hap1 |
| chr19 | chr20_hap1_hsa19 | chr20_pat_hsa19 | chr20_mat_hsa19 | chr20_hap1_hsa19 | chr20_hap1_hsa19 | chr13_hap1, chr17_hap1 |
| chr20 | chr21_hap1_hsa20 | chr21_mat_hsa20 | chr21_pat_hsa20 | chr21_hap1_hsa20 | chr21_hap1_hsa20 | chr24_hap1 |
| chr21 | chr22_hap1_hsa21 | chr22_pat_hsa21 | chr22_mat_hsa21 | chr22_hap1_hsa21 | chr22_hap1_hsa21 | chr5_hap1 |
| chr22 | chr23_hap1_hsa22 | chr23_pat_hsa22 | chr23_mat_hsa22 | chr23_hap1_hsa22 | chr23_hap1_hsa22 | chr18_hap1 |

**Table S7:** Alignment filtering statistics.

Column description:

Species group: The species group used to determine alignment filtering parameters.

Total alignment blocks: number of alignment blocks that passed alignment filtering criteria.

Blocks after book-ended merge: Alignment blocks that passed alignment filtering criteria were merged if they overlap or are immediatey adjacent to each other. A perfect alignment would have 1 block after a book-ended merge because it has no gaps or structural rearrangements.

Fraction of blocks that are continuous: Blocks that are immediately adjacent to each other in linear genomic coordinates. They were calculated by subtracting merged blocks from total blocks and dividing by total blocks.

Base pairs in total alignments: Base pairs in the alignment blocks that passed filtering criteria.

Base pairs after book-ended merge: Base pairs in the alignment blocks that passed filtering criteria, after overlapping or immediately adjacent blocks are merged. A perfect alignment would have the same number of basepairs before and after blocks are merged, where base pairs that are deleted in the merge show up more than once in the alignment (duplication rate)

BP in genome: Total base pairs in each primate T2T genome.

Genome coverage: Amount of the T2T genome covered by alignment blocks that passed alignment filtering criteria.

Duplication rate: Percent of bases that are represented more than once in the alignment blocks that passed alignment filtering criterias.

| Species group | Species | Total alignment blocks | Blocks after book-ended merge | Fraction of blocks that are continuous | Base pair in total alignments | Base pair after book-ended merge | BP in genome | % Genome coverage | Duplication rate | Median alignment block length (base pair) |
|---|---|---|---|---|---|---|---|---|---|---|
| *Hominoidae* | *H. sapiens* | 3502282 | 190564 | 94.6% | 2197182775 | 2193871057 | 3117275501 | 70% | 0.2% | 517 |
| *Hominoidae* | *P. troglodytes* | 3502282 | 860006 | 75.4% | 2188172566 | 2170970234 | 3177739762 | 69% | 0.8% | 515 |
| *Hominoidae* | *P. paniscus* | 3502282 | 851066 | 75.7% | 2187702860 | 2170436891 | 3244888708 | 67% | 0.8% | 515 |
| *Hominoidae* | *G. gorilla* | 3502282 | 1020547 | 70.9% | 2186865502 | 2168220484 | 3545834224 | 62% | 0.9% | 515 |
| *Hominoidae* | *P. pygmaeus* | 3502282 | 1831042 | 47.7% | 2177845310 | 2156365377 | 3220935163 | 68% | 1.0% | 513 |
| *Hominoidae* | *P. abelii* | 3502282 | 1830157 | 47.7% | 2177950252 | 2156475974 | 3259853530 | 67% | 1.0% | 513 |
| *Hominoidae* | *S. syndactylus* | 3502282 | 2136771 | 39.0% | 2168873431 | 2145295961 | 3262887708 | 66% | 1.1% | 511 |
| *Hominidae* | *H. sapiens* | 3629915 | 210843 | 94.2% | 2272908970 | 2269489898 | 3117275501 | 73% | 0.2% | 514 |
| *Hominidae* | *P. troglodytes* | 3629915 | 907421 | 75.0% | 2263485997 | 2244744455 | 3177739762 | 71% | 0.8% | 512 |
| *Hominidae* | *P. paniscus* | 3629915 | 898186 | 75.3% | 2263001512 | 2244215011 | 3244888708 | 70% | 0.8% | 512 |
| *Hominidae* | *G. gorilla* | 3629915 | 1073991 | 70.4% | 2262111768 | 2241779092 | 3545834224 | 64% | 0.9% | 512 |
| *Hominidae* | *P. pygmaeus* | 3629915 | 1917024 | 47.2% | 2252654466 | 2229242458 | 3220935163 | 70% | 1.0% | 511 |
| *Hominidae* | *P. abelii* | 3629915 | 1916256 | 47.2% | 2252761852 | 2229363354 | 3259853530 | 69% | 1.0% | 511 |
| *Homininae* | *H. sapiens* | 3688745 | 224318 | 93.9% | 2308275096 | 2304810669 | 3117275501 | 74% | 0.2% | 515 |
| *Homininae* | *P. troglodytes* | 3688745 | 933992 | 74.7% | 2298596674 | 2277809600 | 3177739762 | 72% | 0.9% | 513 |
| *Homininae* | *P. paniscus* | 3688745 | 924772 | 74.9% | 2298101737 | 2277325313 | 3244888708 | 71% | 0.9% | 512 |
| *Homininae* | *G. gorilla* | 3688745 | 1103270 | 70.1% | 2297096400 | 2274359040 | 3545834224 | 65% | 1.0% | 512 |
| *Hominini* | *H. sapiens* | 3721099 | 231874 | 93.8% | 2325924568 | 2322435343 | 3117275501 | 75% | 0.2% | 516 |
| *Hominini* | *P. troglodytes* | 3721099 | 948405 | 74.5% | 2316037201 | 2294334752 | 3177739762 | 73% | 0.9% | 514 |
| *Hominini* | *P. paniscus* | 3721099 | 939239 | 74.8% | 2315526230 | 2293863564 | 3244888708 | 71% | 0.9% | 514 |

**Table S8:** Unique mapping rates in syntenic regions of the genome. Uniquely mapping alignments are alignments where the primary mapping location is more than 10 times better than the secondary mapping location.

| Species group | Species | Cell line | Technique | Percent unique mapping reads in alignment bl |
|---|---|---|---|---|
| Hominoidae | H. sapiens | GM24385 | PDAL-Seq | 95.0% |
| Hominoidae | H. sapiens | GM24143 | PDAL-Seq | 94.6% |
| Hominoidae | H. sapiens | HEK293T | PDAL-Seq | 88.2% |
| Hominoidae | H. sapiens | CCL86-Raji | PDAL-Seq | 93.1% |
| Hominoidae | P. troglodytes | AG18354 | PDAL-Seq | 92.7% |
| Hominoidae | P. troglodytes | AG18358 | PDAL-Seq | 96.5% |
| Hominoidae | P. paniscus | KB8711 | PDAL-Seq | 97.9% |
| Hominoidae | P. paniscus | KB14571 | PDAL-Seq | 97.3% |
| Hominoidae | G. gorilla | KB3781 | PDAL-Seq | 95.1% |
| Hominoidae | G. gorilla | F7549 | PDAL-Seq | 96.7% |
| Hominoidae | P. pygmaeus | AG05252 | PDAL-Seq | 98.0% |
| Hominoidae | P. abelii | AG06213 | PDAL-Seq | 97.9% |
| Hominoidae | S. syndactylus | JAMBI | PDAL-Seq | 91.4% |
| Hominoidae | S. syndactylus | Karenina | PDAL-Seq | 97.5% |
| Hominoidae | H. sapiens | GM24385 | TruSeq | 98.4% |
| Hominoidae | P. troglodytes | AG18354 | TruSeq | 99.1% |
| Hominoidae | P. paniscus | KB8711 | TruSeq | 99.0% |
| Hominoidae | G. gorilla | KB3781 | TruSeq | 99.0% |
| Hominoidae | P. pygmaeus | AG05252 | TruSeq | 99.2% |
| Hominoidae | P. abelii | AG06213 | TruSeq | 99.2% |

**Table S8:** Unique mapping rates in syntenic regions of the genome. Uniquely mapping alignments are alignments where the primary mapping location is more than 10 times better than the secondary mapping location.

| Species group | Species | Cell line | Technique | Percent unique mapping reads in alignment bl |
|---|---|---|---|---|
| Hominoidae | S. syndactylus | JAMBI | TruSeq | 99.3% |
| Hominidae | H. sapiens | GM24385 | PDAL-Seq | 94.8% |
| Hominidae | H. sapiens | GM24143 | PDAL-Seq | 94.3% |
| Hominidae | H. sapiens | HEK293T | PDAL-Seq | 87.8% |
| Hominidae | H. sapiens | CCL86-Raji | PDAL-Seq | 92.8% |
| Hominidae | P. troglodytes | AG18354 | PDAL-Seq | 92.4% |
| Hominidae | P. troglodytes | AG18358 | PDAL-Seq | 96.3% |
| Hominidae | P. paniscus | KB8711 | PDAL-Seq | 97.8% |
| Hominidae | P. paniscus | KB14571 | PDAL-Seq | 97.2% |
| Hominidae | G. gorilla | KB3781 | PDAL-Seq | 95.0% |
| Hominidae | G. gorilla | F7549 | PDAL-Seq | 96.6% |
| Hominidae | P. pygmaeus | AG05252 | PDAL-Seq | 97.9% |
| Hominidae | P. abelii | AG06213 | PDAL-Seq | 97.7% |
| Hominidae | H. sapiens | GM24385 | TruSeq | 98.3% |
| Hominidae | P. troglodytes | AG18354 | TruSeq | 99.0% |
| Hominidae | P. paniscus | KB8711 | TruSeq | 98.9% |
| Hominidae | G. gorilla | KB3781 | TruSeq | 98.9% |
| Hominidae | P. pygmaeus | AG05252 | TruSeq | 99.1% |
| Hominidae | P. abelii | AG06213 | TruSeq | 99.1% |

**Table S8:** Unique mapping rates in syntenic regions of the genome. Uniquely mapping alignments are alignments where the primary mapping location is more than 10 times better than the secondary mapping location.

| Species group | Species | Cell line | Technique | Percent unique mapping reads in alignment bl |
|---|---|---|---|---|
| Homininae | H. sapiens | GM24385 | PDAL-Seq | 94.2% |
| Homininae | H. sapiens | GM24143 | PDAL-Seq | 93.8% |
| Homininae | H. sapiens | HEK293T | PDAL-Seq | 87.0% |
| Homininae | H. sapiens | CCL86-Raji | PDAL-Seq | 92.0% |
| Homininae | P. troglodytes | AG18354 | PDAL-Seq | 91.8% |
| Homininae | P. troglodytes | AG18358 | PDAL-Seq | 95.9% |
| Homininae | P. paniscus | KB8711 | PDAL-Seq | 97.2% |
| Homininae | P. paniscus | KB14571 | PDAL-Seq | 96.6% |
| Homininae | G. gorilla | KB3781 | PDAL-Seq | 94.5% |
| Homininae | G. gorilla | F7549 | PDAL-Seq | 96.1% |
| Homininae | H. sapiens | GM24385 | TruSeq | 97.9% |
| Homininae | P. troglodytes | AG18354 | TruSeq | 98.7% |
| Homininae | P. paniscus | KB8711 | TruSeq | 98.7% |
| Homininae | G. gorilla | KB3781 | TruSeq | 98.6% |
| Hominini | H. sapiens | GM24385 | PDAL-Seq | 94.0% |
| Hominini | H. sapiens | GM24143 | PDAL-Seq | 93.6% |
| Hominini | H. sapiens | HEK293T | PDAL-Seq | 86.7% |
| Hominini | H. sapiens | CCL86-Raji | PDAL-Seq | 91.7% |
| Hominini | P. troglodytes | AG18354 | PDAL-Seq | 91.6% |

**Table S8:** Unique mapping rates in syntenic regions of the genome. Uniquely mapping alignments are alignments where the primary mapping location is more than 10 times better than the secondary mapping location.

| Species group | Species | Cell line | Technique | Percent unique mapping reads in alignment bl |
|---|---|---|---|---|
| Hominini | P. troglodytes | AG18358 | PDAL-Seq | 95.7% |
| Hominini | P. paniscus | KB8711 | PDAL-Seq | 97.0% |
| Hominini | P. paniscus | KB14571 | PDAL-Seq | 96.3% |
| Hominini | H. sapiens | GM24385 | TruSeq | 97.7% |
| Hominini | P. troglodytes | AG18354 | TruSeq | 98.6% |
| Hominini | P. paniscus | KB8711 | TruSeq | 98.6% |

**Table S9:** Pairwise correlations between sequencing datasets in multi-species alignments. Pairwise correlation coefficients were calculated for PDAL-Seq or TruSeq signal in the 7-way alignments preprocessed using Hominoidae as a species group.

| Sample 1 | Sample 2 | # Pearson correlation coefficient | # Spearman's correlation coefficient |
|---|---|---|---|
| H. sapiens_GM24385_PDAL-Seq | H. sapiens_GM24143_PDAL-Seq | 0.77327306 | 0.76208642 |
| H. sapiens_GM24385_PDAL-Seq | P. troglodytes_AG18354_PDAL-Seq | 0.72299247 | 0.72981992 |
| H. sapiens_GM24143_PDAL-Seq | P. troglodytes_AG18354_PDAL-Seq | 0.74254915 | 0.76042013 |
| H. sapiens_GM24385_PDAL-Seq | P. troglodytes_AG18358_PDAL-Seq | 0.60450563 | 0.62431267 |
| H. sapiens_GM24143_PDAL-Seq | P. troglodytes_AG18358_PDAL-Seq | 0.62750516 | 0.64506195 |
| P. troglodytes_AG18354_PDAL-Seq | P. troglodytes_AG18358_PDAL-Seq | 0.71059393 | 0.67428021 |
| H. sapiens_GM24385_PDAL-Seq | P. paniscus_KB8711_PDAL-Seq | 0.51552898 | 0.55030171 |
| H. sapiens_GM24143_PDAL-Seq | P. paniscus_KB8711_PDAL-Seq | 0.55349448 | 0.57493309 |
| P. troglodytes_AG18354_PDAL-Seq | P. paniscus_KB8711_PDAL-Seq | 0.61292 | 0.58822153 |
| P. troglodytes_AG18358_PDAL-Seq | P. paniscus_KB8711_PDAL-Seq | 0.55418104 | 0.51835309 |
| H. sapiens_GM24385_PDAL-Seq | P. paniscus_KB14571_PDAL-Seq | 0.5640653 | 0.66053289 |
| H. sapiens_GM24143_PDAL-Seq | P. paniscus_KB14571_PDAL-Seq | 0.60226324 | 0.69115527 |
| P. troglodytes_AG18354_PDAL-Seq | P. paniscus_KB14571_PDAL-Seq | 0.72884322 | 0.724426 |
| P. troglodytes_AG18358_PDAL-Seq | P. paniscus_KB14571_PDAL-Seq | 0.66483584 | 0.6391966 |
| P. paniscus_KB8711_PDAL-Seq | P. paniscus_KB14571_PDAL-Seq | 0.66683034 | 0.60478393 |
| H. sapiens_GM24385_PDAL-Seq | G. gorilla_KB3781_PDAL-Seq | 0.54343715 | 0.58584532 |
| H. sapiens_GM24143_PDAL-Seq | G. gorilla_KB3781_PDAL-Seq | 0.57430462 | 0.61039342 |
| P. troglodytes_AG18354_PDAL-Seq | G. gorilla_KB3781_PDAL-Seq | 0.65564697 | 0.62479255 |
| P. troglodytes_AG18358_PDAL-Seq | G. gorilla_KB3781_PDAL-Seq | 0.58600533 | 0.54238614 |
| P. paniscus_KB8711_PDAL-Seq | G. gorilla_KB3781_PDAL-Seq | 0.55095842 | 0.50123735 |
| P. paniscus_KB14571_PDAL-Seq | G. gorilla_KB3781_PDAL-Seq | 0.66396265 | 0.60323587 |
| H. sapiens_GM24385_PDAL-Seq | G. gorilla_F7549_PDAL-Seq | 0.67284706 | 0.70598266 |
| H. sapiens_GM24143_PDAL-Seq | G. gorilla_F7549_PDAL-Seq | 0.69650289 | 0.72705389 |
| P. troglodytes_AG18354_PDAL-Seq | G. gorilla_F7549_PDAL-Seq | 0.73224078 | 0.72824632 |
| P. troglodytes_AG18358_PDAL-Seq | G. gorilla_F7549_PDAL-Seq | 0.64906016 | 0.63289787 |
| P. paniscus_KB8711_PDAL-Seq | G. gorilla_F7549_PDAL-Seq | 0.60031781 | 0.57952579 |
| P. paniscus_KB14571_PDAL-Seq | G. gorilla_F7549_PDAL-Seq | 0.70867935 | 0.70554656 |
| G. gorilla_KB3781_PDAL-Seq | G. gorilla_F7549_PDAL-Seq | 0.66440106 | 0.61676748 |
| H. sapiens_GM24385_PDAL-Seq | P. pygmaeus_AG05252_PDAL-Seq | 0.48789672 | 0.55850127 |
| H. sapiens_GM24143_PDAL-Seq | P. pygmaeus_AG05252_PDAL-Seq | 0.52061553 | 0.58212785 |
| P. troglodytes_AG18354_PDAL-Seq | P. pygmaeus_AG05252_PDAL-Seq | 0.62326744 | 0.61568991 |
| P. troglodytes_AG18358_PDAL-Seq | P. pygmaeus_AG05252_PDAL-Seq | 0.57485051 | 0.5496599 |
| P. paniscus_KB8711_PDAL-Seq | P. pygmaeus_AG05252_PDAL-Seq | 0.5792393 | 0.54044154 |
| P. paniscus_KB14571_PDAL-Seq | P. pygmaeus_AG05252_PDAL-Seq | 0.69946893 | 0.65150827 |
| G. gorilla_KB3781_PDAL-Seq | P. pygmaeus_AG05252_PDAL-Seq | 0.58811569 | 0.52540441 |

**Table S9:** Pairwise correlations between sequencing datasets in multi-species alignments. Pairwise correlation coefficients were calculated for PDAL-Seq or TruSeq signal in the 7-way alignments preprocessed using Hominoidae as a species group.

| Sample 1 | Sample 2 | # Pearson correlation coefficient | # Spearman's correlation coefficient |
|---|---|---|---|
| G. gorilla_F7549_PDAL-Seq | P. pygmaeus_AG05252_PDAL-Seq | 0.61273457 | 0.60577555 |
| H. sapiens_GM24385_PDAL-Seq | P. abelii_AG06213_PDAL-Seq | 0.50230073 | 0.5575477 |
| H. sapiens_GM24143_PDAL-Seq | P. abelii_AG06213_PDAL-Seq | 0.52903226 | 0.57717972 |
| P. troglodytes_AG18354_PDAL-Seq | P. abelii_AG06213_PDAL-Seq | 0.60326032 | 0.59290949 |
| P. troglodytes_AG18358_PDAL-Seq | P. abelii_AG06213_PDAL-Seq | 0.55125223 | 0.52903043 |
| P. paniscus_KB8711_PDAL-Seq | P. abelii_AG06213_PDAL-Seq | 0.51726127 | 0.4810065 |
| P. paniscus_KB14571_PDAL-Seq | P. abelii_AG06213_PDAL-Seq | 0.64492294 | 0.59607442 |
| G. gorilla_KB3781_PDAL-Seq | P. abelii_AG06213_PDAL-Seq | 0.55178639 | 0.49545208 |
| G. gorilla_F7549_PDAL-Seq | P. abelii_AG06213_PDAL-Seq | 0.59965453 | 0.58438396 |
| P. pygmaeus_AG05252_PDAL-Seq | P. abelii_AG06213_PDAL-Seq | 0.66843198 | 0.55177901 |
| H. sapiens_GM24385_PDAL-Seq | S. syndactylus_JAMBI_PDAL-Seq | 0.70984581 | 0.72859097 |
| H. sapiens_GM24143_PDAL-Seq | S. syndactylus_JAMBI_PDAL-Seq | 0.72187112 | 0.75963808 |
| P. troglodytes_AG18354_PDAL-Seq | S. syndactylus_JAMBI_PDAL-Seq | 0.75965646 | 0.76099303 |
| P. troglodytes_AG18358_PDAL-Seq | S. syndactylus_JAMBI_PDAL-Seq | 0.64060715 | 0.64685082 |
| P. paniscus_KB8711_PDAL-Seq | S. syndactylus_JAMBI_PDAL-Seq | 0.53935838 | 0.56897855 |
| P. paniscus_KB14571_PDAL-Seq | S. syndactylus_JAMBI_PDAL-Seq | 0.64269371 | 0.69794094 |
| G. gorilla_KB3781_PDAL-Seq | S. syndactylus_JAMBI_PDAL-Seq | 0.60410478 | 0.61554008 |
| G. gorilla_F7549_PDAL-Seq | S. syndactylus_JAMBI_PDAL-Seq | 0.69205799 | 0.72111041 |
| P. pygmaeus_AG05252_PDAL-Seq | S. syndactylus_JAMBI_PDAL-Seq | 0.56541599 | 0.59602535 |
| P. abelii_AG06213_PDAL-Seq | S. syndactylus_JAMBI_PDAL-Seq | 0.56186251 | 0.58412305 |
| H. sapiens_GM24385_PDAL-Seq | S. syndactylus_Karenina_PDAL-Seq | 0.57026769 | 0.60357544 |
| H. sapiens_GM24143_PDAL-Seq | S. syndactylus_Karenina_PDAL-Seq | 0.59938185 | 0.62245562 |
| P. troglodytes_AG18354_PDAL-Seq | S. syndactylus_Karenina_PDAL-Seq | 0.60424476 | 0.61988478 |
| P. troglodytes_AG18358_PDAL-Seq | S. syndactylus_Karenina_PDAL-Seq | 0.54720352 | 0.54670893 |
| P. paniscus_KB8711_PDAL-Seq | S. syndactylus_Karenina_PDAL-Seq | 0.48450794 | 0.48562456 |
| P. paniscus_KB14571_PDAL-Seq | S. syndactylus_Karenina_PDAL-Seq | 0.56784946 | 0.59249932 |
| G. gorilla_KB3781_PDAL-Seq | S. syndactylus_Karenina_PDAL-Seq | 0.50917559 | 0.50678963 |
| G. gorilla_F7549_PDAL-Seq | S. syndactylus_Karenina_PDAL-Seq | 0.60383382 | 0.60810674 |
| P. pygmaeus_AG05252_PDAL-Seq | S. syndactylus_Karenina_PDAL-Seq | 0.49570672 | 0.50818658 |
| P. abelii_AG06213_PDAL-Seq | S. syndactylus_Karenina_PDAL-Seq | 0.5014799 | 0.50618285 |
| S. syndactylus_JAMBI_PDAL-Seq | S. syndactylus_Karenina_PDAL-Seq | 0.62040059 | 0.63833657 |

**Table S10:** Human state 8 gene enrichment analysis from Gene Ontology (GO) biological process database 2021 and Kyoto Encyclopedia of Genes and Genomes (KEGG) databases. State 9 p-values test the null hypothesis that the same pathways would be observed by randomly sampling human state 9.

Column descriptions:

adj_p_val: The adjusted p-value for the Fisher's exact test was provided by the EnrichR program.

num_genes: number of genes overlapping with each pathway

Obs.state9: Number of times the pathway is detected as enriched by subsampling Human state 9 (chosen because it is gene rich and has consistently high PDAL-Seq signal)

P.stat9: Probability of detecting this pathway as enriched by subsampling Human state 9

| GO database | | | | | | KEGG database | | | | | |
|---|---|---|---|---|---|---|---|---|---|---|---|
| path_name | adj_p_val | num_genes | Data_base | Obs.state9 | p.state9 | path_name | adj_p_val | num_genes | Data_base | Obs.state9 | p.state9 |
| regulation of small GTPase mediated signal tra | 0.00042962 | 56 | GO | 0 | 0 | Human immunodeficiency virus 1 infection | 0.02295982 | 62 | KEGG | 0 | 0 |
| positive regulation of synaptic transmission (G | 0.0009207 | 34 | GO | 0 | 0 | Glucagon signaling pathway | 0.01410424 | 36 | KEGG | 1 | 0.01 |
| regulation of neurotransmitter receptor activity | 0.00221893 | 28 | GO | 0 | 0 | Retrograde endocannabinoid signaling | 0.02128441 | 46 | KEGG | 1 | 0.01 |
| regulation of stress fiber assembly (GO:00514 | 0.00254175 | 33 | GO | 0 | 0 | HIF-1 signaling pathway | 0.03012874 | 35 | KEGG | 2 | 0.02 |
| calcium ion transmembrane transport (GO:007 | 0.00263407 | 37 | GO | 0 | 0 | Human T-cell leukemia virus 1 infection | 0.03012874 | 63 | KEGG | 2 | 0.02 |
| neuron projection development (GO:0031175) | 0.00411642 | 61 | GO | 0 | 0 | Serotonergic synapse | 0.03040746 | 36 | KEGG | 2 | 0.02 |
| regulation of Ras protein signal transduction (( | 0.00473066 | 36 | GO | 0 | 0 | Long-term potentiation | 0.00153742 | 28 | KEGG | 4 | 0.04 |
| neuron development (GO:0048666) | 0.00815221 | 52 | GO | 0 | 0 | Choline metabolism in cancer | 0.00167713 | 37 | KEGG | 4 | 0.04 |
| peptidyl-serine phosphorylation (GO:0018105) | 0.0163682 | 54 | GO | 0 | 0 | Adherens junction | 0.01416273 | 26 | KEGG | 5 | 0.05 |
| homophilic cell adhesion via plasma membrane | 0.01803675 | 26 | GO | 0 | 0 | Longevity regulating pathway | 0.03097038 | 33 | KEGG | 5 | 0.05 |
| receptor-mediated endocytosis (GO:0006898) | 0.0190289 | 50 | GO | 0 | 0 | Neurotrophin signaling pathway | 0.03750504 | 37 | KEGG | 5 | 0.05 |
| neuron projection morphogenesis (GO:004881 | 0.02041292 | 49 | GO | 0 | 0 | Pancreatic secretion | 0.03097038 | 33 | KEGG | 6 | 0.06 |
| modulation of chemical synaptic transmission ( | 0.02511389 | 40 | GO | 0 | 0 | Yersinia infection | 0.04793726 | 41 | KEGG | 6 | 0.06 |
| cell morphogenesis involved in differentiation | 0.02709001 | 29 | GO | 0 | 0 | Colorectal cancer | 0.04419475 | 28 | KEGG | 7 | 0.07 |
| plasma membrane bounded cell projection orga | 0.02766244 | 45 | GO | 0 | 0 | Fc gamma R-mediated phagocytosis | 0.00150642 | 37 | KEGG | 9 | 0.09 |
| peptidyl-serine modification (GO:0018209) | 0.02916296 | 56 | GO | 0 | 0 | Leukocyte transendothelial migration | 0.03250077 | 36 | KEGG | 11 | 0.11 |
| regulation of intracellular signal transduction (( | 0.04059517 | 123 | GO | 0 | 0 | VEGF signaling pathway | 0.0009868 | 26 | KEGG | 13 | 0.13 |
| cellular metal ion homeostasis (GO:0006875) | 0.04977684 | 37 | GO | 0 | 0 | Chemokine signaling pathway | 0.02295982 | 57 | KEGG | 14 | 0.14 |
| chloride transport (GO:0006821) | 0.04977684 | 29 | GO | 0 | 0 | Insulin resistance | 0.000412 | 42 | KEGG | 15 | 0.15 |
| collagen fibril organization (GO:0030199) | 0.00221893 | 38 | GO | 1 | 0.01 | Thyroid hormone signaling pathway | 0.00794813 | 41 | KEGG | 15 | 0.15 |
| protein localization to membrane (GO:007265 | 0.01440414 | 65 | GO | 1 | 0.01 | GABAergic synapse | 0.00023679 | 37 | KEGG | 16 | 0.16 |
| peptidyl-tyrosine modification (GO:0018212) | 0.01574271 | 25 | GO | 1 | 0.01 | Relaxin signaling pathway | 0.00153952 | 46 | KEGG | 16 | 0.16 |
| protein localization to plasma membrane (GO:( | 0.01933112 | 48 | GO | 1 | 0.01 | Human cytomegalovirus infection | 0.03306997 | 64 | KEGG | 16 | 0.16 |
| regulation of neuron projection development (( | 0.0285964 | 55 | GO | 1 | 0.01 | PI3K-Akt signaling pathway | 0.00162431 | 105 | KEGG | 17 | 0.17 |
| protein localization to cell periphery (GO:1990 | 0.03050436 | 48 | GO | 1 | 0.01 | ECM-receptor interaction | 0.00741695 | 32 | KEGG | 19 | 0.19 |
| activation of GTPase activity (GO:0090630) | 0.03443807 | 38 | GO | 1 | 0.01 | Insulin signaling pathway | 0.01410424 | 44 | KEGG | 23 | 0.23 |
| sensory perception of mechanical stimulus (GC | 0.03608726 | 33 | GO | 1 | 0.01 | cGMP-PKG signaling pathway | 0.01343773 | 52 | KEGG | 24 | 0.24 |
| phosphorylation (GO:0016310) | 0.000000382 | 140 | GO | 2 | 0.02 | Platelet activation | 0.01847891 | 40 | KEGG | 24 | 0.24 |
| calcium ion transport (GO:0006816) | 0.00103663 | 47 | GO | 2 | 0.02 | Morphine addiction | 0.000018 | 41 | KEGG | 28 | 0.28 |
| ion transport (GO:0006811) | 0.00250119 | 46 | GO | 2 | 0.02 | Endocytosis | 0.00590071 | 76 | KEGG | 30 | 0.3 |
| regulation of cytoskeleton organization (GO:0( | 0.00837378 | 43 | GO | 2 | 0.02 | GnRH secretion | 0.00000273 | 34 | KEGG | 32 | 0.32 |
| positive regulation of cell migration (GO:0030 | 0.01706136 | 84 | GO | 2 | 0.02 | Non-small cell lung cancer | 0.03041319 | 25 | KEGG | 33 | 0.33 |
| negative regulation of cellular process (GO:00 | 0.02916296 | 156 | GO | 2 | 0.02 | Hepatocellular carcinoma | 0.00211419 | 56 | KEGG | 34 | 0.34 |
| sodium ion transport (GO:0006814) | 0.03050436 | 34 | GO | 3 | 0.03 | Estrogen signaling pathway | 0.00905157 | 45 | KEGG | 37 | 0.37 |
| protein phosphorylation (GO:0006468) | 0.0000235 | 159 | GO | 4 | 0.04 | Regulation of actin cytoskeleton | 0.00648235 | 67 | KEGG | 40 | 0.4 |
| supramolecular fiber organization (GO:009743 | 0.00036512 | 115 | GO | 4 | 0.04 | Gastric acid secretion | 0.03113544 | 26 | KEGG | 40 | 0.4 |
| external encapsulating structure organization (( | 0.00955794 | 72 | GO | 5 | 0.05 | Arrhythmogenic right ventricular cardiomyopa | 0.0007013 | 32 | KEGG | 41 | 0.41 |
| sodium ion transmembrane transport (GO:003 | 0.01120729 | 35 | GO | 5 | 0.05 | Sphingolipid signaling pathway | 0.02492299 | 38 | KEGG | 41 | 0.41 |
| extracellular structure organization (GO:00430 | 0.01186888 | 71 | GO | 5 | 0.05 | Axon guidance | 0.00023372 | 64 | KEGG | 42 | 0.42 |
| protein autophosphorylation (GO:0046777) | 0.03157435 | 53 | GO | 7 | 0.07 | Glioma | 0.04559065 | 25 | KEGG | 43 | 0.43 |
| axonogenesis (GO:0007409) | 0.000000317 | 95 | GO | 8 | 0.08 | cAMP signaling pathway | 0.0001049 | 75 | KEGG | 44 | 0.44 |

| | | | | | |
|---|---|---|---|---|---|
| extracellular matrix organization (GO:003019[illegible] | 0.0000818 | 104 | GO | 8 | 0.08 |
| positive regulation of cell motility (GO:20001[illegible] | 0.02766244 | 70 | GO | 9 | 0.09 |
| regulation of cell migration (GO:0030334) | 0.01120729 | 121 | GO | 11 | 0.11 |
| axon guidance (GO:0007411) | 0.0000000266 | 87 | GO | 12 | 0.12 |
| cation transport (GO:0006812) | 0.00677366 | 61 | GO | 15 | 0.15 |
| potassium ion transmembrane transport (GO:0[illegible] | 0.02782488 | 48 | GO | 16 | 0.16 |
| transmembrane receptor protein tyrosine kinas[illegible] | 0.01120729 | 120 | GO | 18 | 0.18 |
| cytoskeleton organization (GO:0007010) | 0.01613228 | 44 | GO | 22 | 0.22 |
| enzyme linked receptor protein signaling pathv | 0.00141912 | 54 | GO | 24 | 0.24 |
| inorganic cation transmembrane transport (GO | 0.00012075 | 96 | GO | 70 | 0.7 |

| | | | | | |
|---|---|---|---|---|---|
| Dopaminergic synapse | 0.00011241 | 51 | KEGG | 44 | 0.44 |
| AMPK signaling pathway | 0.00057292 | 45 | KEGG | 44 | 0.44 |
| Pathways in cancer | 0.01233083 | 142 | KEGG | 45 | 0.45 |
| Focal adhesion | 0.000018 | 74 | KEGG | 47 | 0.47 |
| Growth hormone synthesis, secretion and actio | 0.00303872 | 42 | KEGG | 49 | 0.49 |
| Glutamatergic synapse | 0.000000681 | 53 | KEGG | 50 | 0.5 |
| Phospholipase D signaling pathway | 0.00016791 | 55 | KEGG | 51 | 0.51 |
| mTOR signaling pathway | 0.00045666 | 55 | KEGG | 51 | 0.51 |
| Human papillomavirus infection | 0.04191574 | 89 | KEGG | 51 | 0.51 |
| Synaptic vesicle cycle | 0.00183459 | 31 | KEGG | 52 | 0.52 |
| Hippo signaling pathway | 0.04130516 | 48 | KEGG | 53 | 0.53 |
| Thyroid hormone synthesis | 0.00196217 | 30 | KEGG | 54 | 0.54 |
| Apelin signaling pathway | 0.02228549 | 43 | KEGG | 56 | 0.56 |
| Circadian entrainment | 0.0000799 | 41 | KEGG | 58 | 0.58 |
| Ras signaling pathway | 0.00032636 | 77 | KEGG | 59 | 0.59 |
| Insulin secretion | 0.0001162 | 37 | KEGG | 67 | 0.67 |
| Cortisol synthesis and secretion | 0.00924809 | 25 | KEGG | 67 | 0.67 |
| GnRH signaling pathway | 0.01520301 | 32 | KEGG | 68 | 0.68 |
| MAPK signaling pathway | 0.0000118 | 101 | KEGG | 73 | 0.73 |
| Parathyroid hormone synthesis, secretion and a | 0.0000757 | 44 | KEGG | 76 | 0.76 |
| Melanogenesis | 0.0059744 | 36 | KEGG | 79 | 0.79 |
| Rap1 signaling pathway | 0.0000102 | 78 | KEGG | 83 | 0.83 |
| Hypertrophic cardiomyopathy | 0.00993434 | 32 | KEGG | 88 | 0.88 |
| Cholinergic synapse | 0.00000236 | 51 | KEGG | 93 | 0.93 |
| Vascular smooth muscle contraction | 0.00885931 | 44 | KEGG | 98 | 0.98 |
| Cushing syndrome | 0.00829775 | 50 | KEGG | 99 | 0.99 |
| Calcium signaling pathway | 0.00022441 | 80 | KEGG | 100 | 1 |
| Inflammatory mediator regulation of TRP chan | 0.01138742 | 34 | KEGG | 102 | 1.02 |
| Dilated cardiomyopathy | 0.00211419 | 36 | KEGG | 105 | 1.05 |
| Aldosterone synthesis and secretion | 0.0000192 | 43 | KEGG | 111 | 1.11 |
| Oxytocin signaling pathway | 0.0000192 | 60 | KEGG | 146 | 1.46 |
| Adrenergic signaling in cardiomyocytes | 0.0000757 | 57 | KEGG | 156 | 1.56 |

**Table S11** Molecular weigth estimation form SEC-UV-MALS-RI analysis of (AATGG)n sequences

Column descriptions:

Injection: Each sample was injected on the SEC system and fractionated by molecular weight

Sample: The composition of the DNA in the injection

Peak: DNA samples that form conformations of different sizes are eluted as multiple peaks

Concentration Source: Data used to determine the concentration of each peak with MALS data to calculate molecular weights

Complex: Proposed size of the complex

| Injection | Sample | Peak | Concentration Source | Complex | # RT | # mw_kDa | # error_percent | error_kDa | # N | # Nerror | Concentration |
|---|---|---|---|---|---|---|---|---|---|---|---|
| 1 | (AATGG)2 | Peak 1 | Refractive index | Monomer | 36.058 | 2288 | 22.236 | 509 | 1 | 0.44472 | Refractive inde |
| 2 | (AATGG)3 | Peak 1 | Refractive index | Monomer | 34.566 | 2025 | 31.139 | 631 | 1 | 0.62278 | Refractive ind |
| 2 | (AATGG)3 | Peak 2 | Refractive index | Dimer | 31.5965 | 4090 | 27.952 | 1143 | 2.01975309 | 1.1934923 | Refractive inde |
| 3 | (AATGG)4 | Peak 1 | Refractive index | Monomer | 33.387 | 8427 | 9.184 | 774 | 1 | 0.18368 | Refractive ind |
| 4 | (AATGG)5 | Peak 1 | Refractive index | Monomer | 32.3545 | 10310 | 12.122 | 1250 | 1 | 0.24244 | Refractive inde |
| 4 | (AATGG)5 | Peak 2 | Refractive index | Dimer | 28.292 | 16790 | 17.134 | 2877 | 1.628516 | 0.47643864 | Refractive inde |
| 5 | (AATGG)6 | Peak 1 | Refractive index | Monomer | 31.328 | 10870 | 7.309 | 794 | 1 | 0.14618 | Refractive inde |
| 1 | (AATGG)2 | Peak 1 | UV | Monomer | 36.058 | 3422 | 22.236 | 761 | 1 | 0.44472 | UV |
| 2 | (AATGG)3 | Peak 1 | UV | Monomer | 34.8985 | 5590 | 21.104 | 1180 | 1 | 0.42208 | UV |
| 2 | (AATGG)3 | Peak 2 | UV | Dimer | 31.5965 | 9083 | 11.239 | 1021 | 1.62486583 | 0.52553036 | UV |
| 3 | (AATGG)4 | Peak 1 | UV | Monomer | 33.2645 | 7591 | 15.746 | 1195 | 1 | 0.31492 | UV |
| 4 | (AATGG)5 | Peak 1 | UV | Monomer | 32.3545 | 10005 | 10.183 | 1019 | 1 | 0.20366 | UV |
| 4 | (AATGG)5 | Peak 2 | UV | Dimer | 28.292 | 23640 | 16.8 | 3972 | 2.36281859 | 0.63755934 | UV |
| 5 | (AATGG)6 | Peak 1 | UV | Monomer | 31.319 | 14930 | 5.7 | 851 | 1 | 0.114 | UV |
